# Polyclonal antibody responses to HIV Env immunogens resolved using cryoEM

**DOI:** 10.1101/2021.01.28.428677

**Authors:** Aleksandar Antanasijevic, Leigh M. Sewall, Christopher A. Cottrell, Diane G. Carnathan, Luis E. Jimenez, Julia T. Ngo, Jennifer B. Silverman, Bettina Groschel, Erik Georgeson, Jinal Bhiman, Raiza Bastidas, Celia LaBranche, Joel D. Allen, Jeffrey Copps, Hailee R. Perrett, Kimmo Rantalainen, Fabien Cannac, Yuhe R. Yang, Alba Torrents de la Peña, Rebeca Froes Rocha, Zachary T. Berndsen, David Baker, Neil P. King, Rogier W. Sanders, John P. Moore, Shane Crotty, Max Crispin, David C. Montefiori, Dennis R. Burton, William R. Schief, Guido Silvestri, Andrew B. Ward

## Abstract

**In Brief:** Herein, we evaluated the immunogenicity of several BG505 SOSIP-based HIV Env immunogens in the rhesus macaque animal model using a combination of serology and biophysical approaches. We applied electron cryo-microscopy for high-resolution mapping of elicited polyclonal antibody responses, which provided detailed insights into the binding modes of the most common classes of antibodies elicited by BG505 SOSIP immunogens as well as the critical differences in immunogenicity that can occur as a consequence of engineered stabilizing mutations and partial glycan occupancy at different sites.

**Summary:** Engineered ectodomain trimer immunogens based on BG505 envelope glycoprotein are widely utilized as components of HIV vaccine development platforms. In this study, we used rhesus macaques to evaluate the immunogenicity of several stabilized BG505 SOSIP constructs both as free trimers and presented on a nanoparticle. We applied a cryoEM-based method for high-resolution mapping of polyclonal antibody responses elicited in immunized animals (cryoEMPEM). Mutational analysis coupled with neutralization assays were used to probe the neutralization potential at each epitope. We demonstrate that cryoEMPEM data can be used for rapid, high-resolution analysis of polyclonal antibody responses without the need for monoclonal antibody isolation. This approach allowed to resolve structurally distinct classes of antibodies that bind overlapping sites. In addition to comprehensive mapping of commonly targeted neutralizing and non-neutralizing epitopes in BG505 SOSIP immunogens, our analysis revealed that epitopes comprising engineered stabilizing mutations and of partially occupied glycosylation sites can be immunogenic.

**Graphical abstract:** 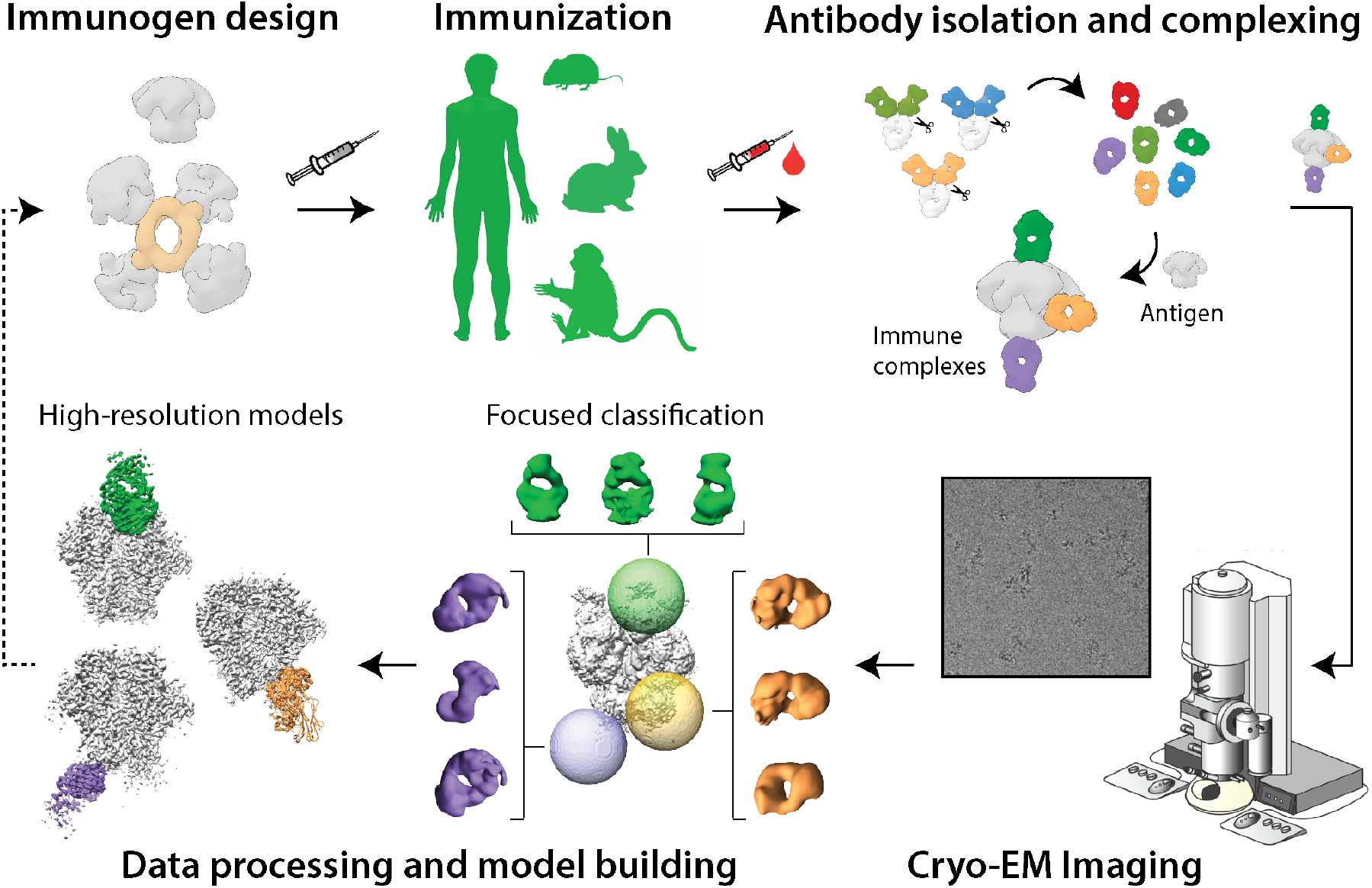

## Introduction

Envelope glycoprotein (Env) trimers derived from the BG505 genotype are the basis of many current HIV vaccine development efforts (Sanders et al., 2013, Whitaker et al., 2019, Medina-Ramirez et al., 2017, Steichen et al., 2019, Ringe et al., 2019, He et al., 2018, Dey et al., 2018, Tokatlian et al., 2018, Cirelli et al., 2019, Pauthner et al., 2017, Sharma et al., 2015, Escolano et al., 2019). When expressed as ectodomain constructs stabilized with SOSIP mutations, trimers based on this clade A sequence can be readily produced at high yields while preserving the nativelike, pre-fusion conformation targeted by known broadly neutralizing antibodies (bnAbs) (Sanders et al., 2013). Importantly, stabilization also reduces the exposure of epitopes for non-neutralizing antibodies (non-nAbs). BG505 SOSIP has been subject to many immunogenicity studies and is under evaluation in human clinical trials (ClinicalTrials.gov Identifiers: NCT03699241, NCT04177355 and NCT04224701).

Since its discovery, efforts have been made to improve the original BG505 SOSIP design by incorporating additional mutations. These aim to maximize the *in vitro* and *in vivo* stability of SOSIP trimers and increase expression levels (Torrents de la Pena et al., 2017, Steichen et al., 2016, Ringe et al., 2017, Chuang et al., 2017). These constructs have also been endowed with germline-targeting mutations, resulting in extensive engineering at the sequence level (Medina-Ramirez et al., 2017, Steichen et al., 2019, Steichen et al., 2016). In parallel, various nanoparticle platforms have been developed for trimer presentation to enhance *in vivo* trafficking properties and interaction with B-cells (Brouwer et al., 2019, Georgiev et al., 2018, Antanasijevic et al., 2020, Ueda et al., 2020, Tokatlian et al., 2018, Ringe et al., 2020, He et al., 2016, Moyer et al., 2020, Martin et al., 2020).

While BG505 SOSIP trimers have consistently elicited autologous neutralizing antibody responses in rabbit and macaque animal models (Pauthner et al., 2017, Klasse et al., 2018, Sanders et al., 2015, Torrents de la Pena et al., 2017), that were shown to be protective in macaques (Pauthner et al., 2019), they have failed to induce broadly neutralizing responses (Klasse et al., 2020). This is partly due to limited accessibility and immunoquiescence of bnAb epitopes, caused by extensive glycan shielding and sequestration of functionally essential protein domains in the quaternary structure (Burton and Hangartner, 2016). Conversely, the antibody response is redirected towards the readily accessible, and, in some cases immunodominant, epitopes that are typically strain-specific. Comprehensive mapping of such “off-target” epitopes in BG505 SOSIP and structural characterization of elicited antibodies will provide essential information for engineering the next generation of BG505-based immunogens with enhanced on-target reactivity. The standard approach would be to apply B-cell sorting to isolate representative monoclonal antibodies (mAbs) from different polyclonal families and perform structural characterization of each mAb. This approach, while valuable, is laborious in nature and impractical for routine large-scale application.

Herein, we introduce cryoEM-based polyclonal epitope mapping (cryoEMPEM): a method for rapid, high-resolution structural characterization of antibody-antigen complexes without the need for mAb isolation. We applied cryoEMPEM in combination with ELISA and pseudovirus inhibition assays to characterize the polyclonal antibody (pAb) responses elicited by BG505 SOSIP immunogens in rhesus macaques. The main goal was to acquire detailed, molecular-level understanding of the immunogenic landscape of stabilized BG505 SOSIP constructs as soluble trimers and presented on a nanoparticle. 21 high-resolution maps of trimer immune complexes with polyclonal Fabs provided detailed insights to the nature of antibody responses at 8 unique epitope clusters predominantly targeted in BG505 SOSIP immunogens.

## Results

### Immunogenicity of stabilized BG505 SOSIP trimer immunogens

Two BG505 SOSIP trimer antigens bearing different sets of stabilizing mutations, BG505 SOSIP MD39 and BG505 SOSIP.v5.2 N241/N289, were evaluated in this study (Table S1) (Torrents de la Pena et al., 2017, Steichen et al., 2016). The BG505 SOSIP.v5.2 N241/N289 construct included glycosylation sites at positions N241 and N289 to reduce access to the immunodominant glycan hole that exists in BG505 but is absent in the majority of HIV strains (McCoy et al., 2016, Klasse et al., 2018).

Two groups of six rhesus macaques were injected with 100 μg of BG505 SOSIP MD39 (Grp 1) or BG505 SOSIP.v5.2 N241/N289 (Grp 2) antigens at four different time points (Figure 1A). Each animal received the same antigen for all four immunizations. Serum/plasma samples were collected at 2-week intervals to monitor the development of antigen-specific antibodies by ELISA and neutralizing antibodies using the TZM-bl pseudovirus inhibition assay. All animals elicited antigen-specific responses that were within ~1 order of magnitude for animals in each group at the corresponding time points (Figure 1B, Table S2). Grp 1 animals had lower average binding titers compared to Grp 2 after the first two immunizations, but this difference ceased after the third and fourth immunizations.

**Figure 1.**
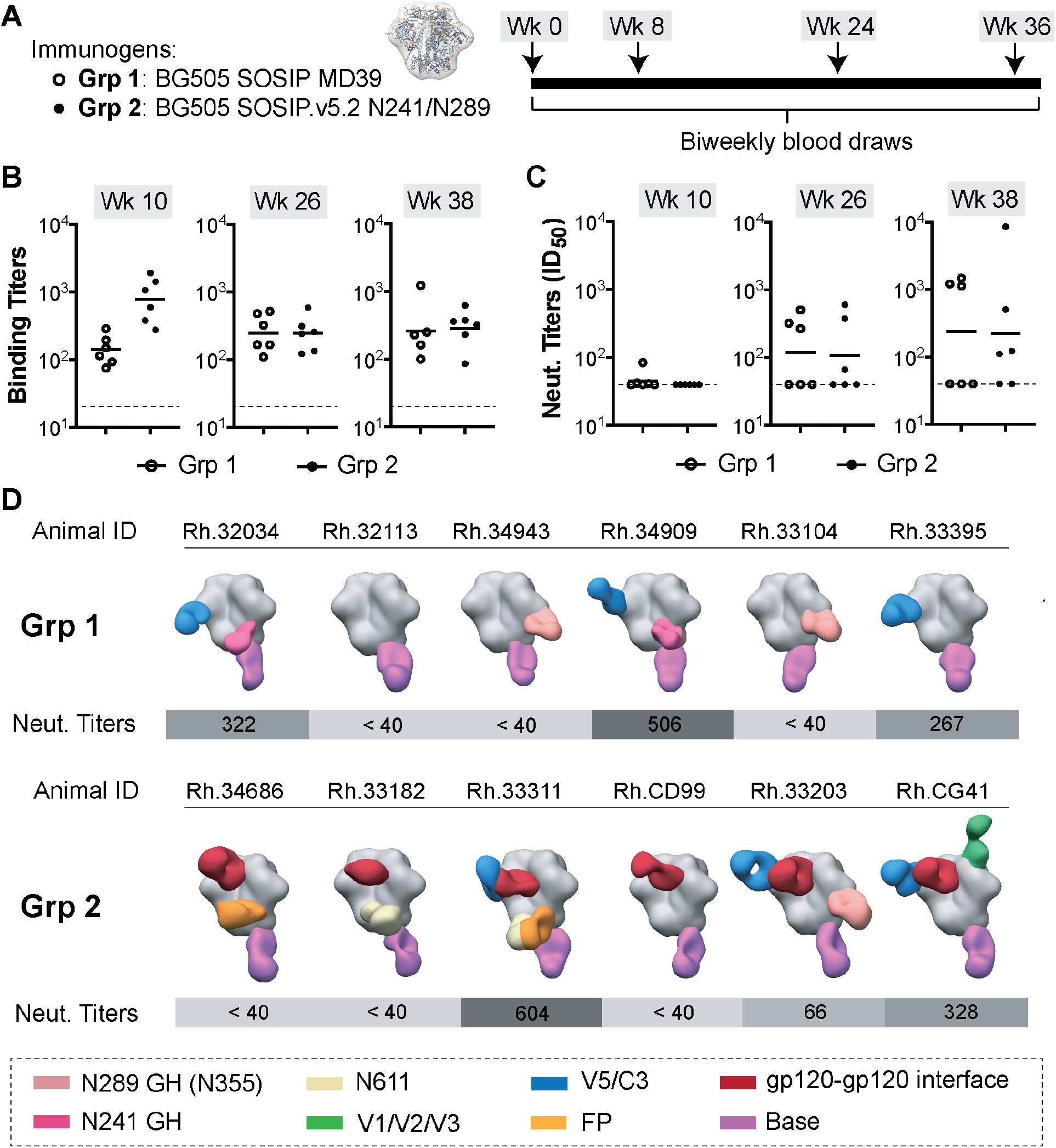
**[A]** Immunogen (left) and immunization schedule (right). **[B]** ELISA binding titers (midpoint) and **[C]** neutralizing antibody titers (ID_50_) for plasma samples collected at weeks 10, 26 and 38 (time points indicated above each graph; open circles – Grp 1; closed circles – Grp 2). Horizontal lines represent the geometric mean values at each time point. **[D]** Composite figures from nsEMPEM analysis of polyclonal responses at week 26. Animal IDs and neutralizing antibody titers (at week 26) for corresponding animals are shown above and below each composite figure, respectively. Color-coding scheme for antibodies targeting different epitope clusters is shown at the bottom. BG505 SOSIP antigen is represented in gray.

Neutralizing antibodies (NAbs) at serum neutralization titers (ID_50_) above 40, were first observed after the second immunization (Figure 1C, Table S3). NAb titers showed substantial variability between animals; ID_50_ values ranged from ~10^1^ – 10^4^ at week 38. Three animals in Grp 1 and two animals in Grp 2 failed to develop detectable levels of NAbs after four immunizations. Altogether, through ELISA and pseudovirus inhibition assays, we detected consistent elicitation of antigen-specific antibodies in all animals, while the levels of NAbs varied significantly between animals.

We applied negative stain EM-based polyclonal epitope mapping (nsEMPEM) to identify the epitope specificities of pAbs elicited by animals in the two groups (Bianchi et al., 2018, Nogal et al., 2020a). Immune complexes were prepared for imaging using either BG505 SOSIP MD39 or BG505 SOSIP.v5.2 N241/N289 with polyclonal Fabs obtained from plasma samples taken at Week 26. The nsEMPEM analysis revealed that all animals developed antibody responses against the base of the trimer, a highly immunodominant epitope in SOSIP constructs (Figures 1D and S1) (Cottrell et al., 2020, Havenar-Daughton et al., 2017, Hu et al., 2015). A C3/V5 response was present in three out of six animals in both groups, and it correlated well with autologous neutralization. Differences were also observed when comparing the vaccine-elicited antibody responses elicited in the two groups of animals. N241/N289 glycan hole responses were more prevalent in animals immunized with BG505 SOSIP MD39 (four animals in Grp 1 had this response but only one in Grp 2), suggesting that the presence of N241 and N289 glycans in the BG505 SOSIP.v5.2 N241/N289 immunogen suppressed the antibody response against this epitope. Overall, antibody responses against more diverse epitopes (e.g., N611 glycan epitope, fusion peptide, gp120-gp120 interface and V1/V2/V3) were detected in animals from Grp 2 than in Grp 1. Antibodies targeting the gp120-gp120 interface, a new epitope, were present in all six animals from Grp 2 but absent in Grp 1 animals.

### CryoEMPEM analysis of the BG505-specific polyclonal antibody responses

Based on the nsEMPEM analysis, we selected polyclonal Fabs isolated from animals Rh.32034 (Grp 1), Rh.33104 (Grp 1) and Rh.33311 (Grp 2) for cryoEM studies. Antibody specificities detected in these three animals represented all unique epitope clusters (Figure 1) except the V1/V2/V3 epitope, which was targeted only in animal Rh.CG41. Antibodies to this epitope were also detected in animals immunized with the nanoparticle immunogen (Grp 3, see below), and cryoEM characterization was instead performed on the samples from Grp 3.

CryoEM grids were prepared and imaged as described in the Methods section; data collection statistics are shown in Table S4. The data processing pipeline for cryoEMPEM is described in the Methods section and illustrated in Figure S2. From the three immune-complex samples, we reconstructed 16 high-resolution maps of structurally unique polyclonal antibody classes (pAbC) bound to the corresponding BG505 SOSIP antigen used in the immunization (Figure 2). Relatively large initial datasets (0.7–1.6 million clean particles after the symmetry-expansion step) enabled discrimination of Fabs bound to overlapping epitopes. This was particularly the case with base-targeting antibodies as they constitute the most prevalent type of antigen-specific antibodies (Figure 2, purple). Multiple pAbC classes targeting the N241/N289 glycan hole were also computationally isolated in the sample from animal Rh.33104, as well as C3/V5- and gp120-gp120 interface-directed antibodies in the sample from animal Rh.33311 (Figure 2).

**Figure 2.**
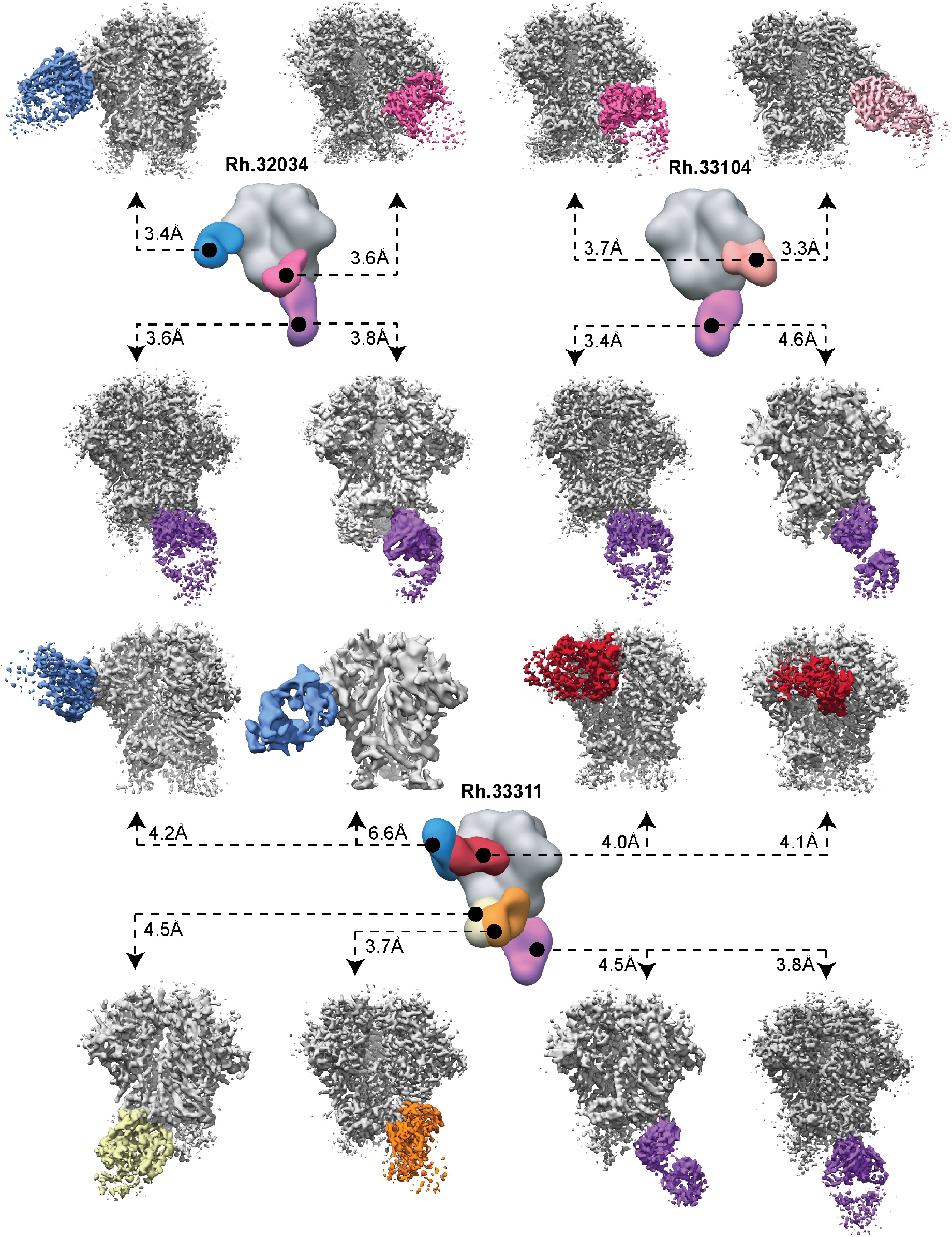
CryoEMPEM analysis of immune complexes generated using polyclonal Fabs isolated from plasma samples from animals Rh.32034 (top left), Rh.33104 (top right), and Rh.33311 (bottom). High resolution trimer-Fab complexes featuring structurally unique antibody specificities detected in cryoEM datasets are shown in the corresponding panels. BG505 SOSIP antigen is represented in gray and Fab densities are colored according to the scheme used in Fig 1. In the center of each panel is a composite figure from nsEMPEM. The apparent resolution of each reconstructed cryoEM map is indicated.

The apparent resolutions of reconstructed maps ranged from 3.3Å to 6.6Å with an average of ~4.0Å (Figure 2 and Figure S3). In most cases, the part of the map corresponding to the trimer was resolved to higher resolution than the Fab, as evident from the local resolution plots displayed in Figure S4. This is likely a consequence of the compositional and conformational heterogeneity stemming from the polyclonal nature of bound antibodies. Nevertheless, epitope-paratope interfaces were well-resolved in all EM maps, allowing for identification of amino acids comprising the epitope for each pAb.

Atomic models of trimer-Fab immune complexes were relaxed into the reconstructed EM maps with apparent resolution ≤ 4.6Å (15/16 maps met this criterion). Each Fab was represented as a poly-alanine backbone and comparison to published rhesus macaque Fab structures (Tran et al., 2014, Gohain et al., 2015, Navis et al., 2014) was used to assign the heavy and light chains. We applied these models and heavy/light assignments to determine the role of specific complementarity determining regions (CDR) and framework regions (FR) in antigen recognition. Structure refinement information can be found in the Methods section and the relevant statistics are shown in Table S5. Model-to-map fit for each complex structure is shown in Figure S5.

For clarity, the analysis of pAbC structures is performed on a per-epitope basis.

### Antibodies targeting the C3/V5 epitope

As described above, a correlation was observed between the elicitation of C3/V5-directed antibody responses and autologous neutralization (Figure 1). CryoEMPEM analysis of polyclonal samples isolated from macaque Rh.32034 and Rh.33311 yielded three maps with pAbs targeting this site (Figure 2, Figure S5A). Analysis of the reconstructed models for Rh.32034 pAbC-1 and Rh.33311 pAbC-5 revealed that these two polyclonal antibodies targeted largely overlapping epitopes at the interface of C3 and V5 regions (residues 354-358 and 459-466, respectively) flanked by glycans at positions N355 and N462 (Figure 3A). This epitope is not well-protected by the glycan shield and can be accessed from different angles. Indeed, the two reconstructed pAbCs displayed significantly different angles of approach and binding modes. In the case of Rh.33311 pAbC-5, the interaction is driven primarily by the heavy chain, while Rh.32034 pAbC-1 binds using both heavy and light chain. Notably, these are the first high resolution structures of Env trimer complexes with antibodies targeting this epitope.

**Figure 3.**
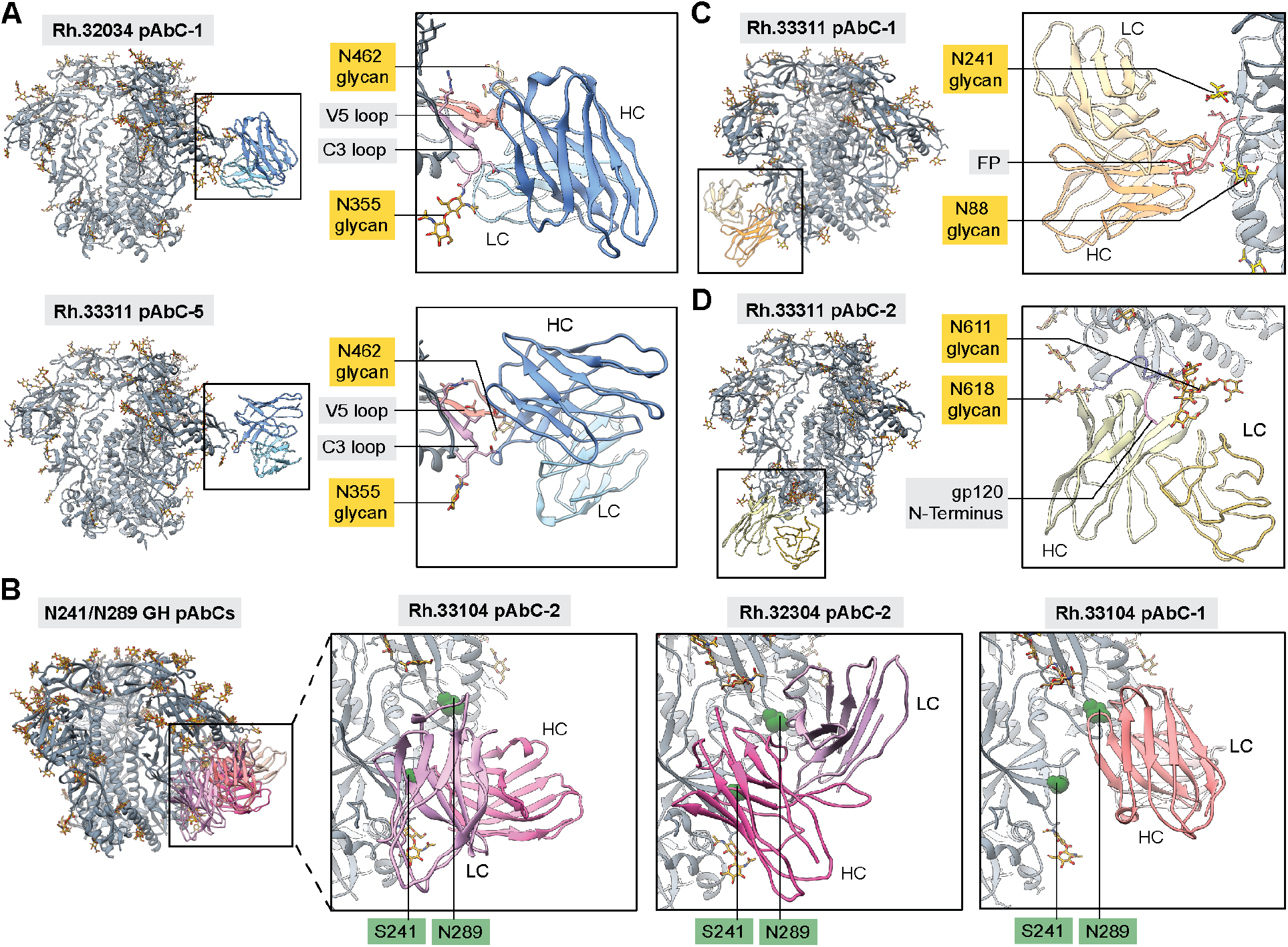
Structures of BG505 SOSIP antigens in complex with polyclonal antibodies targeting C3/V5 **[A]**, N241/N289 glycan hole **[B]**, fusion peptide **[C]**, and N611 glycan epitopes **[D]**. BG505 SOSIP trimers are shown in gray with N-linked glycans in golden yellow. Antibodies are colored using the same scheme as in Figures 1 and 2. Inferred heavy and light chains for each antibody are represented in different shades and labeled in each panel (HC and LC, respectively). The most relevant epitope/paratope components are indicated in the supporting panels. Ribbon and stick representations are used throughout the figure for display of structural components. In panel b), sphere representations are used for residues S241 and N289 to improve visibility.

The neutralization potential of the C3/V5 epitope in BG505 has been reported previously (Lei et al., 2019, Klasse et al., 2018, Nogal et al., 2020a, Zhao et al., 2020, Cirelli et al., 2019, Pauthner et al., 2017). Antibodies bound to this site can sterically block CD4 binding, which represents a potential mechanism of virus neutralization. However, the V5 loop is highly variable in terms of length, amino-acid sequence, and the number and location of N-linked glycosylation sites (Cao et al., 2017, Wang et al., 2013, Travers, 2012). Introduction of a glycan at position 465 (T465N mutation) in BG505 Env-pseudotyped virus results in a strong reduction of serum neutralization titers (ID_50_) for samples collected from both groups of animals at the week 38 time point (Table S6). In the original BG505 sequence, this site is occupied by threonine and is directly interacting with the two abovementioned C3/V5-targeting pAbs. Consistently, in Rh.32034 and Rh.33311 samples the T465N mutation caused a decrease in NAb titers by ~35-fold and ~15-fold, respectively. Altogether, our findings further support that BG505 SOSIP-elicited antibodies targeting the C3/V5 epitope are the primary contributors to autologous neutralization in the rhesus macaque animal model.

### Antibodies targeting the N241/N289 glycan hole epitope

The N241/N289 glycan hole in BG505 is an immunodominant epitope for autologous BG505-specific antibodies (Klasse et al., 2018). Although absent from the BG505 sequence, the N241 and N289 potential N-linked glycosylation sites (PNGS) exist in 97% and 79% of known HIV-1 strains, respectively (McCoy et al., 2016, Yang et al., 2020, Klasse et al., 2018). The Grp 1 immunogen, BG505 SOSIP MD39, did not include glycosylation sites at positions 241 and 289, and four out of six animals developed antibodies targeting this epitope (Figure 1). N241 and N289 glycans were present in the Grp 2 immunogen, BG505 SOSIP.v5.2 N241/N289, which reduced antibody responses against this epitope. Interestingly, one animal in Grp 2 (Rh.33203) still developed antibodies targeting the N289 epitope. This is likely a consequence of incomplete glycosylation at the engineered N289 PNGS in BG505 SOSIP. Site-specific glycan analysis data reported here (Table S7) and elsewhere (Ringe et al., 2019, Antanasijevic et al., 2020) suggests that the N289 site can be up to ~40% unoccupied while the glycosylation efficacy at the N241 site is >90%.

CryoEMPEM analysis of polyclonal antibodies from Grp 1 animals Rh.32034 and Rh.33104 complexed with the BG505 SOSIP MD39 antigen yielded three high resolution maps of pAbCs targeting the N241/N289 glycan hole epitope cluster (Figure 3B, Figure S5B). In sample Rh.33104, two antibodies were computationally sorted, including an N241-directed antibody not previously detected using nsEMPEM. The reconstructed antibodies targeted partially overlapping epitopes but had distinct binding modes. Rh.33104 pAbC-2 was biased toward the N241 glycan hole (residues 83-86 in C1, and residues 229-232, 239-243, 267-269 in C2). Conversely, Rh.33104 pAbC-1, elicited in the same animal, is biased toward the N289 glycan hole (residues 266-270 and 289-290 in C2, as well as the C3/V4 interface and glycans N355 and N398). Rhesus macaque mAbs targeting similar sites on BG505 SOSIP.664 have been recently described and structurally characterized (Cottrell et al., 2020). The third polyclonal antibody, Rh.32034 pAbC-2, featured a different angle of approach compared to the other two and made contact with both, the N241 (residues: 83-85, 227-231, 240-243) and N289 (residues 289-290 and 267-269) regions of the glycan hole epitope.

Consistent with previous findings, our data suggest that the N241/N289 glycan hole epitope in BG505 is immunodominant due to poor glycan shielding. High-resolution analysis revealed that the elicited antibodies can utilize different angles of approach and different combinations of heavy and light chain CDRs to make contact with the exposed peptidic surface consisting of residues from C1, C2, C3, and V4 regions. Serum neutralization experiments performed with mutated BG505 pseudovirus showed that glycan knock-ins at either position N241 or N289 resulted in only minor reduction of ID_50_ titers for Grp 1 samples (Table S6). These results, as well as previous observations (Klasse et al., 2018), suggest that glycan hole-directed antibodies elicited in rhesus macaques do not contribute significantly to autologous neutralization of BG505 virus. In contrast, antibodies targeting this site have been found to be major contributors to polyclonal rabbit serum neutralization (Klasse et al., 2018, McCoy et al., 2016, Brouwer et al., 2019).

### Antibodies targeting the fusion peptide and N611-glycan epitopes

Antibodies against the fusion peptide and proximal N611-glycan epitope were detected in three out of six animals in Grp 2 at Week 26 (Figure 1D). In animal Rh.33311 both types of responses were elicited. Polyclonal complexes with a single pAbC targeting each site were computationally sorted from the cryoEMPEM data (Figure 2), and structural models were derived from each reconstructed map (Figure 3C,D; Figure S5C,D).

Rh.33311 pAbC-1 utilizes all 3 HCDR loops to capture the N-terminal part of the fusion peptide (residues 512-520). Additional contacts are made to the fusion peptide proximal region (FPPR, residues 532-536) and the 664-helix in HR2 (residues 648-655). The light chain contributes only minimally to antigen interaction. Glycans at positions N611, N241, and N88 are visible in the map, but they do not make substantial contact with the antibody.

The fusion peptide epitope is a well-explored target for bnAbs and is the focus of ongoing HIV vaccine design efforts (Kong et al., 2019, Ou et al., 2020, Xu et al., 2018). We compared the Rh.33311 pAbC-1 structure to the available bnAb structures (Figure S6A). When looking at the angle of approach, this pAbC showed the most similarity to DFPH-a.15, a vaccine-elicited fusion peptide antibody isolated from rhesus macaques (Kong et al., 2019). However, the usage of heavy and light chains in binding is dissimilar between the two antibodies and the FP conformation in the Rh.33311 pAbC-1 bound state is more similar to ACS202, another FP-directed bnAb (van Gils et al., 2016, Yuan et al., 2019).

The N611-glycan epitope is proximal to the FP (Cottrell et al., 2020). Importantly, this PNGS is often only partially glycosylated in BG505 SOSIP constructs (Derking et al., 2020, Antanasijevic et al., 2020), leading to elicitation of narrow-specificity antibodies that preferentially recognize BG505 antigens lacking the N611 glycan. Some vaccine-elicited antibodies (e.g., RM20E1) are capable of neutralizing BG505 pseudotyped virus with N611 glycan knock-out but not the wt BG505 pseudovirus (Cottrell et al., 2020). CryoEMPEM analysis of pAbs from animal Rh.33311 yielded one antibody binding to this site (Rh.33311 pAbC-2, Figure 3D). Despite somewhat lower map resolution (~4.5Å), the density corresponding to N611 glycan is clearly discernible in the binding site. In fact, the Rh.33311 pAbC-2 appears to be making extensive contact with the N611 glycan via the HCDR loops 1 and 3 and LCDR loop 2. The peptide part of the epitope consists of HR2 residues 611-618 as well as the N terminus of gp120 (residues 30-33).

Antibodies targeting the N611 or FP epitopes were observed in one third of animals immunized with BG505 SOSIP.v5.2 N241/N289 trimers, but combined analysis of pseudovirus inhibition data suggests that they provide only minor (if any) contribution to autologous neutralization. Knock-out of the N611 glycan in the BG505 pseudovirus (N611A mutation) resulted in a subtle increase in serum neutralization titers for animal Rh.33311 compared to the wt BG505 pseudovirus (Table S6). A similar trend is observed in the other two animals from Grp 2 with N611 - and/or FP-directed antibodies (Rh.34686 and Rh.33182). Further studies with isolated mAbs are required to better characterize the two pAb families observed in animal Rh.33311.

### Antibodies targeting the gp120-gp120 interface epitope

Gp120-gp120 interface antibodies were detected in all six animals from Grp 2 but in none of the animals from Grp 1. CryoEMPEM analysis of animal Rh.33311 yielded two structurally unique polyclonal classes of antibodies with partially overlapping footprints (Figure 4A,B and Figure S5E). Rh.33311 pAbC-3 uses both the heavy and light chain to make contact with the C1 loop (residues 58-71) and the N262 glycan. Antibody binding induces the folding of this typically disordered region of the C1 loop into a short helix (Figure 4A). On the other hand, Rh.33311 pAbC-4 primarily interacts with the V3 tip (residues 304-320) using the heavy chain (Figure 4B). Binding to this epitope requires avoiding the N197 glycan, achieved by a slight rotation (~9°) of gp120 relative to the adjacent protomer, breaking the trimer symmetry (Figure S6B). Interestingly, the LCDR 3 loop of Rh.33311 pAbC-4 makes contact with the CD4 binding site residues 429 and 430; CD4bs is a highly pursued epitope target for HIV vaccine design (Medina-Ramirez et al., 2017, LaBranche et al., 2018, Saunders et al., 2019, Jardine et al., 2015, Jardine et al., 2013).

**Figure 4.**
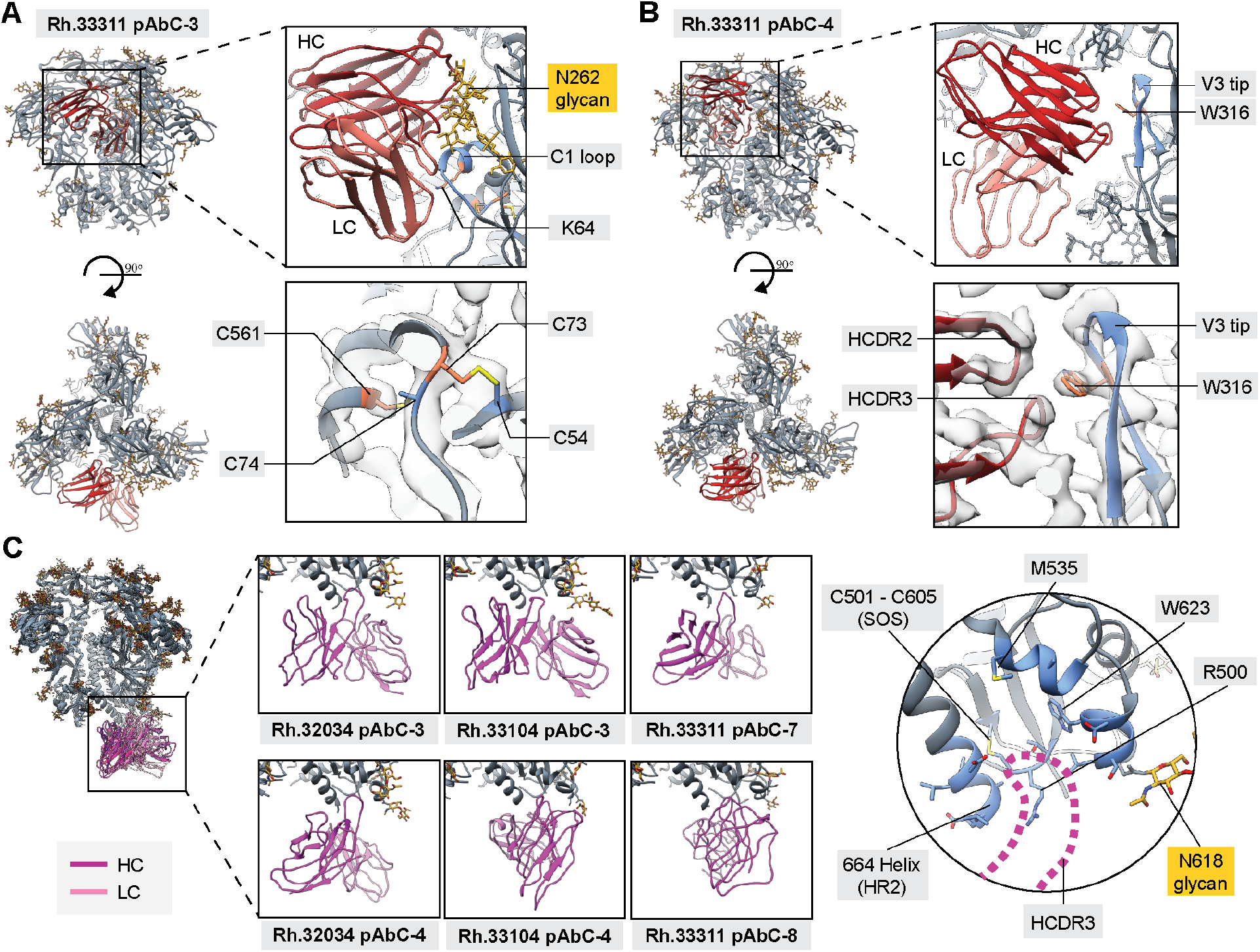
Structures of BG505 SOSIP antigens in complex with polyclonal antibodies targeting gp120-gp120 interface **[A,B]** and base epitopes **[C]**. BG505 SOSIP trimers are shown in gray with N-linked glycans in golden yellow. Antibodies are colored using the same scheme as in Figures 1 and 2. Inferred heavy and light chains for each antibody are represented in different colors and labeled in each panel (HC and LC, respectively). The most relevant epitope/paratope components are indicated in the panels. Trimer residues in direct contact with the antibodies are colored blue (ribbon and stick representation). Engineered stabilizing residues are represented as sticks and colored orange. EM Map is displayed as transparent light gray surfaces in panels **A** and **B.**

Interestingly, both antibodies bind to epitopes that have not previously been described. Closer examination, however, reveals that they make significant contacts with non-native stabilizing residues introduced during the immunogen design process. Rh.33311 pAbC-3 interacts with residue K64 (E in the original BG505 sequence; Figure 4A, top right panel). Additionally, in the immediate proximity to the binding site, engineered cysteine residues C73 and C561 appear to be cross-linking with the native disulfide bridge usually formed between residues C54 and C74 (Figure 4A, bottom right panel). In the high-resolution cryoEM maps of BG505 SOSIP.v5.2 N241/N289 constructs reported here, we have found evidence of an altered disulfide bond network at this site (i.e., the pairing of C73 with C54 and C74 with C561). However, the effect is stochastic in nature, and we have previously reconstructed EM models of BG505 SOSIP.v5.2 with the “intended” pairing of cysteines (Nogal et al., 2020b). The altered disulfide network together with the E64K mutation therefore comprises a neo-epitope not present on wt BG505 Env. Similarly, the Rh.33311 pAbC-4 binds the V3 tip, with the HCDR 2 and 3 contacting residue W316 (Figure 4B, top and bottom right panels). Tryptophan was introduced in place of alanine at this position beginning in SOSIP.v4 constructs to stabilize the trimers in a closed, pre-fusion state (de Taeye et al., 2015, Torrents de la Pena et al., 2017).

We next tested whether the interface-directed antibodies could bind to BG505 in the absence of these stabilizing mutations. Using BG505 SOSIP.v3 trimers that were not engineered with the abovementioned C1 and V3 mutations, we conducted nsEMPEM on the Rh.33311 polyclonal sample; experiments with BG505 SOSIP.v5.2 N241/N289 (the immunogen) were also performed in parallel for comparison (Figure S1B). We observed antibodies to C3/V5, N611, FP and base epitopes with either antigenic probe, suggesting that these epitopes are not significantly affected by the engineered residues. However, interface-specific antibodies were not detected with BG505 SOSIP.v3, indicating that their binding is dependent on the presence of stabilizing mutations. Consequently, these classes of antibodies are unable to recognize native Env trimers or neutralize virus.

### Antibodies targeting the trimer base epitope

The trimer base is the most immunodominant epitope cluster on BG505 SOSIP trimers (Cottrell et al., 2020, Havenar-Daughton et al., 2017, Hu et al., 2015). In native membrane-bound Env trimers, this part of the molecule is connected to the transmembrane domain and shielded from antibodies by the viral membrane (Nogal et al., 2020a). From the three cryoEMPEM datasets (Rh.32034, Rh.33104, Rh.33311), we recovered six high-resolution maps with base-targeting pAbCs (Figure 2, Figure S5E). Analysis of the reconstructed EM maps and modeled structures revealed that antibodies can approach the base at many different angles (Figure 4C, left). Surprisingly, all six pAbCs converge on a common epitope and share similar binding mechanisms. Base-specific pAbCs utilize HCDR3 to make contact with the binding pocket located at the C terminus of BG505 SOSIP (664-helix in HR2; Figure 4C, right). These findings are consistent with previous studies of the base-targeting mAbs RM20A3 and RM19R, isolated from rhesus macaques immunized with BG505 SOSIP.664 (Cottrell et al., 2020, Berndsen et al., 2020).

### Design and characterization of two-component nanoparticle system for presentation of BG505 SOSIP immunogens

To explore the effects of nanoparticle presentation of BG505 SOSIP trimers on the epitope specificities of vaccine-elicited antibodies, we used the computationally designed tetrahedral nanoparticle scaffold T33-31 (King et al., 2014, Bale et al., 2015). This nanoparticle consists of four copies each of two complementary trimeric components, referred to as T33-31A and T33-31B (Figure S6C). Both components have outward-facing N termini that were genetically fused to the C terminus of BG505 SOSIP.v5.2(7S) N241/N289, using a flexible GS linker (Figure 5A, Figure S6C). nsEM analysis of the BG505 SOSIP-T33-31A and -T33-31B antigen-bearing components confirmed that genetic fusion did not result in major issues with antigen folding and stability (Figure S6D). Additional density corresponding to the nanoparticle component is clearly discernible in 3D reconstructions of the two antigen-bearing components.

**Figure 5.**
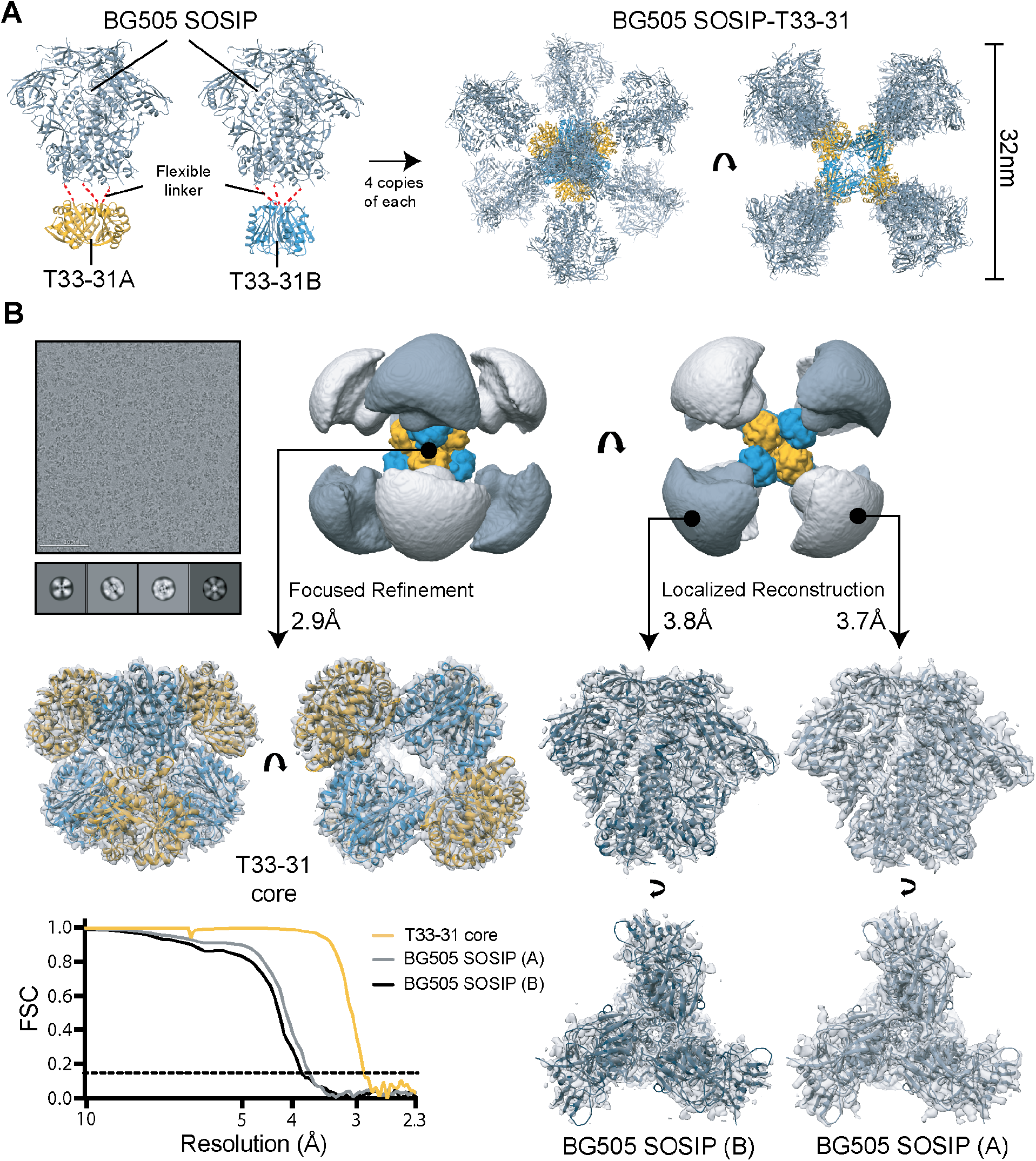
**[A]** Models of the two antigen-bearing components (left) and BG505 SOSIP-T33-31 nanoparticle (right). **[B]** CryoEM analysis of the designed BG505 SOSIP-T33-31 nanoparticle. Representative raw micrograph and sample 2D class-averages of imaged particles are shown in the top left corner. Low-pass filtered map of the entire nanoparticle is shown in the top right corner (orange – T33-31A; blue – T33-31B; light gray – BG505 SOSIP (A); dark gray – BG505 SOSIP (B)). Final maps and relaxed models from subparticle analysis are shown in the middle/bottom panels. Fourier Shell Correlation (FSC) curves for each subparticle map are presented in the bottom left corner.

The BG505 SOSIP-T33-31 nanoparticle was assembled by combining equimolar amounts of BG505 SOSIP-T33-31A and BG505 SOSIP-T33-31B. Analysis using cryoEM showed that nanoparticles assembled as expected with eight flexibly linked trimer antigens on their surface (Figure 5B, top row). We processed the molecular projection data by independently analyzing signal originating from three flexibly bound objects: T33-31 nanoparticle core, BG505 SOSIP fused to component A and BG505 SOSIP fused to component B (Figure 5B, middle and bottom rows). The T33-31 nanoparticle core structure closely matched the published crystal structure (PDB ID: 4zk7; (Bale et al., 2015)) with a backbone RMSD of 0.65Å, confirming that antigen attachment did not affect nanoparticle assembly. Additionally, the analysis of the two BG505 SOSIP structures (A and B) revealed that the presented trimer antigens are folded correctly in the native-like, pre-fusion state.

Consistent with the structural data, biolayer interferometry (BLI) experiments demonstrated that the antibody binding profiles of BG505 SOSIP fused to each nanoparticle were equivalent to those of free trimers (Figure S6E). However, reduced binding to antibodies targeting epitopes in the middle/lower part of the trimer such as the fusion peptide (PGT151, VRC34, ACS202), gp120-gp41 interface (35O22), gp41-gp41 interface (3BC315) and the trimer base (RM19R, RM20A3) was observed in the context of the assembled nanoparticles. This is likely caused by the crowding of trimer antigens on the nanoparticle surface, limiting the accessibility of epitopes near or at the trimer base; an observation consistent with previously reported data using other nanoparticle scaffolds (Antanasijevic et al., 2020, Ueda et al., 2020, Brouwer et al., 2019).

### Immunization experiments with BG505 SOSIP-T33-31 nanoparticle

After confirming the presence of appropriate antigenic and structural features in the designed BG505 SOSIP-T33-31 nanoparticle, we proceeded to use this reagent for rhesus macaque immunizations. We tested whether presentation of BG505 SOSIP on the T33-31 nanoparticles could have a detectable effect on antibody-mediated immune response compared to immunization experiments with soluble BG505 SOSIP trimers (Figure 6A; Grp 3 of animals). We used ELISA and TZM-bl pseudovirus inhibition assays to determine the relative proportions of vaccine-elicited antigen-specific and neutralizing antibodies, respectively (Figure 6B, Tables S2 and S3). ELISA midpoint titers were comparable to those obtained for Grps 1 and 2 at all time points, suggesting that antibody responses of similar magnitude were elicited against the BG505 SOSIP antigen. Serum neutralization titers were somewhat lower compared to Grps 1 and 2, with only two animals developing detectable neutralization titers (ID_50_ > 40) after all four immunizations.

**Figure 6.**
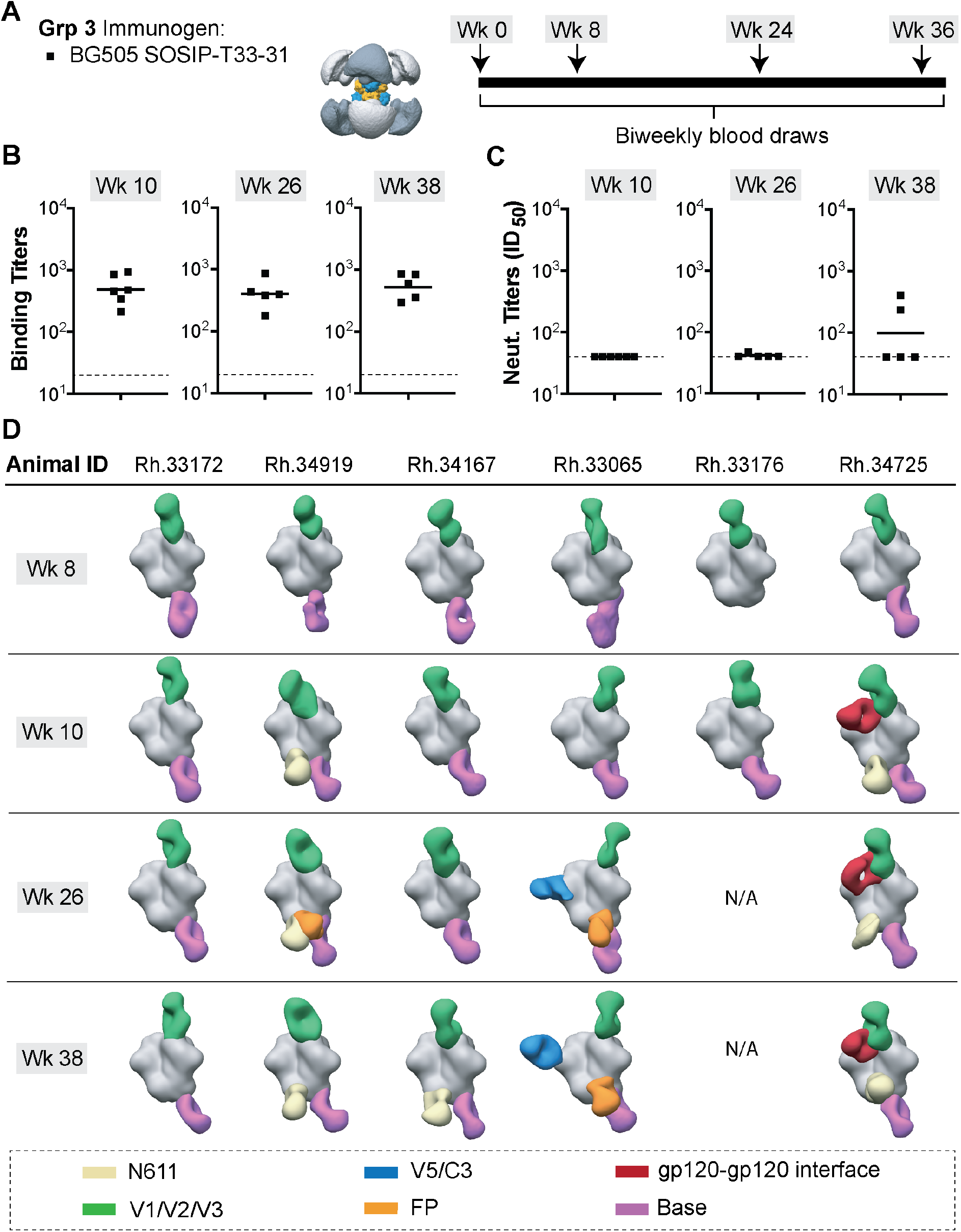
**[A]** Immunization information and schedule. **[B]** ELISA binding titers (midpoint) and **[C]** neutralizing antibody titers (ID_50_) for plasma samples collected at time points indicated above each graph. Horizontal lines represent the geometric mean values at each time point. **[C]** Composite figures from nsEMPEM analysis of polyclonal responses at weeks 8, 10, 26, and 38. Animal IDs are shown above each dataset. Color-coding scheme for antibodies targeting different epitope clusters is shown at the bottom. BG505 SOSIP antigen is represented in gray.

Next, we applied low-resolution polyclonal epitope mapping experiments to assess if nanoparticle presentation altered the epitope specificity/distribution of vaccine-elicited pAbs. The nsEMPEM analysis was performed on polyclonal samples from all Grp 3 animals at weeks 8, 10, 26, and 38 (Figure 6C). Similar to Grps 1 and 2, all animals developed antibodies targeting the base of the BG505 SOSIP trimer. In five of six animals, these antibodies were observable after the first immunogen dose. C3/V5-targeting antibodies were detected in animal Rh.33065 at weeks 26 and 38. Serum neutralization was also detected for this animal at those two time points. In contrast to the soluble trimer immunization experiments, knock-in of the N465 glycan site to BG505 Env did not significantly impact NAb titers for this animal (Table S6.), indicating that antibody responses to other (non-C3/V5) epitopes also contribute to neutralization. Finally, four of six animals developed antibodies to the FP and N611 epitopes after all four immunogen doses. This is similar to the nsEMPEM results from Grp 2 but different from Grp 1.

The greatest difference for Grp 3, compared to Grps 1 and 2, was observed with polyclonal antibodies directed to the V1/V2/V3 epitope cluster (Figure 6C). These antibodies were elicited by all animals after the first immunization, and they appear to target a range of epitopes comprising the V1, V2, and V3 variable loops at the trimer apex. Despite some similarity to apex-specific bnAbs (e.g., PGT121, PG9), the elicited antibodies appear to lack strong neutralizing activity (Table S3). However, animal Rh.33172 developed NAb titers (ID_50_ = 407) after four immunizations and nsEMPEM analysis revealed the presence of pAbs against two epitope clusters across all time points, namely the V1/V2/V3 and the base, suggesting that at least some of the V1/V2/V3-directed antibodies were neutralizing.

### CryoEMPEM analysis of polyclonal antibody responses in animal Rh.33172

Polyclonal Fabs from animal Rh.33172 (week 38) were complexed with BG505 SOSIP.v5.2(7S) N241/N289 (matched to the immunogen presented on T33-31 nanoparticle) and subjected to cryoEMPEM analysis. The analysis yielded five maps, each featuring a unique polyclonal antibody class (Figure 7A). Four pAb classes targeted partially overlapping but distinct V1/V2/V3 epitopes and one antibody class was directed against the trimer base. The average apparent resolution of the reconstructed EM maps was ~4.0Å (Figure 7A, Figure S8). Analysis of the maps and corresponding models revealed that the Rh.33172 pAbC-2 epitope consists mainly of V2 and V3 residues, while the other three antibodies primarily interact with the V1 loop, making minor additional contacts with V2 and V3 (Figure 7B).

**Figure 7.**
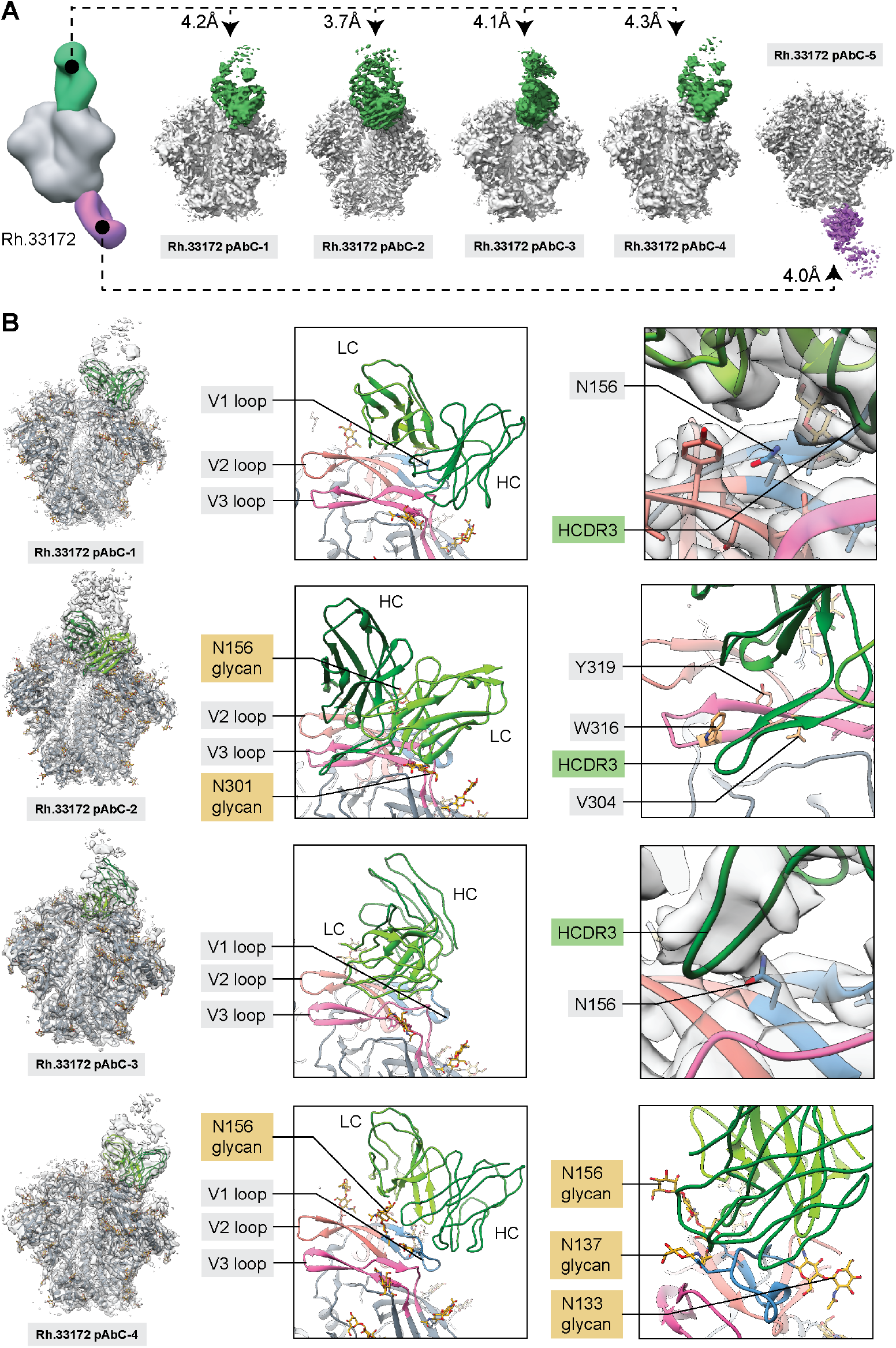
**[A]** CryoEMPEM analysis of the immune complexes generated using polyclonal Fabs isolated from animal Rh.33172 (week 38 plasma samples). BG505 SOSIP antigen is represented in gray and Fab densities are colored according to the scheme used in Fig 6. The apparent resolution of each reconstructed map is indicated. **[B]** Structures of four V1/V2/V3-targeting pAbs. Models (ribbon representation) and maps (light gray mesh) are shown on the left, and close-up views of each structure with the most relevant epitope/paratope components are indicated (V1 loop – blue, V2 loop – salmon, V3 loop – magenta). Inferred heavy and light chains for each antibody are labeled in each panel and represented in a dark and light green color, respectively.

Rh.33172 pAbC-1 and pAbC-3 approach the V1 loop from different angles, but both converge on the highly conserved N-linked glycosylation site N156 (Travers, 2012, Wang et al., 2013, Cao et al., 2017). Strikingly, close inspection of the data revealed no evidence of glycan density at this site in both maps (Figure 7B, left panels). Furthermore, for Rh.33172 pAbC-1, no glycan density was found at N137, a residue that is also within the binding site for this antibody. These results suggest that both classes of antibodies target a small subset of BG505 SOSIP antigens that lack N-linked glycans at one or both of the PNG sites. These pAbs share high similarity to rhesus macaque mAbs elicited with the BG505 SOSIP-based immunogen RC1, in which the conserved V1 glycans: N133, N137, and N156, were knocked out to improve accessibility at this epitope (Escolano et al., 2019). Site-specific glycan analysis data for the BG505 SOSIP.v5.2(7S) N241/N289 construct (Table S7) reveal that while N137 can be up to 50% unoccupied, the N156 PNG site is glycosylated in >89% of the protomers. Furthermore, the N156 site appears to be fully glycosylated in the BG505 SOSIP-T33-31A and -T33-31B components that make up the nanoparticle immunogen. Given the relatively small amount of glycan underoccupancy, it is surprising that antibodies are readily induced against this site.

We tested if knocking out the N156 glycosylation site on a BG505 Env would make the pseudovirus more susceptible to neutralization by sera from this group of animals. Four out of five animals, including Rh.33172, had increased serum neutralization titers (ID_50_) against the N156 mutant pseudovirus compared to the wt BG505 (Table S6). This finding supports our observations from the structural data.

Analysis of the Rh.33172 pAbC-2 structure revealed that this polyclonal antibody class utilizes a long, 22-amino-acid HCDR3 to interact with the V3-tip (residues 304-321). The antibody coordinates glycans N301 and N156 in BG505 and contacts the V2 loop (residues 170-173). This epitope is similar to the Rh.33311 interface-targeting pAbCs; it is partially constituted of engineered residues. In the BG505 SOSIP.v5.2(7S) N241/N289 construct there are three stabilizing mutations within the V3-loop (R304V, A316W, and A319Y), and Rh.33172 pAbC-2 makes contact with all of the mutated residues (Figure 7B, right). nsEMPEM analysis revealed that Rh.33172 pAbC-2 failed to bind to BG505 SOSIP.v3, which lacks the stabilizing mutations in V3 (Figure S1C, Rh.33172 pAbC-2 depicted in olive green). Consequently, this class of polyclonal antibodies are non-neutralizing because they are unable to recognize wt BG505 Env.

Finally, the Rh.33172 pAbC-4 makes extensive contacts with the V1 loop (residues 132 – 156) as well as the V2 loop (residues 186-190) while also coordinating the V1 glycans (N133, N137 and N156). This antibody class can interact with the BG505 SOSIP.v3 antigen (Figure S1C; depicted in light green) and is likely responsible for the serum neutralization observed for animal Rh.33172 at the week 38 time point. Previous studies have shown that V1-targeting antibodies in BG505 can be moderately neutralizing (Klasse et al., 2018, Zhao et al., 2020, Nogal et al., 2020a).

## Discussion

In this study we describe cryoEMPEM, a method for high-resolution mapping of vaccine-induced polyclonal antibody responses directly from sera. Isolation and analysis of mAbs is the gold standard for structural characterization of antibodies and antibody-antigen complexes. However, cryoEMPEM is a faster alternative offering a more comprehensive view of diverse polyclonal antibody classes targeting an antigen at the serum level. Under ideal conditions (i.e., access to state-of-the-art imaging and computational resources), cryoEMPEM analysis can be completed within ~10 days from serum/plasma collection. Therefore, it can be applied to study immunization progress in real time and make appropriate regimen changes, when needed.

Compared to structural characterization of mAbs, there are additional experimental challenges caused by the polyclonal nature of bound antibodies: compositional heterogeneity in the EM data and the lack of sequence information. Using the described focused classification approach, we reconstructed 21 maps of different polyclonal antibody classes with apparent resolution averaging ~4.1 Å, from the immune sera of four animals. The quality and resolution are comparable to cryoEM maps of immune complexes with mAbs. Additionally, high resolution analysis allowed us to resolve multiple structurally unique polyclonal antibody classes that interact with overlapping epitopes within the same polyclonal sample. The maps were of sufficient quality to relax pseudo-atomic models and while the explicit antibody sequences are inherently unknown, the models represented as poly-alanine peptide backbone provided insight into different aspects of antibody binding (i.e., all epitope contacts on the antigen side, and usage of heavy and light chain and different CDRs on the antibody side).

Our study included the comparison of three different BG505-based immunogens and revealed critical differences in the immune response elicited by each design. The three immunogens primarily differed in their content of stabilizing mutations and glycosylation sites. BG505 SOSIP MD39 trimers elicited N241/N289 glycan hole responses in the majority of animals in Grp 1. This highly specific and undesirable response is almost fully absent in animals receiving the other two immunogens (Grps 2 and 3) because of the engineered PNG sites at positions N241 and N289. We speculate that glycan masking of the immunodominant glycan hole epitope in these immunogens drove the development of antibodies against more diverse sites (e.g., FP, N611, V1/V2/V3 sites). Similar observations have been reported previously with suppression of immune response to the trimer base epitope and epitopes created by trimer degradation (Cirelli et al., 2019, Moyer et al., 2020). However, it is still unclear if immuno-masking of all immunodominant autologous epitopes in BG505 SOSIP would lead to elicitation of antibody responses targeting poorly accessible, cross-reactive sites.

The SOSIP.v5.2 stabilizing mutations in V3 (A316W) and C1 (E64K, A73C) in the Grp 2 and 3 immunogens generated stabilized neo-epitopes that were immunogenic. We demonstrated that antibodies targeting these sites are unable to interact with antigen in the absence of stabilizing mutations and consequently with the wt BG505 Env on a virion. They are therefore nonneutralizing. Besides the potential immunodominance of these neo-epitopes, they are also problematic because the antibodies targeting them could potentially sterically block access to bnAb epitopes (e.g., CD4bs and V2-apex).

While nanoparticle display did not improve the overall levels of neutralizing antibody responses, it had a dramatic effect on the epitopes targeted and angles of approach of elicited antibodies, particularly after early immunizations. This finding is consistent with previous studies performed with ConM and BG505-based nanoparticle immunogens (Antanasijevic et al., 2020, Brouwer et al., 2019, Brouwer and Sanders, 2019). While immuno-focusing on the trimer apex is a desirable feature, some of the antibody responses we observed targeted non-native glycan holes that were present on the recombinantly expressed antigen, even at very low levels. This result further emphasizes the need to conduct glycopeptide occupancy analysis of Env immunogens. Interestingly, our analyses revealed that presentation on the nanoparticle failed to eliminate base-directed antibody responses, suggesting that this epitope is still readily accessible on the nanoparticle or becomes accessible upon nanoparticle disassembly *in vivo.*

This work provides a comprehensive reference for HIV vaccine design efforts utilizing BG505 SOSIP immunogens. The presented data can be readily applied to engineer changes that would immuno-silence undesirable epitopes and/or immuno-focus to a specific site in this construct. More broadly, cryoEMPEM is a general method that can be applied to different subunit vaccine platforms for rapid and detailed mapping of antigenic landscapes at high resolution. With continued methodological advances, it may be possible to directly determine the sequences of bound polyclonal antibodies by cryoEMPEM.

## Materials and Methods

### DNA vectors and cloning

Constructs containing BG505-SOSIP.v5.2 N241/N289 and BG505-SOSIP.v5.2(7S) N241/N289 genes codon-optimized for mammalian cell expression were subcloned into a pPPI4 vector. BamHI and NheI restriction sites were used for insertion of genes encoding T33-31A and T33-31B nanoparticle assembly components to the C-terminus (residue 664) of BG505-SOSIP.v5.2(7S) N241/N289. 10-amino-acid flexible linkers (sequence: GSGSGSGSGG) were introduced between the SOSIP trimer and each nanoparticle component. Restriction enzymes and Quick Ligation kits were obtained from New England Biolabs (NEB). Sequences were verified by Sanger sequencing (Genewiz).

### Expression and purification of BG505-SOSIP and antigen-bearing components

BG505 SOSIP-based constructs were purified as described previously(Antanasijevic et al., 2020, Martin et al., 2020). Briefly, BG505 SOSIP.v5.2 N241/N289, BG505 SOSIP.v5.2(7S) N241/N289, BG505 SOSIP-T33-31A, BG505 SOSIP-T33-31B constructs were expressed in HEK293F cells (Invitrogen) and purified using PGT145 immunoaffinity chromatography. BG505 SOSIP MD39 was expressed in FreeStyle 293F cells (Thermo Fisher Scientific) and purified using a 2G12 immunoaffinity column. Samples were eluted from the corresponding immunoaffinity matrix with 3M MgCl2 buffer and subjected to size-exclusion chromatography (SEC) on a HiLoad® 16/600 Superdex® pg200 (GE Healthcare). Samples were subsequently concentrated to 1 mg/ml and stored frozen in DPBS (Thermo Fisher Scientific).

### BG505 SOSIP-T33-31 nanoparticle assembly and purification

The two antigen-bearing nanoparticle components (BG505 SOSIP-T33-31A and BG505 SOSIP-T33-31B) were independently purified using the protocol described above. ~2.5 mg of each component were combined and incubated for 72 hours at 37 °C, to drive co-assembly into nanoparticles. Average nanoparticle yield after 72 hours was ~70% of the starting material. Assembled nanoparticles were purified from residual unassembled components using Sephacryl S-500 HR column with DPBS (Thermo Fisher Scientific) as the running buffer. ToxinSensor™ Single Test Kit (GenScript) was applied to verify that the endotoxin levels of the nanoparticle sample were below 50 EU/kg per dose.

### Biolayer interferometry

The experiments were performed as described previously(Antanasijevic et al., 2020, Ozorowski et al., 2018). Kinetics buffer (DPBS + 0.1 % BSA + 0.02 % Tween-20) was used to prepare antibody and antigen dilutions. IgG versions of antibodies were diluted to 5 μg/ml. The concentrations of trimer and nanoparticle samples were adjusted to equimolar concentration (500 nM) of the BG505-SOSIP.v5.2(7S) N241/N289 antigen in each sample. BG505-SOSIP.v5.2(7S) N241/N289 trimer was included as a reference. Octet Red96 instrument (ForteBio) was used for data collection. Antibodies were immobilized onto antihuman IgG Fc capture (AHC) biosensors (ForteBio). The lengths of association and dissociation steps were adjusted to 180 s and 300 s, respectively. Octet System Data Analysis v9.0 (FortéBio) software package was used for data processing. The background was corrected by subtracting the negative control datasets (kinetics buffer). The baseline step immediately preceding the association step was used for alignment of y-axes. An interstep correction between the association and dissociation steps was also introduced. Final data plots were prepared in Excel. Experiments were performed in duplicate to test reproducibility, but only one measurement is presented.

### Site-specific glycan analysis using mass spectrometry

The experiments were performed with BG505 SOSIP-T33-31A and BG505 SOSIP-T33-31B antigen-bearing components as described previously (Antanasijevic et al., 2020, Behrens et al., 2016). A standard library for BG505 SOSIP.664 expressed in HEK293F cells was used to search the MS data. The relative amount of glycan at each site was determined by comparing the extracted chromatographic areas for different glycopeptides with an identical peptide sequence. For analysis, we applied precursor mass tolerance of 4 ppm and 10 ppm for fragments with a false discovery rate (FDR) of 1%. The relative amount of glycan at each site and the unoccupied proportion were determined by comparing the extracted ion chromatographic areas for different glycopeptides with an identical peptide sequence. Glycan analysis data for the underlying BG505 SOSIP.v5.2(7S) N241/N289 Env construct (as free trimer) used for the generation of antigen-bearing components has been published elsewhere (Antanasijevic et al., 2020). It is presented in this study as a reference for comparisons with BG505 SOSIP-T33-31A and BG505 SOSIP-T33-31B.

### Immunization experiments

Rhesus macaque immunizations and blood draws were performed at the Yerkes National Primate Research Center, Atlanta, GA, USA. All procedures were approved by Emory University Institutional Animal Care and Use Committee protocol 201700723, and animal care facilities are accredited by the U.S. Department of Agriculture and the Association for Assessment and Accreditation of Laboratory Animal Care International. Three groups of Indian rhesus macaques (6 animals per group) were immunized at weeks 0, 8, 24 and 36 with BG505 SOSIP MD39 (Grp 1, 100 μg per dose), BG505 SOSIP.v5.2 N241/N289 (Grp 2, 100 μg per dose) and BG505 SOSIP-T33-31 nanoparticle (Grp 3, 119 μg per dose). Total dose was normalized to administer an equivalent molar concentration of BG505 SOSIP antigen across all groups. Immunogens were formulated with Matrix-M™ (Novavax, Inc.; 75 μg per dose) for the first three immunizations (Weeks 0, 8, 24). For the week 36 immunization, the adjuvant was changed to SMNP (Darrell Irvine lab, MIT). 750 U of SMNP was used per immunogen dose. Animals were immunized subcutaneously with antigen-adjuvant formulation. The dose was split in half and injected into both hind limbs. Blood draws were performed biweekly.

### ELISA binding assays

Sandwich ELISA experiments were performed using sera samples from weeks 6, 10, 26 and 38. All buffer additions and wash steps were performed using a BioStack Microplate Stacker system (BioTek). 12N antibody (as IgG) was diluted to 3 μg/ml and immobilized onto high-binding, 96-well microplates (Greiner Bio-One) for 2 hours at room temperature. The plates were washed three times with TBS + 0.1% Tween-20 (TBST). Plates were blocked with TBS + 5% bovine serum albumin (BSA) + 0.05% Tween-20 overnight at 4°C. Plates were washed three times with TBST before the addition of antigen solution (PBS + 1% BSA + 3 μg/ml of BG505 SOSIP). BG505 SOSIP constructs were matched to the antigen used for the immunization of each group of animals. Plates were treated with antigen for 2 hours at room temperature and subsequently washed three times with TBST. Serial two-fold sera dilutions (starting at 1:20) were prepared in PBS and loaded onto the antigen-coated plates (2-hour incubation at room temperature). Following three washes with TBST, AP-conjugated AffiniPure goat anti-human IgG, (Jackson Immunoresearch, Cat # 109-055-097) in TBS + 1% BSA was added for 1 hour at room temperature. Detection antibody was diluted 1:4000. Plates were washed three times with TBST, followed by the addition of 1-Step PNPP Substrate Solution (Thermo-Fisher Scientific). Colorimetric endpoint development was allowed to proceed for ~1 hour before termination by 2 M NaOH. Data was recorded on a Synergy H1 plate reader (BioTek) by measuring the absorbance at a wavelength of 405 nm. Midpoint titers were determined using Graphpad Prism software. Experiments were performed in triplicate.

### Pseudovirus inhibition assays

Pseudovirus inhibition (neutralization) assays were performed with Env-pseudotyped viruses and TZM-bl cells. The experiments with wt BG505 Env presented in Figure 1, Figure 6 and Table S3 were performed at The Scripps Research Institute, La Jolla, CA, USA, as described previously (Pauthner et al., 2019). The experiments involving mutant BG505 Env, presented in Table S6 were performed at Duke University Medical Center in Durham, NC, USA, as previously reported (Montefiori, 2009). Serial 3-fold dilutions of sera samples were pre-mixed with Env-pseudotyped virus and added to TZM-bl cells. Starting sera dilutions were 1:40 for the first and 1:20 for the latter group of experiments. Midpoint neutralization titers (ID_50_) were determined as the serum dilution at which pseudovirus infectivity was inhibited by 50%.

### nsEM-based polyclonal epitope mapping (nsEMPEM) – Preparation of Fab and complex samples

Experiments were implemented as described previously (Bianchi et al., 2018, Antanasijevic et al., 2020). Plasma samples (weeks 8, 10, 26, and 38) from three groups of six immunized rhesus macaques were chosen for polyclonal epitope mapping. IgGs were purified from ~1 ml of plasma with equal volume of settled Protein A Sepharose resin (GE Healthcare). Samples were eluted from the resin with 0.1 M glycine at pH 2.5 and immediately neutralized with 1 M Tris-HCl pH 8. Amicon ultrafiltration units with a 30 kDa cutoff (Millipore Sigma) were used to concentrate and buffer exchange the purified IgG to the digestion buffer (PBS + 10 mM EDTA + 20 mM cysteine, pH 7.4). IgG samples were digested for 10 hours at 37°C using 50 μl of settled papain-agarose resin (Thermo Fisher Scientific). Fc and non-digested IgG were removed through a 1-hour incubation at room temperature with Protein A Sepharose resin using 0.2 ml packed resin per 1 mg of IgG. Fab samples were concentrated to ~6-8 mg/ml and buffer exchanged to TBS using Amicon ultrafiltration units with a 10 kDa cutoff (EMD Millipore Sigma). Final Fab yields were ~1 mg. The complexes were assembled using 1 mg of purified polyclonal Fab and 15 μg of the corresponding BG505 SOSIP antigen used in the immunization (BG505 SOSIP MD39 for Grp 1 samples; BG505 SOSIP.v5.2 N241/N289 for Grp 2 samples; BG505 SOSIPv5.2(7S) N241/N289 for Grp 3 samples). For Fab samples from Rh.33311 and Rh.33172 (week 26 time point), the complexing was also performed with BG505 SOSIP.v3 under equivalent conditions. The reactions were incubated for ~18 hours at room temperature. Immune complexes were purified from residual Fab using SEC (Superose 6 Increase column) with TBS as a running buffer, concentrated with Amicon ultrafiltration units with a 10 kDa cutof, and immediately placed onto carbon-coated 400-mesh Cu grids as described in the nsEM section.

### Negative stain electron microscopy (nsEM)

Negative stain electron microscopy experiments were performed as described previously (Antanasijevic et al., 2020, Bianchi et al., 2018). Purified EMPEM complexes were diluted to 30-50 μg/ml and applied to carbon-coated 400-mesh Cu grids (glow-discharged at 15 mA for 25 s) for 10 s and then blotted. BG505 SOSIP-T33-31A and BG505 SOSIP-T33-31B trimer samples were diluted to 20 μg/ml and loaded onto grids following the same protocol. Grids were negatively stained using uranyl-formate, 2% (w/v), for 40 s. Data was collected on either a Tecnai Spirit electron microscope, operating at 120 keV, or a Tecnai TF20 electron microscope, operating at 200 keV. Nominal magnification was 52,000 X, with a pixel size of 2.05 Å (at the specimen plane) for the Spirit and 62,000 X, with a pixel size at 1.77 Å for the TF20. Electron dose was calibrated to 25 e^-^/Å^2^ and the defocus was set at −1.50 μm. Micrographs were recorded using a Tietz 4k x 4k TemCam-F416 CMOS camera in both cases. The Leginon automated imaging interface (Suloway et al., 2005) was used for data acquisition and the Appion data processing suite (Lander et al., 2009) was applied for initial processing steps. Relion/3.0 (Zivanov et al., 2018) was used for 2D and 3D classification steps.

### nsEMPEM – Data processing

Preliminary processing was conducted through the Appion data processing package where approximately 100,000–150,000 particles were auto-picked and extracted. Particles were then 2D-classified using Relion/3.0 into 250 classes (50 iterations), and particles with antigen-Fab qualities (roughly 50-80% of the original particles) were selected for 3D analysis. Initial 3D classification was performed using 40 classes. A low-resolution model of non-liganded HIV Env ectodomain was used as a reference for all 3D steps. Particles from similar looking classes were then combined and reclassified, and a subgroup of 3D classes with unique structural features was further processed using 3D auto-refinement (Relion 3.0). UCSF Chimera 1.13 (Pettersen et al., 2004) was used to visualize and segment the 3D refined maps. Finally, 3D refinement was conducted on a subgroup of particles selected after the 2D classification step (and prior to any 3D classification), and the refined model has been submitted to EMDB. Full particle stacks and 3D models used for Fab segmentation and generation of composite figures are available upon request.

### cryoEM-based polyclonal epitope mapping (cryoEMPEM) – Preparation of Fab and complex samples

We used the approach described above to generate 6-10 mg of purified polyclonal Fab samples from week 26 plasma extracted from animals Rh.32034, Rh.33104 and Rh.33311. For Rh.33172 animal, we used plasma collected at week 38. Polyclonal Fab samples were complexed with 200 μg of the corresponding BG505 SOSIP antigen used in the immunization (BG505 SOSIP MD39 for Rh.32034 and Rh.33104 samples; BG505 SOSIP.v5.2 N241/N289 for Rh.33311 sample; BG505 SOSIPv5.2(7S) N241/N289 for Rh.33172 sample) and incubated for ~18 hours at room temperature. Trimer-Fab immune complexes were SEC-purified (TBS was used as the running buffer) and concentrated to 5-7 mg/ml for application onto cryoEM grids.

### cryoEM-based polyclonal epitope mapping (cryoEMPEM) – Grid preparation and cryoEM imaging

A Vitrobot mark IV (Thermo Fisher Scientific) was used for cryo-grid preparation with the four immune complex samples described above. The temperature inside the chamber was set to 10°C, humidity was maintained at 100 %, blotting time was varied within a 4-7 s range, blotting force was set to 0, and wait time was set to 10 s. For cryo-grid preparation, we used lauryl maltose neopentyl glycol (LMNG) at a final concentration of 0.005 mM. Two types of grids were used: UltrAuFoil R 1.2/1.3 (Au, 300-mesh; Quantifoil Micro Tools GmbH) and Quantifoil R 2/1 (Cu, 400-mesh; Quantifoil Micro Tools GmbH). The grids were treated with Ar/O2 plasma (Solarus plasma cleaner, Gatan) for 10 s before sample application. An appropriate volume of 0.04 mM LMNG stock solution was mixed with the sample and 3 μl were immediately loaded onto the grid. After the blot step, the grids were plunge-frozen into liquid-nitrogen-cooled liquid ethane. Cryogrids were loaded into a FEI Titan Krios electron microscope (Thermo Fisher Scientific) operating at 300 kV. The microscope is equipped with a K2 Summit direct electron detector camera (Gatan) and sample autoloader. Exposure magnification was set to 29,000 with the resulting pixel size at the specimen plane of 1.03 Å. Leginon software was used for automated data collection (Suloway et al., 2005). Specific information on the imaging of individual polyclonal complex samples can be found in Table S4.

### cryoEM-based polyclonal epitope mapping (cryoEMPEM) – Data processing

Micrograph movie frames were aligned and dose-weighted using MotionCor2(Zheng et al., 2017). Initial data processing was performed in cryoSPARCv2.15 (Punjani et al., 2017). GCTF (Zhang, 2016) was used for estimation of CTF parameters. Immune-complex particles were picked using template picker, and two rounds of 2D classification were applied to eliminate bad particles picks and disassembled trimers. Particle stacks were transferred to Relion/3.0 (Zivanov et al., 2018) for further processing. After one round of 2D classification in Relion, the particles belonging to classes with appropriate trimer-Fab immune complex qualities were selected and subjected to a round of 3D refinement (C3 symmetry; soft solvent mask around the trimer core). A low-pass filtered map of BG505 SOSIP trimer was used as an initial model for all 3D steps to avoid initial model bias when reconstructing maps with unknown polyclonal Fabs. 3D-aligned particles were symmetry-expanded around the C3 axis because of the trimeric nature of the antigen. This expansion collapses all epitope-paratope interfaces onto a single protomer, which simplifies the 3D classification process. However, to prevent symmetry-related copies of individual particles from aligning to themselves, particle alignment was constrained in the subsequent 3D classification and refinement steps. 3D classification steps were performed without image alignment (--skip_align, T = 16) and 3D refinement steps on the selected subsets of particles were performed with local angular searches only (starting at 3.7° or 1.8° per iteration). The first round of 3D classification was run with an 80 Å sphere mask around the epitope-paratope interface and centered on the Fab-corresponding density. This is done separately for each epitope cluster where polyclonal Fabs can be detected. The number of output 3D classes was adjusted for each epitope based on the relative occupancy of bound polyclonal Fabs (typically 8-30 classes). 80 Å sphere masks allowed for sorting on the absence/presence of Fabs at each site as well as the relative orientation. 3D classes of particles featuring structurally unique polyclonal Fabs (in terms of epitopes and orientation) were selected separately and processed independently from this point on. After a round of 3D refinement for all selected subsets of particles featuring structurally unique polyclonal Fabs, the 2^nd^ round of 3D classification was run (120 Å sphere mask; K = 3; T = 16). The larger mask allowed sorting based on the epitope-paratope features. 3D classes of particles with the highest quality and resolution were selected and subjected to 3D refinement. Soft solvent masks around the specific trimer-Fab complex were used. A final round of 3D classification was run with a full trimer-Fab complex mask prepared in the previous step (K = 3; T = 16). The highest-resolution classes were selected for every unique trimer-Fab complex, 3D-refined and postprocessed (solvent mask around the complex; MTF correction). The postprocessed maps were used for model building and submission to EMDB. This workflow is further illustrated in Figure S2. Relevant data processing information (particle count at different stages, apparent map resolutions, B-factors used for map sharpening) are presented in Table S5.

### cryoEM-based polyclonal epitope mapping (cryoEMPEM) – Model building and refinement

Postprocessed maps from Relion generated in the previous step were used for model building and refinement. The BG505 SOSIP structure from PDB entry 6vfl (Antanasijevic et al., 2020) was used as an initial model for the antigen-corresponding parts of the maps. The sequence was adjusted to match the exact BG505 SOSIP variant of the imaged polyclonal complexes (BG505 SOSIP MD39, BG505 SOSIP.v5.2 N241/N289 or BG505 SOSIP.v5.2(7S) N241/N289).

Initial Fab models were generated from PDB entry 4kte (Tran et al., 2014) by mutating all of the amino acids to alanine. The BG505 SOSIP and Fab models were docked into each map in UCSF Chimera to generate a starting model for refinement. Structural homology to published rhesus macaque antibody structures (PDB IDs: 4KTE, 4KTD, 4RFE, 4Q2Z) was used to assign antibody heavy (H) and light (L) chains (Gohain et al., 2015, Tran et al., 2014, Navis et al., 2014). The principal factors for assignment were the comparisons of CDR2 and CDR3 lengths and the conformations of FR2 and FR3 regions between the H and L chains in the reconstructed cryoEM maps. Iterative rounds of manual model refinement in Coot (Emsley and Crispin, 2018) and automated model refinement in Rosetta (Wang et al., 2016) were used to refine the models into the reconstructed maps. CDR lengths of poly-alanine Fab models had to be adjusted by insertion/deletion of one or more alanine residues to match with the corresponding structural constraints imposed by the cryoEM maps. Final models were evaluated using MolProbity (Williams et al., 2018) and EMRinger (Barad et al., 2015), and the refinement statistics are shown in Table S5. The structures were submitted to the PDB database.

### CryoEM analysis of BG505 SOSIP-T33-31 nanoparticle – Grid preparation and cryoEM imaging

BG505 SOSIP-T33-31 nanoparticle sample (in TBS) was concentrated to 4.1 mg/ml for cryo-grid application. Cryo-grids were prepared using a Vitrobot mark IV under the equivalent conditions as the polyclonal immune-complex samples described above. BG505 SOSIP-T33-31 sample, premixed with 0.005 mM LMNG, was loaded onto plasma-cleaned Quantifoil R 2/1 (Cu, 400-mesh; Quantifoil Micro Tools GmbH) grids. Following the blotting step, the grids were plunge-frozen into liquid-nitrogen-cooled liquid ethane. Cryo-grids were imaged on an FEI Talos Arctica (Thermo Fisher Scientific) microscope operating at 200 kV, equipped with a K2 Summit direct electron detector camera (Gatan) and sample autoloader. Exposure magnification was set to 36,000 and the pixel size at the specimen plane was 1.15 Å. Automated data collection was performed using the Leginon software suite (Suloway et al., 2005). Data collection information can be found in Table S4.

### CryoEM analysis of BG505 SOSIP-T33-31 nanoparticle – Data processing

Micrograph movie frames were aligned and dose-weighted using MotionCor2 (Zheng et al., 2017). All further processing steps were performed in Relion/3.0 (Zivanov et al., 2018). GCTF (Zhang, 2016) was used for estimation of CTF parameters. 233,999 particles were auto-picked and extracted. Following a round of 2D classification, 214,224 particles were selected for 3D steps. Several iterative rounds of 3D refinement and 3D classification in Relion were performed to identify a subpopulation of 110,369 particles that were used in the final 3D reconstruction of the nanoparticle core. Tetrahedral symmetry (T) was applied for the 3D steps. A soft solvent mask around the T33-31 nanoparticle core was introduced for the final 3D classification, refinement and postprocessing steps. This solvent mask excluded the density corresponding to flexibly-linked BG505 SOSIP antigens. The final map resolution of the T33-31 nanoparticle core was 2.9 Å after postprocessing. BG505 SOSIP trimers were connected to both nanoparticle components via flexible linkers. The flexibility prevents a joint analysis with the nanoparticle core. We applied localized reconstruction v1.2.0 (Ilca et al., 2015) to extract BG505 SOSIP trimer subparticles connected to T33-31A and T33-31B components (termed BG505 SOSIP(A) and BG505 SOSIP(B)). Subparticle vectors were defined by marker files generated in UCSF Chimera (Pettersen et al., 2004), and trimer subparticles were extracted from pre-aligned nanoparticles after the subtraction of the signal corresponding to the nanoparticle core. Each nanoparticle displayed four trimers on each component (eight total) so the number of extracted BG505 SOSIP(A) and BG505 SOSIP(B) subparticles was 441,476 (4 × 110,369). A and B subsets were processed independently in Relion. Each subparticle subset was subjected to two rounds of 2D classification and one round of 3D classification. After eliminating the cropped and low-resolution classes of subparticles, 106,478 subparticles of BG505 SOSIP(A) and 64,726 subparticles corresponding to BG505 SOSIP(B) were subjected to 3D autorefinements with C3 symmetry. A soft solvent mask was applied for refinement and postprocessing steps. Final map resolutions were 3.7Å and 3.8Å for BG505 SOSIP(A) and BG505 SOSIP(B). Relevant data processing parameters are reported in Table S5.

### CryoEM analysis of BG505 SOSIP-T33-31 nanoparticle – Model building and refinement

Postprocessed maps corresponding to the T33-31 nanoparticle core, BG505 SOSIP(A) and BG505 SOSIP(B) generated in the previous step were used to generate atomic models. Crystal structures of non-liganded T33-31 nanoparticle (PDB entry 4zk7) and BG505 SOSIP trimer (PDB entry 6vfl) were used as initial models for refinement of the T33-31 nanoparticle core and BG505 SOSIP(A) and (B), respectively(Antanasijevic et al., 2020, Bale et al., 2015). Symmetry was applied for all automated refinement steps (T for the nanoparticle and C3 for the BG505 SOSIP trimers). Several rounds of Rosetta relaxed refinement (Wang et al., 2016) and manual refinement in Coot (Emsley and Crispin, 2018) were performed to generate final models. EMRinger (Barad et al., 2015) and MolProbity (Williams et al., 2018) analyses were used for model validation and generation of statistics reported in Table S5. Refined models were submitted to the Protein Data Bank.

## Author contributions

A.B.W., S.C., W.R.S., D.R.B. and G.S. conceived the rhesus macaque immunization study. C.A.C., D.G.C., and A.A. helped design the immunization experiments. D.G.C. L.E.J., J.T.N., and J.B.S. executed the immunization experiments and collected serum/plasma samples. A.A. produced the BG505 SOSIP-bearing T33-31 nanoparticles and performed structural and antigenic characterization. Z.T.B. helped with processing of cryoEM data. J.C. helped produce the T33-31 nanoparticle components. C.A.C. engineered and produced the BG505 SOSIP.v5.2 N241/N289 immunogen. E.G., and B.G. produced the BG505 SOSIP MD39 immunogen. J.D.A. ran sitespecific glycosylation analysis on purified antigen-bearing nanoparticle components. C.A.C., L.M.S. and A.A. optimized and performed the ELISA experiments. J.B., R.B. and C.L. performed the pseudovirus neutralization experiments. C.A.C., L.M.S. and A.A. executed the nsEMPEM experiments. L.M.S. isolated Fabs from plasma samples for all EMPEM experiments. A.A. acquired and processed the cryoEMPEM data. H.R.P., K.R., F.C., Y.R.Y., A.T.d.l.P., R.F.R. and A.A. built and refined the atomic models into the cryoEM maps. A.B.W conceptualized the study. A.B.W., G.S., S.C., W.R.S., D.R.B., M.C. and D.C.M. supervised the study and provided essential guidance. D.B., N.P.K., R.W.S. and J.P.M. gave critical feedback regarding study design and data interpretation. A.B.W. and A.A. wrote the original draft of the manuscript. All authors contributed to the manuscript text by assisting in writing and/or providing critical feedback.

## Acknowledgements

The authors express sincere gratitude to Darrell Irvine lab (MIT) and Novavax. Inc, for providing the SMNP and Matrix-M™ adjuvants used in the immunization experiments. The authors thank Bill Anderson, Hannah L. Turner, Charles A. Bowman and Jean-Christophe Ducom (The Scripps Research Institute) for their help with electron microscopy, data acquisition and processing. The authors also acknowledge Lauren Holden for her help on the preparation of this manuscript.

## Declaration of Interests

### Funding statement for submission

This work was supported by grants from the National Institute of Allergy and Infectious Diseases, Center for HIV/AIDS Vaccine Immunology and Immunogen Discovery UM1AI100663 (M.C., A.B.W., D.R.B., G.S.), Center for HIV/AIDS Vaccine Development UM1AI144462 (M.C., A.B.W., D.R.B., G.S.), P01 AI110657 (J.P.M., R.W.S., and A.B.W.); and by the Bill and Melinda Gates Foundation and the Collaboration for AIDS Vaccine Discovery (CAVD) OPP1156262 (N.P.K., D.B.), OPP1115782/INV-002916 (A.B.W.), OPP1132237 (J.P.M.), OPP1146996 (D.C.M.), INV-002022 (R.W.S.) and OPP1196345 (A.B.W.); and by the National Science Foundation grant DMREF 1629214 (N.P.K., D.B.); and by the Howard Hughes Medical Institute (D.B.). Yerkes National Primate Research Center is supported by the base grant P51 OD011132. This work was also supported by the European Union’s Horizon 2020 research and innovation program under grant agreement No. 681137 (M.C., R.W.S.). C.A.C. is supported by a NIH F31 Ruth L. Kirschstein Predoctoral Award Al131873 and by the Achievement Rewards for College Scientists Foundation. R.W.S. is supported by the Vici fellowship from the Netherlands Organization for Scientific Research (NWO). This work was partially funded by IAVI with the generous support of USAID, Ministry of Foreign Affairs of the Netherlands, and the Bill & Melinda Gates Foundation; a full list of IAVI donors is available at www.iavi.org. The contents of this manuscript are the responsibility of the authors and do not necessarily reflect the views of USAID or the US Government. The funders had no role in study design, data collection and analysis, decision to publish, or preparation of the manuscript.

### Data availability statement

3D maps and models from the EM analysis have been deposited to the Electron Microscopy Databank (http://www.emdatabank.org/) and the Protein Data Bank (http://www.rcsb.org/), respectively. The accession numbers are listed in the table below.

| Sample description | EMDB ID | PDB ID |
| --- | --- | --- |
| nsEM maps of Grp 1 polyclonal samples (Wk 26) | 23175 | N/A |
| nsEM maps of Grp 2 polyclonal samples (Wk 26) | 23176 | N/A |
| nsEM maps of Grp 3 polyclonal samples (Wk 8) | 23177 | N/A |
| nsEM maps of Grp 3 polyclonal samples (Wk 10) | 23178 | N/A |
| nsEM maps of Grp 3 polyclonal samples (Wk 26) | 23179 | N/A |
| nsEM maps of Grp 3 polyclonal samples (Wk 38) | 23180 | N/A |
| nsEM map of Rh.33172 (Wk 26) polyclonal sample (antigen: BG505 SOSIP.v3) | 23182 | N/A |
| nsEM map of Rh.33172 (Wk 26) polyclonal sample (antigen: BG505 SOSIP.v5.2(7S) N241/N289) | 23181 | N/A |
| nsEM map of Rh.33311 (Wk 26) polyclonal sample (antigen: BG505 SOSIP.v3) | 23183 | N/A |
| nsEM map of Rh.33311 (Wk 26) polyclonal sample (antigen: BG505 SOSIP.v5.2(7S) N241/N289) | 23184 | N/A |
| nsEM map of the BG505 SOSIP-T33-31A nanoparticle component | 23185 | N/A |
| nsEM map of the BG505 SOSIP-T33-31A nanoparticle component | 23186 | N/A |
| cryoEM map and model of the BG505 SOSIP (A) subparticle of the BG505 SOSIP-T33-31 nanoparticle | 23218 | 7L7T |
| cryoEM map of the BG505 SOSIP (B) subparticle of the BG505 SOSIP-T33-31 nanoparticle | 23219 | 7L7U |
| cryoEM map and model of the T33-31 core subparticle of the BG505 SOSIP-T33-31 nanoparticle | 23222 | 7L85 |
| cryoEM map and model of the Rh.32034 pAbC-1 (antigen: BG505 SOSIP MD39) | 23223 | 7L86 |
| cryoEM map and model of the Rh.32034 pAbC-2 (antigen: BG505 SOSIP MD39) | 23224 | 7L87 |
| cryoEM map and model of the Rh.32034 pAbC-3 (antigen: BG505 SOSIP MD39) | 23225 | 7L88 |
| cryoEM map and model of the Rh.32034 pAbC-4 (antigen: BG505 SOSIP MD39) | 23226 | 7L89 |
| cryoEM map and model of the Rh.33104 pAbC-1 (antigen: BG505 SOSIP MD39) | 23227 | 7L8A |
| cryoEM map and model of the Rh.33104 pAbC-2 (antigen: BG505 SOSIP MD39) | 23228 | 7L8B |
| cryoEM map and model of the Rh.33104 pAbC-3 (antigen: BG505 SOSIP MD39) | 23229 | 7L8C |
| cryoEM map and model of the Rh.33104 pAbC-4 (antigen: BG505 SOSIP MD39) | 23230 | 7L8D |
| cryoEM map and model of the Rh.33311 pAbC-1 (antigen: BG505 SOSIP.v5.2 N241/N289) | 23236 | 7L8T |
| cryoEM map and model of the Rh.33311 pAbC-2 (antigen: BG505 SOSIP.v5.2 N241/N289) | 23237 | 7L8U |
| cryoEM map and model of the Rh.33311 pAbC-3 (antigen: BG505 SOSIP.v5.2 N241/N289) | 23238 | 7L8W |
| cryoEM map and model of the Rh.33311 pAbC-4 (antigen: BG505 SOSIP.v5.2 N241/N289) | 23239 | 7L8X |
| cryoEM map and model of the Rh.33311 pAbC-5 (antigen: BG505 SOSIP.v5.2 N241/N289) | 23240 | 7L8Y |
| cryoEM map of the Rh.33311 pAbC-6 (antigen: BG505 SOSIP.v5.2 N241/N289) | 23241 | N/A |
| cryoEM map and model of the Rh.33311 pAbC-7 (antigen: BG505 SOSIP.v5.2 N241/N289) | 23242 | 7L8Z |
| cryoEM map and model of the Rh.33311 pAbC-8 (antigen: BG505 SOSIP.v5.2 N241/N289) | 23243 | 7L90 |
| cryoEM map and model of the Rh.33172 pAbC-1 (antigen: BG505 SOSIP.v5.2(7S) N241/N289) | 23231 | 7L8E |
| cryoEM map and model of the Rh.33172 pAbC-2 (antigen: BG505 SOSIP.v5.2(7S) N241/N289) | 23232 | 7L8F |
| cryoEM map and model of the Rh.33172 pAbC-3 (antigen: BG505 SOSIP.v5.2(7S) N241/N289) | 23233 | 7L8G |
| cryoEM map and model of the Rh.33172 pAbC-4 (antigen: BG505 SOSIP.v5.2(7S) N241/N289) | 23235 | 7L8S |
| cryoEM map of the Rh.33172 pAbC-5 (antigen: BG505 SOSIP.v5.2(7S) N241/N289) | 23234 | N/A |

## Notes

### Competing Interest Statement

The authors have declared no competing interest.

## Literature

Antanasijevic, A., Ueda, G., Brouwer, P. J. M., Copps, J., Huang, D., Allen, J. D., Cottrell, C.A., Yasmeen, A., Sewall, L. M., Bontjer, I., Ketas, T. J., Turner, H. L., Berndsen, Z. T., Montefiori, D. C., Klasse, P. J., Crispin, M., Nemazee, D., Moore, J. P., Sanders, R. W., King, N. P., Baker, D. & Ward, A. B. 2020. Structural and functional evaluation of de novo-designed, two-component nanoparticle carriers for HIV Env trimer immunogens. PLoS Pathog, 16, e1008665.

Bale, J. B., Park, R. U., Liu, Y., Gonen, S., Gonen, T., Cascio, D., King, N. P., Yeates, T. O. & Baker, D. 2015. Structure of a designed tetrahedral protein assembly variant engineered to have improved soluble expression. Protein Sci, 24, 1695–701.

Barad, B. A., Echols, N., Wang, R. Y., Cheng, Y., Dimaio, F., Adams, P. D. & Fraser, J. S. 2015. EMRinger: side chain-directed model and map validation for 3D cryo-electron microscopy. Nat Methods, 12, 943–6.

Behrens, A. J., Vasiljevic, S., Pritchard, L. K., Harvey, D. J., Andev, R. S., Krumm, S. A., Struwe, W. B., Cupo, A., Kumar, A., Zitzmann, N., Seabright, G. E., Kramer, H. B., Spencer, D. I., Royle, L., Lee, J. H., Klasse, P. J., Burton, D. R., Wilson, I. A., Ward, A. B., Sanders, R. W., Moore, J. P., Doores, K. J. & Crispin, M. 2016. Composition and Antigenic Effects of Individual Glycan Sites of a Trimeric HIV-1 Envelope Glycoprotein. Cell Rep, 14, 2695–706.

Berndsen, Z. T., Chakraborty, S., Wang, X., Cottrell, C. A., Torres, J. L., Diedrich, J. K., Lopez, C. A., Yates, J. R., 3RD, van Gils, M. J., Paulson, J. C., Gnanakaran, S. & Ward, A. B. 2020. Visualization of the HIV-1 Env glycan shield across scales. Proc Natl Acad Sci U S A, 117, 28014–28025.

Bianchi, M., Turner, H. L., Nogal, B., Cottrell, C. A., Oyen, D., Pauthner, M., Bastidas, R., Nedellec, R., Mccoy, L. E., Wilson, I. A., Burton, D. R., Ward, A. B. & Hangartner, L. 2018. Electron-Microscopy-Based Epitope Mapping Defines Specificities of Polyclonal Antibodies Elicited during HIV-1 BG505 Envelope Trimer Immunization. Immunity, 49, 288–300 e8.

Brouwer, P. J. M., Antanasijevic, A., Berndsen, Z., Yasmeen, A., Fiala, B., Bijl, T. P. L., Bontjer, I., Bale, J. B., Sheffler, W., Allen, J. D., Schorcht, A., Burger, J. A., Camacho, M., Ellis, D., Cottrell, C. A., Behrens, A. J., Catalano, M., del Moral-Sanchez, I., Ketas, T. J., Labranche, C., van Gils, M. J., Sliepen, K., Stewart, L. J., Crispin, M., Montefiori, D. C., Baker, D., Moore, J. P., Klasse, P. J., Ward, A. B., King, N. P. & Sanders, R. W. 2019. Enhancing and shaping the immunogenicity of native-like HIV-1 envelope trimers with a two-component protein nanoparticle. Nat Commun, 10, 4272.

Brouwer, P. J. M. & Sanders, R. W. 2019. Presentation of HIV-1 envelope glycoprotein trimers on diverse nanoparticle platforms. Curr Opin HIV AIDS, 14, 302–308.

Burton, D. R. & Hangartner, L. 2016. Broadly Neutralizing Antibodies to HIV and Their Role in Vaccine Design. Annu Rev Immunol, 34, 635–59.

Cao, L., Diedrich, J. K., Kulp, D. W., Pauthner, M., He, L., Park, S. R., Sok, D., Su, C. Y., Delahunty, C. M., Menis, S., Andrabi, R., Guenaga, J., Georgeson, E., Kubitz, M., Adachi, Y., Burton, D. R., Schief, W. R., Yates, J. R., III & Paulson, J. C. 2017. Global site-specific N-glycosylation analysis of HIV envelope glycoprotein. Nat Commun, 8, 14954.

Chuang, G. Y., Geng, H., Pancera, M., Xu, K., Cheng, C., Acharya, P., Chambers, M., Druz, A., Tsybovsky, Y., Wanninger, T. G., Yang, Y., Doria-Rose, N. A., Georgiev, I. S., Gorman, J., Joyce, M. G., O’Dell, S., Zhou, T., Mcdermott, A. B., Mascola, J. R. & Kwong, P. D. 2017. Structure-Based Design of a Soluble Prefusion-Closed HIV-1 Env Trimer with Reduced CD4 Affinity and Improved Immunogenicity. J Virol, 91.

Cirelli, K. M., Carnathan, D. G., Nogal, B., Martin, J. T., Rodriguez, O. L., Upadhyay, A. A., Enemuo, C. A., Gebru, E. H., Choe, Y., Viviano, F., Nakao, C., Pauthner, M. G., Reiss, S., Cottrell, C. A., Smith, M. L., Bastidas, R., Gibson, W., Wolabaugh, A. N., Melo, M. B., Cossette, B., Kumar, V., Patel, N. B., Tokatlian, T., Menis, S., Kulp, D. W., Burton, D. R., Murrell, B., Schief, W. R., Bosinger, S. E., Ward, A. B., Watson, C. T., Silvestri, G., Irvine, D. J. & Crotty, S. 2019. Slow Delivery Immunization Enhances HIV Neutralizing Antibody and Germinal Center Responses via Modulation of Immunodominance. Cell, 177, 1153–1171 e28.

Cottrell, C. A., van Schooten, J., Bowman, C. A., Yuan, M., Oyen, D., Shin, M., Morpurgo, R., van der Woude, P., van Breemen, M., Torres, J. L., Patel, R., Gross, J., Sewall, L. M., Copps, J., Ozorowski, G., Nogal, B., Sok, D., Rakasz, E. G., Labranche, C., Vigdorovich, V., Christley, S., Carnathan, D. G., Sather, D. N., Montefiori, D., Silvestri, G., Burton, D. R., Moore, J. P., Wilson, I. A., Sanders, R. W., Ward, A. B. & van Gils, M. J. 2020. Mapping the immunogenic landscape of near-native HIV-1 envelope trimers in non-human primates. PLoS Pathog, 16, e1008753.

de Taeye, S. W., Ozorowski, G., Torrents de la Pena, A., Guttman, M., Julien, J. P., van den Kerkhof, T. L., Burger, J. A., Pritchard, L. K., Pugach, P., Yasmeen, A., Crampton, J., Hu, J., Bontjer, I., Torres, J. L., Arendt, H., Destefano, J., Koff, W. C., Schuitemaker, H., Eggink, D., Berkhout, B., Dean, H., Labranche, C., Crotty, S., Crispin, M., Montefiori, D. C., Klasse, P. J., Lee, K. K., Moore, J. P., Wilson, I. A., Ward, A. B. & Sanders, R. W. 2015. Immunogenicity of Stabilized HIV-1 Envelope Trimers with Reduced Exposure of Non-neutralizing Epitopes. Cell, 163, 1702–15.

Derking, R., Allen, J. D., Cottrell, C. A., Sliepen, K., Seabright, G. E., Lee, W. H., Rantalainen, K., Antanasijevic, A., Copps, J., Yasmeen, A., van der Woude, P., De TAEYE, S. W., van den Kerkhof, T. L., Klasse, P. J., van Gils, M. J., Moore, J. P., Ward, A. B., Crispin, M. & Sanders, R. W. 2020. Enhancing glycan occupancy of soluble HIV-1 envelope trimers to mimic the native viral spike. bioRxiv.

Dey, A. K., Cupo, A., Ozorowski, G., Sharma, V. K., Behrens, A. J., Go, E. P., Ketas, T. J., Yasmeen, A., Klasse, P. J., Sayeed, E., Desaire, H., Crispin, M., Wilson, I. A., Sanders, R. W., Hassell, T., Ward, A. B. & Moore, J. P. 2018. cGMP production and analysis of BG505 SOSIP.664, an extensively glycosylated, trimeric HIV-1 envelope glycoprotein vaccine candidate. Biotechnol Bioeng, 115, 885–899.

Emsley, P. & Crispin, M. 2018. Structural analysis of glycoproteins: building N-linked glycans with Coot. Acta Crystallogr D Struct Biol, 74, 256–263.

Escolano, A., Gristick, H. B., Abernathy, M. E., Merkenschlager, J., Gautam, R., Oliveira, T. Y., Pai, J., West, A. P., JR., Barnes, C. O., Cohen, A. A., Wang, H., Golijanin, J., Yost, D., Keeffe, J. R., Wang, Z., Zhao, P., Yao, K. H., Bauer, J., Nogueira, L., Gao, H., Voll, A. V., Montefiori, D. C., Seaman, M. S., Gazumyan, A., Silva, M., Mcguire, A. T., Stamatatos, L., Irvine, D. J., Wells, L., Martin, M. A., Bjorkman, P. J. & Nussenzweig, M. C. 2019. Immunization expands B cells specific to HIV-1 V3 glycan in mice and macaques. Nature, 570, 468–473.

Georgiev, I. S., Joyce, M. G., Chen, R. E., Leung, K., Mckee, K., Druz, A., van Galen, J. G., Kanekiyo, M., Tsybovsky, Y., Yang, E. S., Yang, Y., Acharya, P., Pancera, M., Thomas, P. V., Wanninger, T., Yassine, H. M., Baxa, U., Doria-Rose, N. A., Cheng, C., Graham, B. S., Mascola, J. R. & Kwong, P. D. 2018. Two-Component Ferritin Nanoparticles for Multimerization of Diverse Trimeric Antigens. ACS Infect Dis, 4, 788–796.

Gohain, N., Tolbert, W. D., Acharya, P., Yu, L., Liu, T., Zhao, P., Orlandi, C., Visciano, M. L., Kamin-Lewis, R., Sajadi, M. M., Martin, L., Robinson, J. E., Kwong, P. D., Devico, A. L., Ray, K., Lewis, G. K. & Pazgier, M. 2015. Cocrystal Structures of Antibody N60-i3 and Antibody JR4 in Complex with gp120 Define More Cluster A Epitopes Involved in Effective Antibody-Dependent Effector Function against HIV-1. J Virol, 89, 8840–54.

Havenar-Daughton, C., Lee, J. H. & Crotty, S. 2017. Tfh cells and HIV bnAbs, an immunodominance model of the HIV neutralizing antibody generation problem. Immunol Rev, 275, 49–61.

He, L., De Val, N., Morris, C. D., Vora, N., Thinnes, T. C., Kong, L., Azadnia, P., Sok, D., Zhou, B., Burton, D. R., Wilson, I. A., Nemazee, D., Ward, A. B. & Zhu, J. 2016. Presenting native-like trimeric HIV-1 antigens with self-assembling nanoparticles. Nat Commun, 7, 12041.

He, L., Kumar, S., Allen, J. D., Huang, D., Lin, X., Mann, C. J., Saye-Francisco, K. L., Copps, J., Sarkar, A., Blizard, G. S., Ozorowski, G., Sok, D., Crispin, M., Ward, A. B., Nemazee, D., Burton, D. R., Wilson, I. A. & Zhu, J. 2018. HIV-1 vaccine design through minimizing envelope metastability. Sci Adv, 4, eaau6769.

Hu, J. K., Crampton, J. C., Cupo, A., Ketas, T., van Gils, M. J., Sliepen, K., De TAEYE, S. W., Sok, D., Ozorowski, G., Deresa, I., Stanfield, R., Ward, A. B., Burton, D. R., Klasse, P. J., Sanders, R. W., Moore, J. P. & Crotty, S. 2015. Murine Antibody Responses to Cleaved Soluble HIV-1 Envelope Trimers Are Highly Restricted in Specificity. J Virol, 89, 10383–98.

Ilca, S. L., Kotecha, A., Sun, X., Poranen, M. M., Stuart, D. I. & Huiskonen, J. T. 2015. Localized reconstruction of subunits from electron cryomicroscopy images of macromolecular complexes. Nat Commun, 6, 8843.

Jardine, J., Julien, J. P., Menis, S., Ota, T., Kalyuzhniy, O., McGuire, A., Sok, D., Huang, P. S., Macpherson, S., Jones, M., Nieusma, T., Mathison, J., Baker, D., Ward, A. B., Burton, D. R., Stamatatos, L., Nemazee, D., Wilson, I. A. & Schief, W. R. 2013. Rational HIV immunogen design to target specific germline B cell receptors. Science, 340, 711–6.

Jardine, J. G., Ota, T., Sok, D., Pauthner, M., Kulp, D. W., Kalyuzhniy, O., Skog, P. D., Thinnes, T. C., Bhullar, D., Briney, B., Menis, S., Jones, M., Kubitz, M., Spencer, S., Adachi, Y., Burton, D. R., Schief, W. R. & Nemazee, D. 2015. HIV-1 VACCINES. Priming a broadly neutralizing antibody response to HIV-1 using a germline-targeting immunogen. Science, 349, 156–61.

King, N. P., Bale, J. B., Sheffler, W., Mcnamara, D. E., Gonen, S., Gonen, T., Yeates, T. O. & Baker, D. 2014. Accurate design of co-assembling multi-component protein nanomaterials. Nature, 510, 103–8.

Klasse, P. J., Ketas, T. J., Cottrell, C. A., Ozorowski, G., Debnath, G., Camara, D., Francomano, E., Pugach, P., Ringe, R. P., Labranche, C. C., van Gils, M. J., Bricault, C. A., Barouch, D. H., Crotty, S., Silvestri, G., Kasturi, S., Pulendran, B., Wilson, I. A., Montefiori, D. C., Sanders, R. W., Ward, A. B. & Moore, J. P. 2018. Epitopes for neutralizing antibodies induced by HIV-1 envelope glycoprotein BG505 SOSIP trimers in rabbits and macaques. PLoS Pathog, 14, e1006913.

Klasse, P. J., Ozorowski, G., Sanders, R. W. & Moore, J. P. 2020. Env Exceptionalism: Why Are HIV-1 Env Glycoproteins Atypical Immunogens? Cell Host Microbe, 27, 507–518.

Kong, R., Duan, H., Sheng, Z., Xu, K., Acharya, P., Chen, X., Cheng, C., Dingens, A. S., Gorman, J., Sastry, M., Shen, C. H., Zhang, B., Zhou, T., Chuang, G. Y., Chao, C. W., Gu, Y., Jafari, A. J., Louder, M. K., O’Dell, S., Rowshan, A. P., Viox, E. G., Wang, Y., Choi, C. W., Corcoran, M. M., Corrigan, A. R., Dandey, V. P., Eng, E. T., Geng, H., Foulds, K. E., Guo, Y., Kwon, Y. D., Lin, B., Liu, K., Mason, R. D., Nason, M. C., Ohr, T. Y., Ou, L., Rawi, R., Sarfo, E. K., Schon, A., Todd, J. P., Wang, S., Wei, H., Wu, W., Program, N. C. S., Mullikin, J. C., Bailer, R. T., Doria-Rose, N. A., Karlsson Hedestam, G. B., Scorpio, D. G., Overbaugh, J., Bloom, J. D., Carragher, B., Potter, C. S., Shapiro, L., Kwong, P. D. & Mascola, J. R. 2019. Antibody Lineages with Vaccine-Induced Antigen-Binding Hotspots Develop Broad HIV Neutralization. Cell, 178, 567–584 e19.

Labranche, C. C., McGuire, A. T., Gray, M. D., Behrens, S., Kwong, P. D. K., Chen, X., Zhou, T., Sattentau, Q. J., Peacock, J., Eaton, A., Greene, K., Gao, H., Tang, H., Perez, L. G., Chen, X., Saunders, K. O., Kwong, P. D., Mascola, J. R., Haynes, B. F., Stamatatos, L. & Montefiori, D. C. 2018. HIV-1 envelope glycan modifications that permit neutralization by germline-reverted VRC01-class broadly neutralizing antibodies. PLoS Pathog, 14, e1007431.

Lander, G. C., Stagg, S. M., Voss, N. R., Cheng, A., Fellmann, D., Pulokas, J., Yoshioka, C., Irving, C., Mulder, A., Lau, P. W., Lyumkis, D., Potter, C. S. & Carragher, B. 2009. Appion: an integrated, database-driven pipeline to facilitate EM image processing. J Struct Biol, 166, 95–102.

Lei, L., Yang, Y. R., Tran, K., Wang, Y., Chiang, C. I., Ozorowski, G., Xiao, Y., Ward, A. B., Wyatt, R. T. & Li, Y. 2019. The HIV-1 Envelope Glycoprotein C3/V4 Region Defines a Prevalent Neutralization Epitope following Immunization. Cell Rep, 27, 586–598 e6.

Martin, J. T., Cottrell, C. A., Antanasijevic, A., Carnathan, D. G., Cossette, B., Enemuo, C. A., Gebru, E. H., Choe, Y., Viviano, F., Tokatlian, T., Cirelli, K. M., Ueda, G., Copps, J., Schiffner, T., Menis, S., Schief, W. R., Crotty, S., King, N. P., Silvestri, G., Ward, A. B. & Irvine, D. J. 2020. Targeting HIV Env immunogens to B cell follicles in non-human primates through immune complex or protein nanoparticle formulations. bioRxiv.

Mccoy, L. E., van Gils, M. J., Ozorowski, G., Messmer, T., Briney, B., Voss, J. E., Kulp, D. W., Macauley, M. S., Sok, D., Pauthner, M., Menis, S., Cottrell, C. A., Torres, J. L., Hsueh, J., Schief, W. R., Wilson, I. A., Ward, A. B., Sanders, R. W. & Burton, D. R. 2016. Holes in the Glycan Shield of the Native HIV Envelope Are a Target of Trimer-Elicited Neutralizing Antibodies. Cell Rep, 16, 2327–38.

Medina-Ramirez, M., Garces, F., Escolano, A., Skog, P., de Taeye, S. W., del Moral-Sanchez, I., Mcguire, A. T., Yasmeen, A., Behrens, A. J., Ozorowski, G., van den Kerkhof, T., Freund, N. T., Dosenovic, P., Hua, Y., Gitlin, A. D., Cupo, A., van der Woude, P., Golabek, M., Sliepen, K., Blane, T., Kootstra, N., van Breemen, M. J., Pritchard, L. K., Stanfield, R. L., Crispin, M., Ward, A. B., Stamatatos, L., Klasse, P. J., Moore, J. P., Nemazee, D., Nussenzweig, M. C., Wilson, I. A. & Sanders, R. W. 2017. Design and crystal structure of a native-like HIV-1 envelope trimer that engages multiple broadly neutralizing antibody precursors in vivo. J Exp Med, 214, 2573–2590.

Montefiori, D. C. 2009. Measuring HIV neutralization in a luciferase reporter gene assay. Methods Mol Biol, 485, 395–405.

Moyer, T. J., Kato, Y., Abraham, W., Chang, J. Y. H., Kulp, D. W., Watson, N., Turner, H. L., Menis, S., Abbott, R. K., Bhiman, J. N., Melo, M. B., Simon, H. A., Herrera-de la Mata, S., Liang, S., Seumois, G., Agarwal, Y., Li, N., Burton, D. R., Ward, A. B., Schief, W. R., Crotty, S. & Irvine, D. J. 2020. Engineered immunogen binding to alum adjuvant enhances humoral immunity. Nat Med, 26, 430–440.

Navis, M., Tran, K., Bale, S., Phad, G. E., Guenaga, J., Wilson, R., Soldemo, M., Mckee, K., Sundling, C., Mascola, J., Li, Y., Wyatt, R. T. & Karlsson Hedestam, G. B. 2014. HIV- 1 receptor binding site-directed antibodies using a VH1-2 gene segment orthologue are activated by Env trimer immunization. PLoS Pathog, 10, e1004337.

Nogal, B., Bianchi, M., Cottrell, C. A., Kirchdoerfer, R. N., Sewall, L. M., Turner, H. L., Zhao, F., Sok, D., Burton, D. R., Hangartner, L. & Ward, A. B. 2020a. Mapping Polyclonal Antibody Responses in Non-human Primates Vaccinated with HIV Env Trimer Subunit Vaccines. Cell Rep, 30, 3755–3765 e7.

Nogal, B., McCoy, L. E., van Gils, M. J., Cottrell, C. A., Voss, J. E., Andrabi, R., Pauthner, M., Liang, C. H., Messmer, T., Nedellec, R., Shin, M., Turner, H. L., Ozorowski, G., Sanders, R. W., Burton, D. R. & Ward, A. B. 2020b. HIV envelope trimer-elicited autologous neutralizing antibodies bind a region overlapping the N332 glycan supersite. Sci Adv, 6, eaba0512.

Ou, L., Kong, W. P., Chuang, G. Y., Ghosh, M., Gulla, K., O’Dell, S., Varriale, J., Barefoot, N., Changela, A., Chao, C. W., Cheng, C., Druz, A., Kong, R., Mckee, K., Rawi, R., Sarfo, E. K., Schon, A., Shaddeau, A., Tsybovsky, Y., Verardi, R., Wang, S., Wanninger, T. G., Xu, K., Yang, G. J., Zhang, B., Zhang, Y., Zhou, T., Program, V. R. C. P., Arnold, F. J., Doria-Rose, N. A., Lei, Q. P., Ryan, E. T., Vann, W. F., Mascola, J. R. & Kwong, P. D. 2020. Preclinical Development of a Fusion Peptide Conjugate as an HIV Vaccine Immunogen. Sci Rep, 10, 3032.

Ozorowski, G., Cupo, A., Golabek, M., Lopiccolo, M., Ketas, T. A., Cavallary, M., Cottrell, C. A., Klasse, P. J., Ward, A. B. & Moore, J. P. 2018. Effects of Adjuvants on HIV-1 Envelope Glycoprotein SOSIP Trimers In Vitro. J Virol, 92.

Pauthner, M., Havenar-Daughton, C., Sok, D., Nkolola, J. P., Bastidas, R., Boopathy, A. V., Carnathan, D. G., Chandrashekar, A., Cirelli, K. M., Cottrell, C. A., Eroshkin, A. M., Guenaga, J., Kaushik, K., Kulp, D. W., Liu, J., Mccoy, L. E., Oom, A. L., Ozorowski, G., Post, K. W., Sharma, S. K., Steichen, J. M., De TAEYE, S. W., Tokatlian, T., Torrents de la Pena, A., Butera, S. T., Labranche, C. C., Montefiori, D. C., Silvestri, G., Wilson, I. A., Irvine, D. J., Sanders, R. W., Schief, W. R., Ward, A. B., Wyatt, R. T., Barouch, D. H., Crotty, S. & Burton, D. R. 2017. Elicitation of Robust Tier 2 Neutralizing Antibody Responses in Nonhuman Primates by HIV Envelope Trimer Immunization Using Optimized Approaches. Immunity, 46, 1073–1088 e6.

Pauthner, M. G., Nkolola, J. P., Havenar-Daughton, C., Murrell, B., Reiss, S. M., Bastidas, R., Prevost, J., Nedellec, R., von Bredow, B., Abbink, P., Cottrell, C. A., Kulp, D. W., Tokatlian, T., Nogal, B., Bianchi, M., Li, H., Lee, J. H., Butera, S. T., Evans, D. T., Hangartner, L., Finzi, A., Wilson, I. A., Wyatt, R. T., Irvine, D. J., Schief, W.R., Ward, A. B., Sanders, R. W., Crotty, S., Shaw, G. M., Barouch, D. H. & Burton, D. R. 2019. Vaccine-Induced Protection from Homologous Tier 2 SHIV Challenge in Nonhuman Primates Depends on Serum-Neutralizing Antibody Titers. Immunity, 50, 241–252 e6.

Pettersen, E. F., Goddard, T. D., Huang, C. C., Couch, G. S., Greenblatt, D. M., Meng, E. C. & Ferrin, T. E. 2004. UCSF Chimera--a visualization system for exploratory research and analysis. J Comput Chem, 25, 1605–12.

Punjani, A., Rubinstein, J. L., Fleet, D. J. & Brubaker, M. A. 2017. cryoSPARC: algorithms for rapid unsupervised cryo-EM structure determination. Nat Methods, 14, 290–296.

Ringe, R. P., Cruz Portillo, V. M., Dosenovic, P., Ketas, T. J., Ozorowski, G., Nogal, B., Perez, L., Labranche, C. C., Lim, J., Francomano, E., Wilson, I. A., Sanders, R. W., Ward, A. B., Montefiori, D. C., Nussenzweig, M. C., Klasse, P. J., Cupo, A. & Moore, J. P. 2020. Neutralizing Antibody Induction by HIV-1 Envelope Glycoprotein SOSIP Trimers on Iron Oxide Nanoparticles May Be Impaired by Mannose Binding Lectin. J Virol, 94.

Ringe, R. P., Ozorowski, G., Rantalainen, K., Struwe, W. B., Matthews, K., Torres, J. L., Yasmeen, A., Cottrell, C. A., Ketas, T. J., Labranche, C. C., Montefiori, D. C., Cupo, A., Crispin, M., Wilson, I. A., Ward, A. B., Sanders, R. W., Klasse, P. J. & Moore, J. P. 2017. Reducing V3 Antigenicity and Immunogenicity on Soluble, Native-Like HIV-1 Env SOSIP Trimers. J Virol, 91.

Ringe, R. P., Pugach, P., Cottrell, C. A., Labranche, C. C., Seabright, G. E., Ketas, T. J., Ozorowski, G., Kumar, S., Schorcht, A., van Gils, M. J., Crispin, M., Montefiori, D.C., Wilson, I. A., Ward, A. B., Sanders, R. W., Klasse, P. J. & Moore, J. P. 2019. Closing and Opening Holes in the Glycan Shield of HIV-1 Envelope Glycoprotein SOSIP Trimers Can Redirect the Neutralizing Antibody Response to the Newly Unmasked Epitopes. J Virol, 93.

Sanders, R. W., Derking, R., Cupo, A., Julien, J. P., Yasmeen, A., De VAL, N., Kim, H. J., Blattner, C., de la Pena, A. T., Korzun, J., Golabek, M., de los Reyes, K., Ketas, T. J., van Gils, M. J., King, C. R., Wilson, I. A., Ward, A. B., Klasse, P. J. & Moore, J. P. 2013. A next-generation cleaved, soluble HIV-1 Env trimer, BG505 SOSIP.664 gp140, expresses multiple epitopes for broadly neutralizing but not non-neutralizing antibodies. PLoS Pathog, 9, e1003618.

Sanders, R. W., van Gils, M. J., Derking, R., Sok, D., Ketas, T. J., Burger, J. A., Ozorowski, G., Cupo, A., Simonich, C., Goo, L., Arendt, H., Kim, H. J., Lee, J. H., Pugach, P., Williams, M., Debnath, G., Moldt, B., van Breemen, M. J., Isik, G., Medina-Ramirez, M., Back, J. W., Koff, W. C., Julien, J. P., Rakasz, E. G., Seaman, M. S., Guttman, M., Lee, K. K., Klasse, P. J., Labranche, C., Schief, W. R., Wilson, I. A., Overbaugh, J., Burton, D. R., Ward, A. B., Montefiori, D. C., Dean, H. & Moore, J. P. 2015. HIV-1 VACCINES. HIV-1 neutralizing antibodies induced by native-like envelope trimers. Science, 349, aac4223.

Saunders, K. O., Wiehe, K., Tian, M., Acharya, P., Bradley, T., Alam, S. M., Go, E. P., Scearce, R., Sutherland, L., Henderson, R., Hsu, A. L., Borgnia, M. J., Chen, H., Lu, X., Wu, N. R., Watts, B., Jiang, C., Easterhoff, D., Cheng, H. L., McGovern, K., Waddicor, P., Chapdelaine-Williams, A., Eaton, A., Zhang, J., Rountree, W., Verkoczy, L., Tomai, M., Lewis, M. G., Desaire, H. R., Edwards, R. J., Cain, D. W., Bonsignori, M., Montefiori, D., Alt, F. W. & Haynes, B. F. 2019. Targeted selection of HIV-specific antibody mutations by engineering B cell maturation. Science, 366.

Sharma, S. K., de Val, N., Bale, S., Guenaga, J., Tran, K., Feng, Y., Dubrovskaya, V., Ward, A. B. & Wyatt, R. T. 2015. Cleavage-independent HIV-1 Env trimers engineered as soluble native spike mimetics for vaccine design. Cell Rep, 11, 539–50.

Steichen, J. M., Kulp, D. W., Tokatlian, T., Escolano, A., Dosenovic, P., Stanfield, R. L., Mccoy, L. E., Ozorowski, G., Hu, X., Kalyuzhniy, O., Briney, B., Schiffner, T., Garces, F., Freund, N. T., Gitlin, A. D., Menis, S., Georgeson, E., Kubitz, M., Adachi, Y., Jones, M., Mutafyan, A. A., Yun, D. S., Mayer, C. T., Ward, A. B., Burton, D. R., Wilson, I. A., Irvine, D. J., Nussenzweig, M. C. & Schief, W. R. 2016. HIV Vaccine Design to Target Germline Precursors of Glycan-Dependent Broadly Neutralizing Antibodies. Immunity, 45, 483–496.

Steichen, J. M., Lin, Y. C., Havenar-Daughton, C., Pecetta, S., Ozorowski, G., Willis, J. R., Toy, L., Sok, D., Liguori, A., Kratochvil, S., Torres, J. L., Kalyuzhniy, O., Melzi, E., Kulp, D. W., Raemisch, S., Hu, X., Bernard, S. M., Georgeson, E., Phelps, N., Adachi, Y., Kubitz, M., Landais, E., Umotoy, J., Robinson, A., Briney, B., Wilson, I. A., Burton, D. R., Ward, A. B., Crotty, S., Batista, F. D. & Schief, W. R. 2019. A generalized HIV vaccine design strategy for priming of broadly neutralizing antibody responses. Science, 366.

Suloway, C., Pulokas, J., Fellmann, D., Cheng, A., Guerra, F., Quispe, J., Stagg, S., Potter, C. S. & Carragher, B. 2005. Automated molecular microscopy: the new Leginon system. J Struct Biol, 151, 41–60.

Tokatlian, T., Kulp, D. W., Mutafyan, A. A., Jones, C. A., Menis, S., Georgeson, E., Kubitz, M., Zhang, M. H., Melo, M. B., Silva, M., Yun, D. S., Schief, W. R. & Irvine, D. J. 2018. Enhancing Humoral Responses Against HIV Envelope Trimers via Nanoparticle Delivery with Stabilized Synthetic Liposomes. Sci Rep, 8, 16527.

Torrents de la Pena, A., Julien, J. P., de Taeye, S. W., Garces, F., Guttman, M., Ozorowski, G., Pritchard, L. K., Behrens, A. J., Go, E. P., Burger, J. A., Schermer, E. E., Sliepen, K., Ketas, T. J., Pugach, P., Yasmeen, A., Cottrell, C. A., Torres, J. L., Vavourakis, C. D., van Gils, M. J., Labranche, C., Montefiori, D. C., Desaire, H., Crispin, M., Klasse, P. J., Lee, K. K., Moore, J. P., Ward, A. B., Wilson, I. A. & Sanders, R. W. 2017. Improving the Immunogenicity of Native-like HIV-1 Envelope Trimers by Hyperstabilization. Cell Rep, 20, 1805–1817.

Tran, K., Poulsen, C., Guenaga, J., De VAL, N., Wilson, R., Sundling, C., Li, Y., Stanfield, R. L., Wilson, I. A., Ward, A. B., Karlsson HEDESTAM, G. B. & Wyatt, R. T. 2014. Vaccine-elicited primate antibodies use a distinct approach to the HIV-1 primary receptor binding site informing vaccine redesign. Proc Natl Acad Sci U S A, 111, E738–47.

Travers, S. A. 2012. Conservation, Compensation, and Evolution of N-Linked Glycans in the HIV- 1 Group M Subtypes and Circulating Recombinant Forms. ISRN AIDS, 2012, 823605.

Ueda, G., Antanasijevic, A., Fallas, J. A., Sheffler, W., Copps, J., Ellis, D., Hutchinson, G. B., Moyer, A., Yasmeen, A., Tsybovsky, Y., Park, Y. J., Bick, M. J., Sankaran, B., Gillespie, R. A., Brouwer, P. J., Zwart, P. H., Veesler, D., Kanekiyo, M., Graham, B. S., Sanders, R. W., Moore, J. P., Klasse, P. J., Ward, A. B., King, N. P. & Baker, D. 2020. Tailored design of protein nanoparticle scaffolds for multivalent presentation of viral glycoprotein antigens. Elife, 9.

van Gils, M. J., van den Kerkhof, T. L., Ozorowski, G., Cottrell, C. A., Sok, D., Pauthner, M., Pallesen, J., De VAL, N., Yasmeen, A., De TAEYE, S. W., Schorcht, A., Gumbs, S., Johanna, I., Saye-Francisco, K., Liang, C. H., Landais, E., Nie, X., Pritchard, L. K., Crispin, M., Kelsoe, G., Wilson, I. A., Schuitemaker, H., Klasse, P. J., Moore, J. P., Burton, D. R., Ward, A. B. & Sanders, R. W. 2016. An HIV-1 antibody from an elite neutralizer implicates the fusion peptide as a site of vulnerability. Nat Microbiol, 2, 16199.

Wang, R. Y., Song, Y., Barad, B. A., Cheng, Y., Fraser, J. S. & Dimaio, F. 2016. Automated structure refinement of macromolecular assemblies from cryo-EM maps using Rosetta. Elife, 5.

Wang, W., Nie, J., Prochnow, C., Truong, C., Jia, Z., Wang, S., Chen, X. S. & Wang, Y. 2013. A systematic study of the N-glycosylation sites of HIV-1 envelope protein on infectivity and antibody-mediated neutralization. Retrovirology, 10, 14.

Whitaker, N., Hickey, J. M., Kaur, K., Xiong, J., Sawant, N., Cupo, A., Lee, W. H., Ozorowski, G., Medina-Ramirez, M., Ward, A. B., Sanders, R. W., Moore, J. P., Joshi, S. B., Volkin, D. B. & Dey, A. K. 2019. Developability Assessment of Physicochemical Properties and Stability Profiles of HIV-1 BG505 SOSIP.664 and BG505 SOSIP.v4.1-GT1.1 gp140 Envelope Glycoprotein Trimers as Candidate Vaccine Antigens. J Pharm Sci, 108, 2264–2277.

Williams, C. J., Headd, J. J., Moriarty, N. W., Prisant, M. G., Videau, L. L., Deis, L. N., Verma, V., Keedy, D. A., Hintze, B. J., Chen, V. B., Jain, S., Lewis, S. M., Arendall, W. B., 3RD, Snoeyink, J., Adams, P. D., Lovell, S. C., Richardson, J. S. & Richardson, D. C. 2018. MolProbity: More and better reference data for improved all-atom structure validation. Protein Sci, 27, 293–315.

Xu, K., Acharya, P., Kong, R., Cheng, C., Chuang, G. Y., Liu, K., Louder, M. K., O’Dell, S., Rawi, R., Sastry, M., Shen, C. H., Zhang, B., Zhou, T., Asokan, M., Bailer, R. T., Chambers, M., Chen, X., Choi, C. W., Dandey, V. P., Doria-Rose, N. A., Druz, A., Eng, E.T., Farney, S. K., Foulds, K. E., Geng, H., Georgiev, I. S., Gorman, J., Hill, K. R., Jafari, A. J., Kwon, Y. D., Lai, Y. T., Lemmin, T., Mckee, K., Ohr, T. Y., Ou, L., Peng, D., Rowshan, A. P., Sheng, Z., Todd, J. P., Tsybovsky, Y., Viox, E. G., Wang, Y., Wei, H., Yang, Y., Zhou, A. F., Chen, R., Yang, L., Scorpio, D. G., McDermott, A. B., Shapiro, L., Carragher, B., Potter, C. S., Mascola, J. R. & Kwong, P. D. 2018. Epitope-based vaccine design yields fusion peptide-directed antibodies that neutralize diverse strains of HIV-1. Nat Med, 24, 857–867.

Yang, Y. R., Mccoy, L. E., van Gils, M. J., Andrabi, R., Turner, H. L., Yuan, M., Cottrell, C. A., Ozorowski, G., Voss, J., Pauthner, M., Polveroni, T. M., Messmer, T., Wilson, I. A., Sanders, R. W., Burton, D. R. & Ward, A. B. 2020. Autologous Antibody Responses to an HIV Envelope Glycan Hole Are Not Easily Broadened in Rabbits. J Virol, 94.

Yuan, M., Cottrell, C. A., Ozorowski, G., van Gils, M. J., Kumar, S., Wu, N. C., Sarkar, A., Torres, J. L., de Val, N., Copps, J., Moore, J. P., Sanders, R. W., Ward, A. B. & Wilson, I. A. 2019. Conformational Plasticity in the HIV-1 Fusion Peptide Facilitates Recognition by Broadly Neutralizing Antibodies. Cell Host Microbe, 25, 873–883 e5.

Zhang, K. 2016. Gctf: Real-time CTF determination and correction. J Struct Biol, 193, 1–12.

Zhao, F., Joyce, C., Burns, A., Nogal, B., Cottrell, C. A., Ramos, A., Biddle, T., Pauthner, M., Nedellec, R., Qureshi, H., Mason, R., Landais, E., Briney, B., Ward, A. B., Burton, D. R. & Sok, D. 2020. Mapping Neutralizing Antibody Epitope Specificities to an HIV Env Trimer in Immunized and in Infected Rhesus Macaques. Cell Rep, 32, 108122.

Zheng, S. Q., Palovcak, E., Armache, J. P., Verba, K. A., Cheng, Y. & Agard, D. A. 2017. MotionCor2: anisotropic correction of beam-induced motion for improved cryo-electron microscopy. Nat Methods, 14, 331–332.

Zivanov, J., Nakane, T., Forsberg, B. O., Kimanius, D., Hagen, W. J., Lindahl, E. & Scheres, S. H. 2018. New tools for automated high-resolution cryo-EM structure determination in RELION-3. Elife, 7.

