## Supplemental Information for "Polyclonal antibody responses to HIV Env immunogens resolved using cryoEM"

### Characterization of Polyclonal Antibody Responses to HIV Env Immunogens Using CryoEMPEM

#### Supplementary Information

**Table S1. Stabilizing and glycan knock-in mutations in BG505 SOSIP constructs used in this study (green – mutation present; red – mutation not present).**

| | BG505 SOSIP | | | |
|---|---|---|---|---|
| **Mutations** | **v3** | **MD39** | **v5.2 N241/N289** | **v5.2(7S) N241/N289** |
| **A501C** | green | green | green | green |
| **T605C** | green | green | green | green |
| **I559P** | green | green | green | green |
| **E64K** | red | red | green | green |
| **A73C** | red | red | green | green |
| **A316W** | red | red | green | green |
| **A561C** | red | red | green | green |
| **M271I** | red | green | green | green |
| **A319Y** | red | green | red | green |
| **R585H** | red | green | red | green |
| **L568D** | red | green | red | green |
| **V570H** | red | green | red | green |
| **R304V** | red | green | red | green |
| **F519S** | red | green | red | green |
| **T106E** | red | green | red | red |
| **A561P** | red | green | red | red |
| **N363Q** | red | green | red | red |
| **P240T** | red | red | green | green |
| **S241N** | red | red | green | green |
| **F288L** | red | green | green | green |
| **T290E** | red | red | green | green |
| **P291S** | red | red | green | green |
| **Q658K** | red | red | red | red |
| **S613T** | red | red | green | red |

**Table S2. Anti-trimer binding antibody titers (midpoint, $EC_{50}$) determined via ELISA for plasma samples collected at indicated time points**. Color coding: white - value not determined (N/D) or material not available (N/A); light-gray - no detectable binding ($EC_{50}$ < 20); light orange - 20 < $EC_{50}$ < 100); medium orange - 100 < $EC_{50}$ < 500; dark orange - 500 < $EC_{50}$ < 1000; brown - $EC_{50}$ > 1000.

| **Group / Immunogen** | Animal ID | **Week 6** | **Week 10** | **Week 26** | **Week 38** |
|---|---|---|---|---|---|
| **Grp 1** BG505 SOSIP MD39 | 32034 | < 20 | 93 | 167 | 164 |
| | 32113 | < 20 | 196 | 479 | 1233 |
| | 34943 | < 20 | 75 | 166 | 101 |
| | 34909 | < 20 | 115 | 109 | N/D |
| | 33104 | < 20 | 151 | 323 | 231 |
| | 33395 | < 20 | 288 | 516 | 253 |
| **Grp 2** BG505 SOSIP.v5.2 N241/N289 | 34686 | 61 | 277 | 134 | 86 |
| | 33182 | 36 | 1073 | 587 | 364 |
| | 33311 | 35 | 1921 | 121 | 626 |
| | CD99 | 54 | 595 | 242 | 325 |
| | 33203 | 47 | 381 | 310 | 233 |
| | CG41 | 42 | 1430 | 256 | 378 |
| **Grp 3** BG505 SOSIP.v5.2(7S) N241/N289 T33-31 NP | 33172 | 33 | 858 | 431 | 595 |
| | 34919 | < 20 | 342 | 384 | 356 |
| | 34167 | 64 | 929 | 860 | 832 |
| | 33065 | 30 | 211 | 178 | 295 |
| | 33176[#] | 54 | 451 | N/A | N/A |
| | 34725 | 57 | 480 | 397 | 850 |

### This animal was euthanized early due to chronic health issues. Sera samples were not available for Week 26 and Week 38 time points (see Methods section for details).

N/D – Not determined

N/A – Samples not available

**Table S3. Autologous neutralization titers ($ID_{50}$) against BG505-based pseudovirus.** Midpoint neutralization titers ($ID_{50}$) were determined using sera samples collected at different time points (indicated in the top row). Neutralization assays with VRC01 were also performed (positive control). Color coding: light gray - no neutralization ($ID_{50} < 40$); light orange - very weak neutralization ($40 < ID_{50} < 100$); medium orange - moderate neutralization ($100 < ID_{50} < 500$); dark orange - strong neutralization ($500 < ID_{50} < 2000$); brown - very strong neutralization ($ID_{50} > 2000$).

| | | Week 10 | Week 26 | Week 38 |
|---|---|---|---|---|
| | Virus | BG505 | BG505 | BG505 |
| Group / Immunogen | Animal ID | | | |
| **Grp 1** BG505 SOSIP MD39 | 32034 | 41 | 322 | 1131 |
| | 32113 | < 40 | < 40 | < 40 |
| | 34943 | < 40 | < 40 | < 40 |
| | 34909 | 84 | 506 | 1212 |
| | 33104 | < 40 | < 40 | < 40 |
| | 33395 | 42 | 267 | 1477 |
| **Grp 2** BG505 SOSIP.v5.2 N241/N289 | 34686 | < 40 | < 40 | < 40 |
| | 33182 | < 40 | < 40 | < 40 |
| | 33311 | < 40 | 604 | 8599 |
| | CD99 | < 40 | < 40 | 111 |
| | 33203 | < 40 | 66 | 123 |
| | CG41 | < 40 | 378 | 507 |
| **Grp 3** BG505 SOSIP.v5.2(7S) N241/N289 T33-31 NP | 33172 | < 40 | < 40 | 407 |
| | 34919 | < 40 | < 40 | < 40 |
| | 34167 | < 40 | < 40 | < 40 |
| | 33065 | < 40 | 47 | 232 |
| | 33176# | < 40 | N/A | N/A |
| | 34725 | < 40 | < 40 | < 40 |
| VRC01 | $IC_{50}$ (µg/ml) | 0.05 | 0.04 | 0.05 |

### This animal was euthanized early in the study due to chronic health issues. Sera samples were not available for Week 26 and Week 38 time points (see Methods section for details)

N/A – Samples not available

**Table S4. Cryo-EM data collection information**

| | Rh.32034 Poly. Fab + BG505 SOSIP MD39 | Rh.33104 Poly. Fab + BG505 SOSIP MD39 | Rh.33311 Poly. Fab + BG505 SOSIP v5.2 N241/N289 | Rh.33172 Poly. Fab + BG505 SOSIP v5.2(7S) N241/N289 | BG505 SOSIP T33-31-NP |
|---|---|---|---|---|---|
| **Microscope** | Titan Krios | Titan Krios | Titan Krios | Titan Krios | Talos Arctica |
| **Voltage (kV)** | 300 | 300 | 300 | 300 | 200 |
| **Detector** | Gatan K2 Summit | Gatan K2 Summit | Gatan K2 Summit | Gatan K2 Summit | Gatan K2 Summit |
| **Recording mode** | Counting | Counting | Counting | Counting | Counting |
| **Magnification** | 29,000 | 29,000 | 29,000 | 29,000 | 36,000 |
| **Movie micrograph pixel size** | 1.03 | 1.03 | 1.03 | 1.03 | 1.15 |
| **Dose rate ($e^-$/Å$^2$/s)** | 5.98 | 4.55 | 4.55 | 5.63 | 4.88 |
| **No. of frames per movie micrograph** | 30 | 39 | 39 | 36 | 41 |
| **Frame exposure time (ms)** | 250 | 250 | 250 | 250 | 250 |
| **Movie micrograph exposure time (s)** | 7.50 | 9.75 | 9.75 | 9.00 | 10.25 |
| **Total dose ($e^-$/Å$^2$)** | 44.9 | 44.3 | 44.3 | 50.7 | 50.0 |
| **Grid Type** | UltrAuFoil R 1.2/1.3 | UltrAuFoil R 1.2/1.3 | UltrAuFoil R 1.2/1.3 | Quantifoil R 2/1 | Quantifoil R 2/1 |
| **Under focus range (μm)** | 0.6 – 1.6 | 0.6 – 1.6 | 0.6 – 1.6 | 0.6 – 1.6 | 0.8 – 2.0 |
| **Number of movie micrographs** | 3,194 | 3,580 | 6,268 | 4,521 | 1,751 |

#### Table S5. Map and model refinement information

| Animal ID | Rh.32034 (Grp 1) | | | | Rh.33104 (Grp 1) | | | | Rh.33311 (Grp 2) | | | | | | | |
|---|---|---|---|---|---|---|---|---|---|---|---|---|---|---|---|---|
| **Complex ID** | **pAbC-1** | **pAbC-2** | **pAbC-3** | **pAbC-4** | **pAbC-1** | **pAbC-2** | **pAbC-3** | **pAbC-4** | **pAbC-1** | **pAbC-2** | **pAbC-3** | **pAbC-4** | **pAbC-5** | **pAbC-6** | **pAbC-7** | **pAbC-8** |
| **Antigen** | **BG505 SOSIP MD39** | | | | **BG505 SOSIP MD39** | | | | **BG505 SOSIP.v5.2 N241/N289** | | | | | | | |
| **Number of picked particles** | 591,593 | | | | 502,068 | | | | 1,267,693 | | | | | | | |
| **Particles after 2D classification** | 327,148 | | | | 230,034 | | | | 503,502 | | | | | | | |
| **Particles after symmetry expansion** | 981,444 | | | | 690,102 | | | | 1,510,506 | | | | | | | |
| **Particles in the final map** | 48,919 | 37,870 | 47,950 | 106,016 | 81,561 | 27,719 | 58,378 | 11,534 | 82,126 | 28,188 | 39,028 | 27,483 | 52,082 | 12,653 | 70,307 | 29,622 |
| **Map symmetry** | C1 | C1 | C1 | C1 | C1 | C1 | C1 | C1 | C1 | C1 | C1 | C1 | C1 | C1 | C1 | C1 |
| **Map sharpening B-factor** | -79.6 | -94.3 | -7.1 | -12.1 | -87.6 | -90.8 | -83.4 | -93.9 | -103.8 | -118.7 | -122.1 | -103.6 | -126.5 | -131.0 | -121.8 | -117.2 |
| **Map Resolution** | 3.4 | 3.6 | 3.6 | 3.8 | 3.3 | 3.7 | 3.4 | 4.6 | 3.7 | 4.5 | 4.1 | 4 | 4.2 | 6.6 | 3.8 | 4.5 |
| **EMDB ID** | 23223 | 23224 | 23225 | 23226 | 23227 | 23228 | 23229 | 23230 | 23236 | 23237 | 23238 | 23239 | 23240 | 23241 | 23242 | 23243 |
| **Residues** | 2010 | 2004 | 1992 | 1988 | 2007 | 2003 | 2022 | 1972 | 2082 | 1978 | 2012 | 2039 | 2031 | N/A | 2026 | 1976 |
| **Amino acids** | 1910 | 1901 | 1898 | 1888 | 1910 | 1909 | 1911 | 1871 | 1994 | 1893 | 1926 | 1936 | 1951 | N/A | 1930 | 1897 |
| **Carbohydrates** | 100 | 103 | 94 | 100 | 97 | 94 | 111 | 101 | 88 | 85 | 86 | 103 | 80 | N/A | 96 | 79 |
| **RMSD Bonds (4σ)** | 0.022 | 0.021 | 0.021 | 0.022 | 0.023 | 0.021 | 0.025 | 0.021 | 0.021 | 0.021 | 0.02 | 0.021 | 0.02 | N/A | 0.022 | 0.02 |
| **RMSD Angles (4σ)** | 1.653 | 1.722 | 1.684 | 1.723 | 1.709 | 1.687 | 1.698 | 1.669 | 1.674 | 1.702 | 1.679 | 1.647 | 1.633 | N/A | 1.716 | 1.71 |
| **Ramachandran** | | | | | | | | | | | | | | | | |
| **Outliers (%)** | 0 | 0 | 0 | 0 | 0 | 0 | 0 | 0 | 0 | 0 | 0 | 0 | 0 | N/A | 0 | 0 |
| **Allowed (%)** | 1.88 | 3.5 | 3.19 | 2.61 | 2.09 | 2.57 | 2.41 | 2.25 | 2.19 | 3.35 | 2.01 | 2.53 | 2.66 | N/A | 3.59 | 2.86 |
| **Favored (%)** | 98.12 | 96.5 | 96.81 | 97.39 | 97.91 | 97.43 | 97.59 | 97.75 | 97.81 | 96.65 | 97.99 | 97.47 | 97.34 | N/A | 96.41 | 97.14 |
| **Rotamer outliers (%)** | 0 | 0 | 0 | 0 | 0 | 0 | 0 | 0 | 0 | 0 | 0 | 0 | 0 | N/A | 0 | 0 |
| **Clash score** | 1.22 | 1.99 | 2.17 | 2.07 | 1.19 | 2.16 | 0.9 | 2.6 | 1.45 | 2.67 | 1.34 | 0.91 | 1.87 | N/A | 1.01 | 2.97 |
| **Molprobity score** | 0.84 | 1.2 | 1.19 | 1.1 | 0.85 | 1.1 | 1.11 | 1.1 | 0.93 | 1.27 | 0.87 | 0.88 | 1.08 | N/A | 1.04 | 1.24 |
| **EMRinger score** | 4.17 | 3.36 | 3.75 | 3.71 | 4.33 | 3.73 | 4.33 | 0.77 | 3.25 | 0.93 | 2.29 | 2.79 | 2.39 | N/A | 2.99 | 1.38 |
| **PDB ID** | 7L86 | 7L87 | 7L88 | 7L89 | 7L8A | 7L8B | 7L8C | 7L8D | 7L8T | 7L8U | 7L8W | 7L8X | 7L8Y | N/A | 7L8Z | 7L90 |

#### Table S5. Map and model refinement information (continued)

| **Animal ID** | **N/A** | | | **Rh.33172 (Grp 3)** | | | | |
|---|---|---|---|---|---|---|---|---|
| **Complex ID** | **T33-31 NP core** | **BG505 SOSIP Comp A** | **BG505 SOSIP Comp B** | **pAbC-1** | **pAbC-2** | **pAbC-3** | **pAbC-4** | **pAbC-5** |
| **Antigen** | N/A | | | **BG505 SOSIP.v5.2(7S) N241/N289** | | | | |
| **Number of picked particles** | 223,099 | 441,476* | 441,476* | 1,121,581 | | | | |
| **Particles after 2D classification** | 214,224 | 353,064 | 316,582 | 456,268 | | | | |
| **Particles after symmetry expansion** | N/A | | | 1,368,804 | | | | |
| **Particles in the final map** | 110,369 | 106,478 | 64,726 | 31,914 | 66,171 | 33,361 | 28,929 | 33,802 |
| **Map symmetry** | T | C3 | C3 | C1 | C1 | C1 | C1 | C1 |
| **Map sharpening B-factor** | -53.1 | -70.5 | -164.5 | -124.0 | -105.9 | -95.1 | -112.4 | -120.8 |
| **Map Resolution** | 2.9 | 3.7 | 3.8 | 4.2 | 3.7 | 4.1 | 4.3 | 4.0 |
| **EMDB ID** | 23222 | 23218 | 23219 | 23231 | 23232 | 23233 | 23235 | 23234 |
| **Residues** | 2664 | 1872 | 1857 | 2007 | 2027 | 2002 | 2029 | N/A |
| **Amino acids** | 2664 | 1770 | 1761 | 1912 | 1939 | 1904 | 1925 | N/A |
| **Carbohydrates** | 0 | 102 | 96 | 95 | 88 | 98 | 104 | N/A |
| **RMSD Bonds (4σ)** | 0.025 | 0.023 | 0.022 | 0.021 | 0.021 | 0.021 | 0.021 | N/A |
| **RMSD Angles (4σ)** | 1.653 | 1.701 | 1.645 | 1.691 | 1.654 | 1.701 | 1.687 | N/A |
| **Ramachandran** | | | | | | | | |
| **Outliers (%)** | 0 | 0 | 0 | 0 | 0 | 0 | 0 | N/A |
| **Allowed (%)** | 2.29 | 1.21 | 1.39 | 1.76 | 2 | 2.15 | 1.91 | N/A |
| **Favored (%)** | 97.71 | 98.79 | 98.61 | 98.24 | 98 | 97.85 | 98.09 | N/A |
| **Rotamer outliers (%)** | 0 | 0 | 0 | 0 | 0 | 0 | 0 | N/A |
| **Clash score** | 2.46 | 0.2 | 1.76 | 2.15 | 1.34 | 1.06 | 1.48 | N/A |
| **Molprobity score** | 1.1 | 0.58 | 0.93 | 0.99 | 0.86 | 0.84 | 0.89 | N/A |
| **EMRinger score** | 4.84 | 2.9 | 2.40 | 2.19 | 3.31 | 1.87 | 1.9 | N/A |
| **PDB ID** | 7L85 | 7L7T | 7L7U | 7L8E | 7L8F | 7L8G | 7L8S | N/A |

**Table S6. Neutralization titers ($ID_{50}$) against mutant BG505 pseudoviruses.** Midpoint neutralization titers ($ID_{50}$) were determined using sera samples collected at the week 38 time point. Neutralization assays with MLV (negative control) and CH01-31 (positive control and reference) were performed. Color coding: light gray - no neutralization ($ID_{50} < 20$); light orange - very weak neutralization ($20 < ID_{50} < 100$); medium orange - moderate neutralization ($100 < ID_{50} < 500$); dark orange - strong neutralization ($500 < ID_{50} < 2000$); brown - very strong neutralization ($ID_{50} > 2000$).

| | | **Week 38** | | | | | | |
|---|---|---|---|---|---|---|---|---|
| | Virus | SVA **MLV** | BG505 **T332N** | BG505 **T332N.T465N** | BG505 **T332N.S241N** | BG505 **T332N.P291T** | BG505 **T332N.N156A** | BG505 **T332N.N611A** |
| Group / Immunogen | Animal ID | | | | | | | |
| **Grp 1** BG505 SOSIP MD39 | 32034 | <20 | 714 | <20 | 870 | 413 | 417 | N/D |
| | 32113 | <20 | <20 | <20 | <20 | <20 | <20 | N/D |
| | 34943 | <20 | 36 | <20 | <20 | <20 | <20 | N/D |
| | 34909 | <20 | 285 | <20 | 314 | 126 | <20 | N/D |
| | 33104 | <20 | <20 | <20 | <20 | <20 | <20 | N/D |
| | 33395 | <20 | 1129 | 338 | 1137 | 749 | 627 | N/D |
| **Grp 2** BG505 SOSIP.v5.2 N241/N289 | 34686 | <20 | <20 | <20 | N/D | <20 | N/D | 25 |
| | 33182 | <20 | <20 | <20 | N/D | <20 | N/D | 72 |
| | 33311 | <20 | 3073 | 200 | N/D | 1383 | N/D | 4237 |
| | CD99 | <20 | 67 | 41 | N/D | 34 | N/D | 60 |
| | 33203 | <20 | 27 | <20 | N/D | <20 | N/D | 41 |
| | CG41 | <20 | 1608 | <20 | N/D | 759 | N/D | 1595 |
| **Grp 3** BG505 SOSIP.v5.2(7S) N241/N289 T33-31 NP | 33172 | <20 | 229 | 230 | 207 | 177 | 320 | N/D |
| | 34919 | <20 | <20 | <20 | <20 | <20 | <20 | N/D |
| | 34167 | <20 | <20 | <20 | <20 | <20 | 107 | N/D |
| | 33065 | <20 | 171 | 142 | 172 | 110 | 299 | N/D |
| | 33176# | N/A | N/A | N/A | N/A | N/A | N/A | N/A |
| | 34725 | <20 | <20 | <20 | <20 | <20 | 370 | N/D |
| CH01-31 | $IC_{50}$ (μg/ml) | >25 | 0.03 | 0.028 | 0.028 | 0.013 | 0.036 | 0.019 |

#This animal was euthanized early in the study due to chronic health issues. Sera samples were not available for Week 26 and Week 38 time points (see Methods section for details)

N/D – Not determined

N/A – Samples not available

**Table S7. Site-specific glycosylation analysis of BG505-SOSIP-presenting nanoparticle components and free BG505-SOSIPv5.2(7S) N241/N289.** Glycan compositions corresponding to oligomannose/hybrid-type (green) and fully processed complex type (magenta) are shown at each potential N-linked glycosylation site (PNGS). Unglycosylated PNGS are shown in grey. Oligomannose-type glycans are divided with respect to the number of mannose residues present while hybrid-type glycans are categorized according to the presence/absence of fucose. For complex-type glycans, the categorization was performed according to the number of processed antenna and the presence of fucose. For PNGS where only low intensity peptides were present, the categorization was not performed and so in the table they are merged to cover all possible oligomannose/hybrid compositions or complex-type glycans. The unliganded trimer data is adapted from a manuscript by Antanasijevic et al., 2020; and displayed here as a reference for comparison.

**BG505 SOSIP.v5.2(7S) N241/N289**

| | N88 | N133 | N137 | N156 | N160 | N185e | N185h | N197 | N234 | N241 | N262 | N276 | N289 | N295 | N301 | N332 | N339 | N355 | N363 | N386 | N392 | N398 | N406 | N411 | N448 | N462 | N611 | N618 | N625 | N637 |
|---|---|---|---|---|---|---|---|---|---|---|---|---|---|---|---|---|---|---|---|---|---|---|---|---|---|---|---|---|---|---|
| High Mannose | 21 | 60 | 40 | 88 | 100 | 6 | 0 | 66 | 84 | 85 | 100 | 67 | 81 | 98 | 81 | 100 | 64 | 22 | 100 | n.d. | n.d. | n.d. | 0 | 100 | 100 | 1 | 1 | 0 | 24 | 34 |
| M9 | 0 | | | | 1 | 0 | | 10 | | | | 0 | | | | 29 | 37 | 0 | 63 | | | | | | 50 | 0 | | 0 | | 0 |
| M8 | 0 | | | | 21 | 2 | | 11 | | | | 5 | | | | 26 | 21 | 1 | 28 | | | | | | 31 | 0 | | 0 | | 0 |
| M7 | 0 | | | | 26 | 1 | | 14 | | | | 5 | | | | 31 | 4 | 1 | 5 | | | | | | 7 | 0 | | 0 | | 9 |
| M6 | 6 | | | | 21 | 0 | | 6 | | | | 0 | | | | 2 | 0 | 2 | 2 | | | | | | 3 | 0 | | 0 | | 0 |
| M5 | 13 | | | | 24 | 3 | | 17 | | | | 6 | | | | 10 | 2 | 8 | 1 | | | | | | 6 | 0 | | 0 | | 6 |
| M4 | 0 | | | | 7 | 0 | | 0 | | | | 2 | | | | 1 | 1 | 2 | 0 | | | | | | 2 | 0 | | 0 | | 0 |
| M3 | 0 | | | | 0 | 0 | | 0 | | | | 0 | | | | 0 | 0 | 1 | 0 | | | | | | 0 | 0 | | 0 | | 0 |
| Hybrid | 2 | | | | 0 | 0 | | 7 | | | | 37 | | | | 0 | 0 | 2 | 0 | | | | | | 0 | 0 | | 0 | | 8 |
| FHybrid | 0 | | | | 0 | 0 | | 0 | | | | 12 | | | | 0 | 0 | 6 | 0 | | | | | | 0 | 0 | | 0 | | 11 |
| A1 | 10 | | | | 0 | 0 | | 0 | | | | 0 | | | | 0 | 0 | 1 | 0 | | | | | | 0 | 0 | | 0 | | 0 |
| FA1 | 0 | | | | 0 | 2 | | 0 | | | | 2 | | | | 0 | 0 | 11 | 0 | | | | | | 0 | 2 | | 0 | | 0 |
| A2/A1B | 28 | | | | 0 | 0 | | 0 | | | | 13 | | | | 0 | 0 | 0 | 0 | | | | | | 0 | 0 | | 0 | | 0 |
| FA2/FA1B | 11 | 11 | 15 | 1 | 0 | 58 | 100 | 27 | 1 | 2 | 0 | 9 | 0 | 2 | 0 | 0 | 0 | 32 | 0 | n.d. | n.d. | n.d. | 100 | 0 | 0 | 45 | 50 | 26 | 1 | 13 |
| A3/A2B | 13 | | | | 0 | 0 | | 0 | | | | 4 | | | | 0 | 0 | 0 | 0 | | | | | | 0 | 0 | | 0 | | 0 |
| FA3/FA2B | 17 | | | | 0 | 15 | | 0 | | | | 5 | | | | 0 | 0 | 29 | 0 | | | | | | 0 | 50 | | 65 | | 0 |
| A4/A3B | 0 | | | | 0 | 0 | | 0 | | | | 0 | | | | 0 | 0 | 0 | 0 | | | | | | 0 | 0 | | 0 | | 0 |
| FA4/FA3B | 0 | | | | 0 | 0 | | 0 | | | | 0 | | | | 0 | 0 | 1 | 0 | | | | | | 0 | 2 | | 9 | | 0 |
| Unoccupied | 0 | 30 | 44 | 11 | 0 | 19 | 0 | 0 | 15 | 13 | 0 | 0 | 19 | 0 | 19 | 0 | 36 | 0 | 0 | n.d. | n.d. | n.d. | 0 | 0 | 0 | 0 | 49 | 0 | 75 | 44 |

**BG505 SOSIP-T33-31A**

| | N88 | N133 | N137 | N156 | N160 | N185e | N185h | N197 | N234 | N241 | N262 | N276 | N289 | N295 | N301 | N332 | N339 | N355 | N363 | N386 | N392 | N398 | N406 | N411 | N448 | N462 | N611 | N618 | N625 | N637 |
|---|---|---|---|---|---|---|---|---|---|---|---|---|---|---|---|---|---|---|---|---|---|---|---|---|---|---|---|---|---|---|
| High Mannose | 87 | 87 | 57 | 100 | 100 | 18 | 3 | 100 | 100 | 99 | 100 | 98 | 73 | 97 | 93 | 100 | 100 | 85 | 100 | 100 | 100 | 74 | 64 | 94 | 100 | 6 | 0 | 77 | 71 | 26 |
| M9 | 0 | | | | 11 | 0 | 0 | 54 | | | | 0 | | | | 81 | 64 | 0 | 62 | | | | | | 69 | 0 | 0 | | | 2 |
| M8 | 0 | | | | 36 | 1 | 0 | 32 | | | | 41 | | | | 16 | 25 | 11 | 32 | | | | | | 22 | 0 | 0 | | | 8 |
| M7 | 6 | | | | 23 | 2 | 0 | 15 | | | | 25 | | | | 2 | 6 | 17 | 4 | | | | | | 5 | 1 | 0 | | | 5 |
| M6 | 16 | | | | 17 | 3 | 0 | 0 | | | | 8 | | | | 0 | 3 | 11 | 2 | | | | | | 2 | 1 | 0 | | | 2 |
| M5 | 56 | | | | 14 | 10 | 3 | 0 | | | | 6 | | | | 0 | 2 | 37 | 0 | | | | | | 1 | 3 | 0 | | | 0 |
| M4 | 4 | | | | 0 | 0 | 0 | 0 | | | | 2 | | | | 0 | 0 | 2 | 0 | | | | | | 0 | 0 | 0 | | | 0 |
| M3 | 0 | | | | 0 | 0 | 0 | 0 | | | | 0 | | | | 0 | 0 | 0 | 0 | | | | | | 0 | 0 | 0 | | | 0 |
| Hybrid | 5 | | | | 0 | 1 | 0 | 0 | | | | 15 | | | | 0 | 0 | 4 | 0 | | | | | | 0 | 0 | 0 | | | 4 |
| FHybrid | 0 | | | | 0 | 0 | 0 | 0 | | | | 0 | | | | 0 | 0 | 3 | 0 | | | | | | 0 | 1 | 0 | | | 4 |
| A1 | 8 | | | | 0 | 1 | 0 | 0 | | | | 0 | | | | 0 | 0 | 1 | 0 | | | | | | 0 | 3 | 0 | | | 0 |
| FA1 | 0 | | | | 0 | 3 | 2 | 0 | | | | 0 | | | | 0 | 0 | 3 | 0 | | | | | | 0 | 4 | 0 | | | 1 |
| A2/A1B | 4 | | | | 0 | 0 | 0 | 0 | | | | 1 | | | | 0 | 0 | 0 | 0 | | | | | | 0 | 2 | 0 | | | 0 |
| FA2/FA1B | 0 | 0 | 11 | 0 | 0 | 36 | 77 | 0 | 0 | 0 | 0 | 1 | 0 | 0 | 0 | 0 | 0 | 4 | 0 | 0 | 0 | 26 | 36 | 6 | 0 | 53 | 0 | 21 | 29 | 45 |
| A3/A2B | 1 | | | | 0 | 0 | 0 | 0 | | | | 0 | | | | 0 | 0 | 0 | 0 | | | | | | 0 | 1 | 0 | | | 0 |
| FA3/FA2B | 0 | | | | 0 | 15 | 18 | 0 | | | | 0 | | | | 0 | 0 | 4 | 0 | | | | | | 0 | 28 | 90 | | | 27 |
| A4/A3B | 0 | | | | 0 | 0 | 0 | 0 | | | | 0 | | | | 0 | 0 | 0 | 0 | | | | | | 0 | 0 | 0 | | | 0 |
| FA4/FA3B | 0 | | | | 0 | 0 | 0 | 0 | | | | 0 | | | | 0 | 0 | 0 | 0 | | | | | | 0 | 2 | 10 | | | 0 |
| Unoccupied | 0 | 13 | 32 | 0 | 0 | 26 | 0 | 0 | 0 | 1 | 0 | 0 | 26 | 3 | 7 | 0 | 0 | 0 | 0 | 0 | 0 | 0 | 0 | 0 | 0 | 0 | 0 | 2 | 0 | 0 |

**BG505 SOSIP-T33-31B**

| | N88 | N133 | N137 | N156 | N160 | N185e | N185h | N197 | N234 | N241 | N262 | N276 | N289 | N295 | N301 | N332 | N339 | N355 | N363 | N386 | N392 | N398 | N406 | N411 | N448 | N462 | N611 | N618 | N625 | N637 |
|---|---|---|---|---|---|---|---|---|---|---|---|---|---|---|---|---|---|---|---|---|---|---|---|---|---|---|---|---|---|---|
| High Mannose | 55 | 92 | 54 | 100 | 100 | 9 | 3 | 60 | 100 | 100 | 100 | 89 | 69 | 91 | 79 | 100 | 76 | 79 | 100 | 100 | 100 | 69 | 57 | 98 | 100 | 3 | 26 | 31 | 78 | 57 |
| M9 | 0 | | | | 3 | 0 | 0 | 8 | | | | 0 | | | | 66 | 58 | 0 | 74 | 100 | | | | | 57 | 0 | | | | 0 |
| M8 | 2 | | | | 0 | 1 | 0 | 8 | | | | 21 | | | | 25 | 17 | 7 | 21 | 0 | | | | | 29 | 0 | | | | 6 |
| M7 | 4 | | | | 29 | 1 | 0 | 6 | | | | 19 | | | | 4 | 0 | 11 | 4 | 0 | | | | | 7 | 0 | | | | 12 |
| M6 | 16 | | | | 28 | 1 | 0 | 5 | | | | 10 | | | | 2 | 1 | 11 | 1 | 0 | | | | | 4 | 0 | | | | 8 |
| M5 | 18 | | | | 37 | 5 | 3 | 13 | | | | 9 | | | | 3 | 0 | 36 | 0 | 0 | | | | | 2 | 2 | | | | 5 |
| M4 | 2 | | | | 3 | 0 | 0 | 2 | | | | 4 | | | | 1 | 0 | 2 | 0 | 0 | | | | | 1 | 0 | | | | 0 |
| M3 | 5 | | | | 0 | 0 | 0 | 0 | | | | 1 | | | | 0 | 0 | 0 | 0 | 0 | | | | | 0 | 0 | | | | 0 |
| Hybrid | 7 | | | | 0 | 0 | 0 | 11 | | | | 24 | | | | 0 | 0 | 6 | 0 | 0 | | | | | 0 | 0 | | | | 13 |
| FHybrid | 1 | | | | 0 | 0 | 0 | 6 | | | | 0 | | | | 0 | 0 | 6 | 0 | 0 | | | | | 0 | 0 | | | | 13 |
| A1 | 8 | | | | 0 | 1 | 0 | 1 | | | | 3 | | | | 0 | 0 | 1 | 0 | 0 | | | | | 0 | 1 | | | | 1 |
| FA1 | 1 | | | | 0 | 4 | 2 | 7 | | | | 1 | | | | 0 | 0 | 6 | 0 | 0 | | | | | 0 | 4 | | | | 13 |
| A2/A1B | 29 | | | | 0 | 0 | 0 | 1 | | | | 5 | | | | 0 | 0 | 0 | 0 | 0 | | | | | 0 | 14 | | | | 0 |
| FA2/FA1B | 0 | 0 | 8 | 0 | 0 | 46 | 83 | 26 | 0 | 0 | 0 | 1 | 0 | 2 | 1 | 0 | 0 | 7 | 0 | 0 | 0 | 29 | 42 | 0 | 0 | 50 | 66 | 68 | 0 | 29 |
| A3/A2B | 0 | | | | 0 | 0 | 0 | 0 | | | | 2 | | | | 0 | 0 | 0 | 0 | 0 | | | | | 0 | 3 | | | | 0 |
| FA3/FA2B | 1 | | | | 0 | 17 | 12 | 5 | | | | 0 | | | | 0 | 0 | 5 | 0 | 0 | | | | | 0 | 24 | | | | 0 |
| A4/A3B | 0 | | | | 0 | 0 | 0 | 0 | | | | 0 | | | | 0 | 0 | 0 | 0 | 0 | | | | | 0 | 0 | | | | 0 |
| FA4/FA3B | 0 | | | | 0 | 0 | 0 | 0 | | | | 0 | | | | 0 | 0 | 0 | 0 | 0 | | | | | 0 | 1 | | | | 0 |
| Unoccupied | 5 | 7 | 39 | 0 | 0 | 22 | 0 | 0 | 0 | 0 | 0 | 0 | 31 | 7 | 20 | 0 | 24 | 0 | 0 | 0 | 0 | 2 | 2 | 2 | 0 | 0 | 8 | 0 | 22 | 0 |

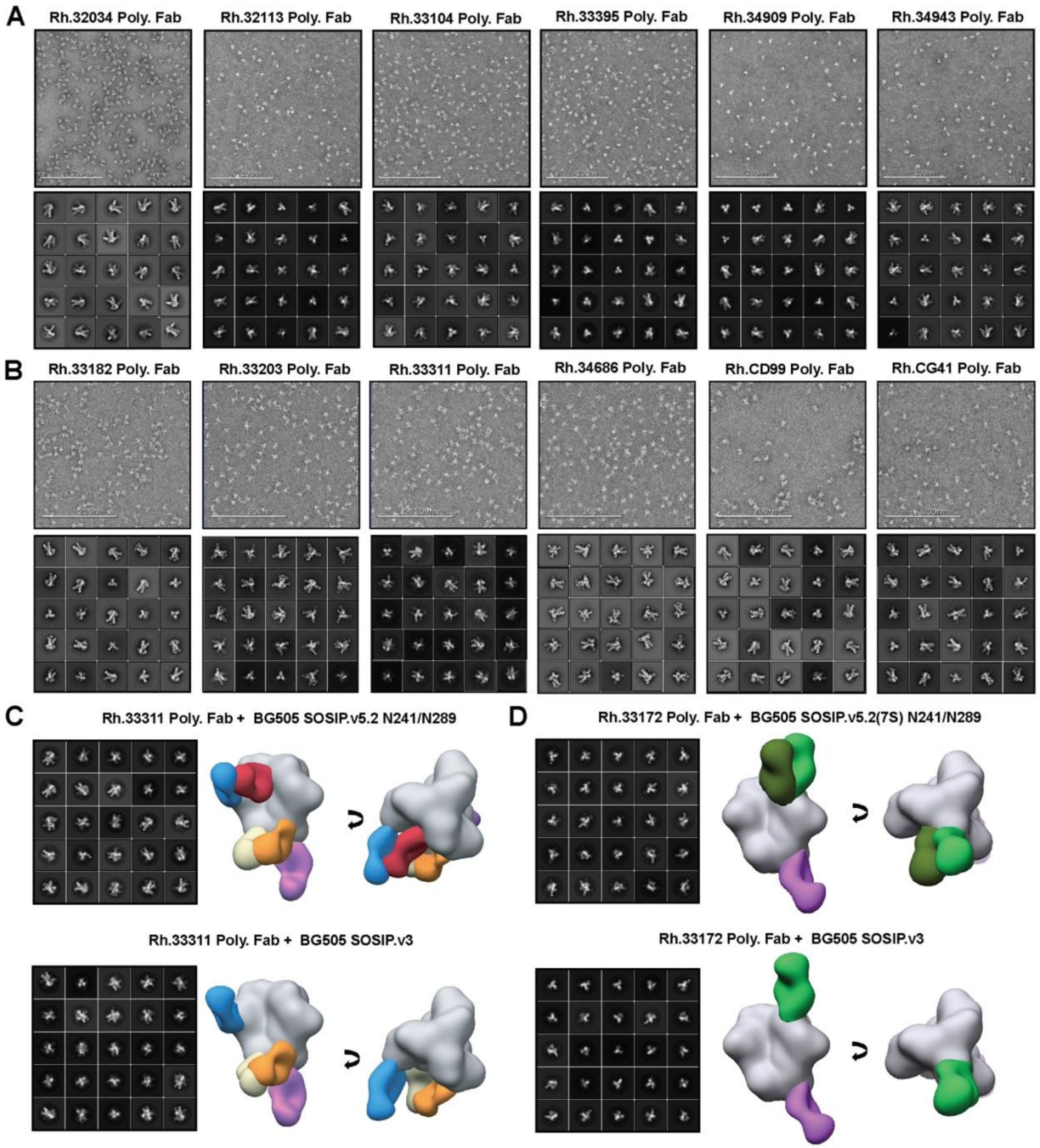


**Figure S1. Extended nsEMPEM data. [A,B]** Representative raw EM micrographs (top) and 2D class averages (bottom) from the nsEMPEM datasets used for the generation of Grp 1 and Grp 2 composite figures presented in Figure 1d. Immune complexes were generated with BG505 SOSIP MD39 for Grp 1 and BG505 SOSIP.v5.2 N241/N289 for Grp 2. **[C]** An equivalent amount of polyclonal Fab sample from animal Rh.33311 (wk 26) was complexed with BG505 SOSIP.v5.2 N241/N289 and BG505 SOSIP.v3 and imaged using EM to investigate the effect engineered stabilizing mutations have on antibody binding. Raw EM micrographs, 2D class averages and composite figures for the two nsEMPEM experiments are shown. **[D]** Similar experiment as in panel **(c)** was performed with Rh.33172 polyclonal Fab (wk 26). BG505 SOSIP.v5.2(7S) N241/N289 and BG505 SOSIP.v3 antigens were used for complexing. For color scheme, see the legend in Figure 1. In panel (c) two different shades of green (olive and bright green) were used to depict different polyclonal antibody classes targeting partially overlapping V1/V2/V3 epitopes.

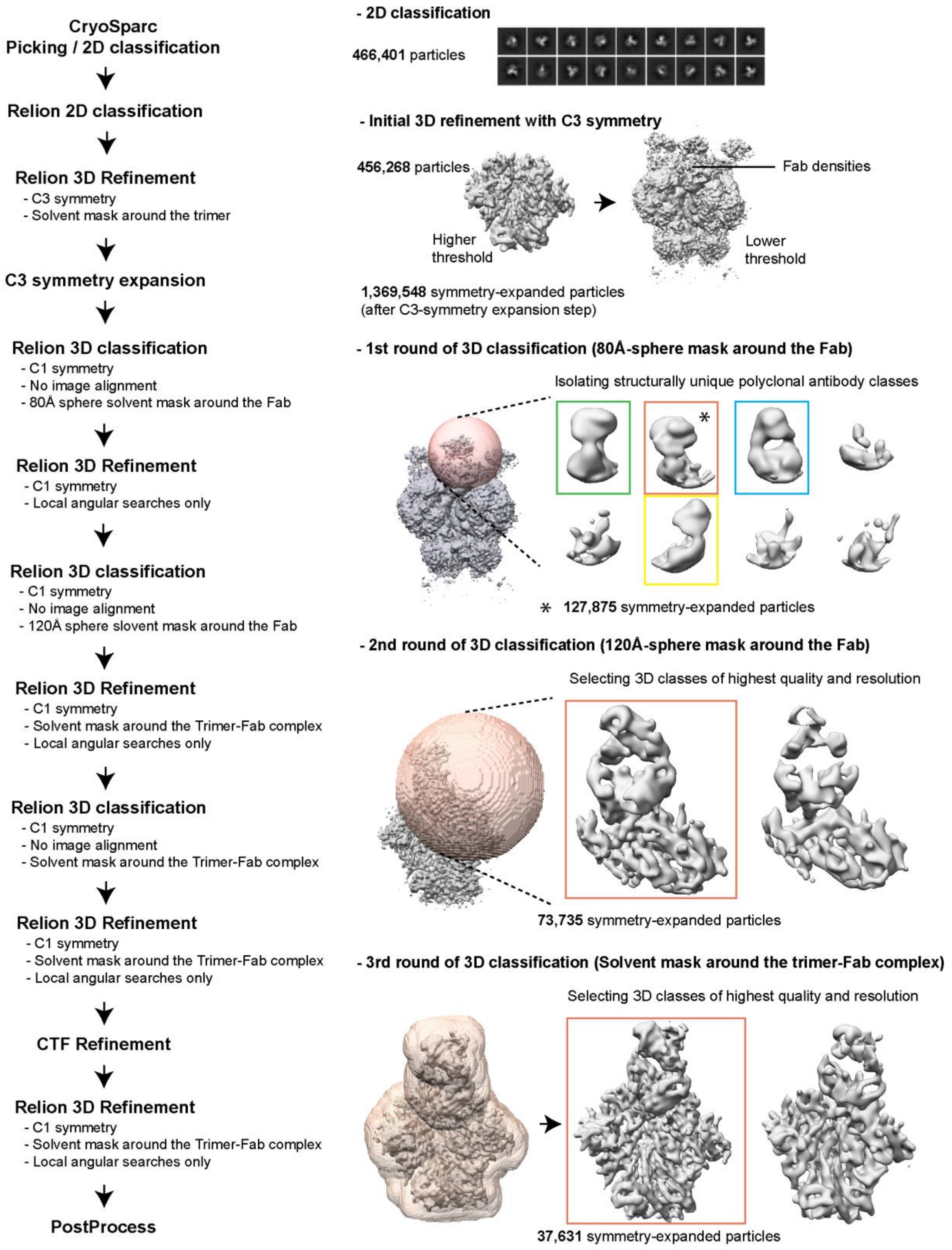


**Figure S2.** Schematic representation of the focused classification approach used for processing of cryoEMPEM data. Full data processing workflow is shown on the left and the examples of intermediate results from the Rh.33172 dataset are shown on the right.

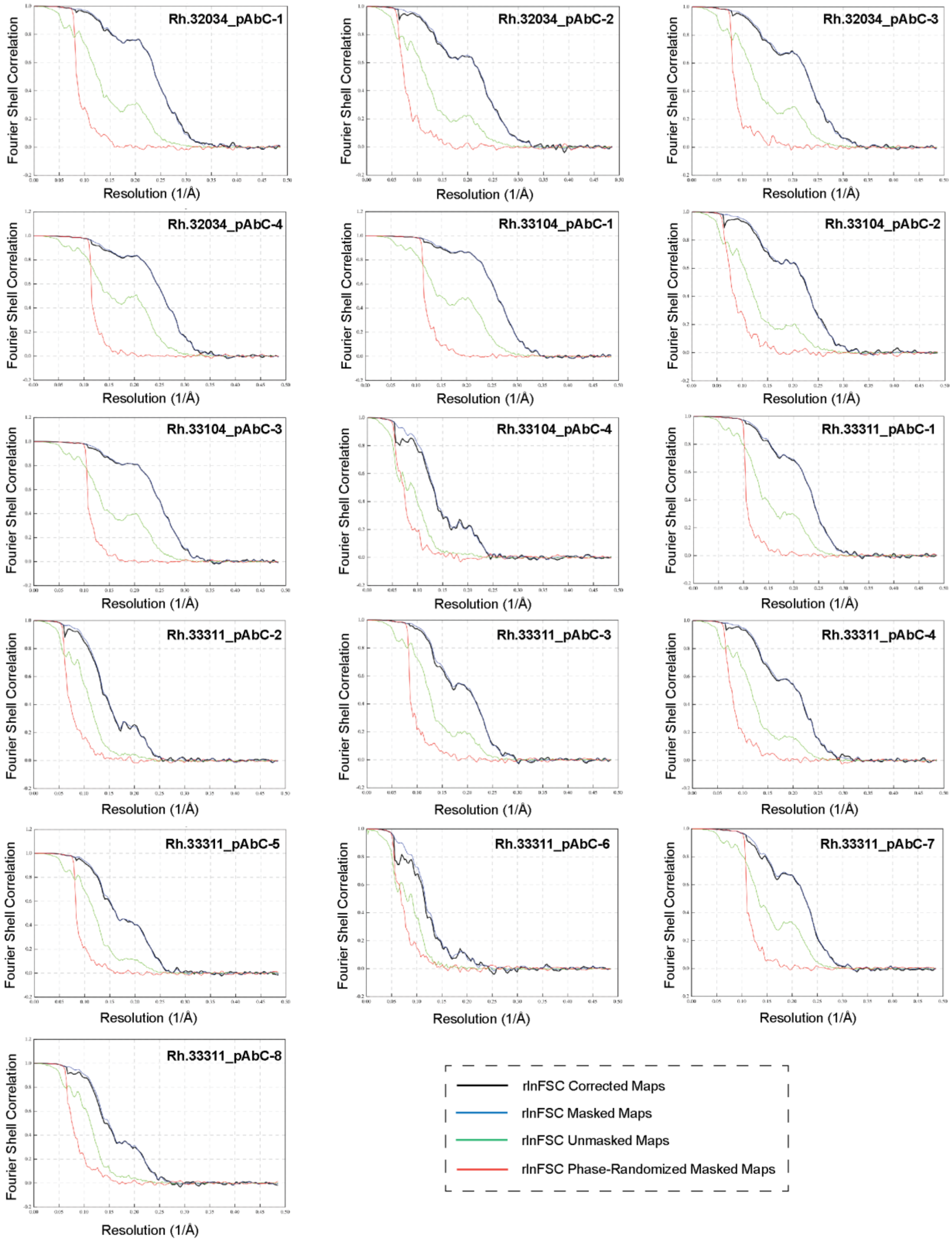


**Figure S3.** FSC resolution plots for EM maps reconstructed by cryoEMPEM analysis of polyclonal Fab samples isolated from animals Rh.32034, Rh.33104 and Rh.33311. The plots were generated in Relion/3.0.

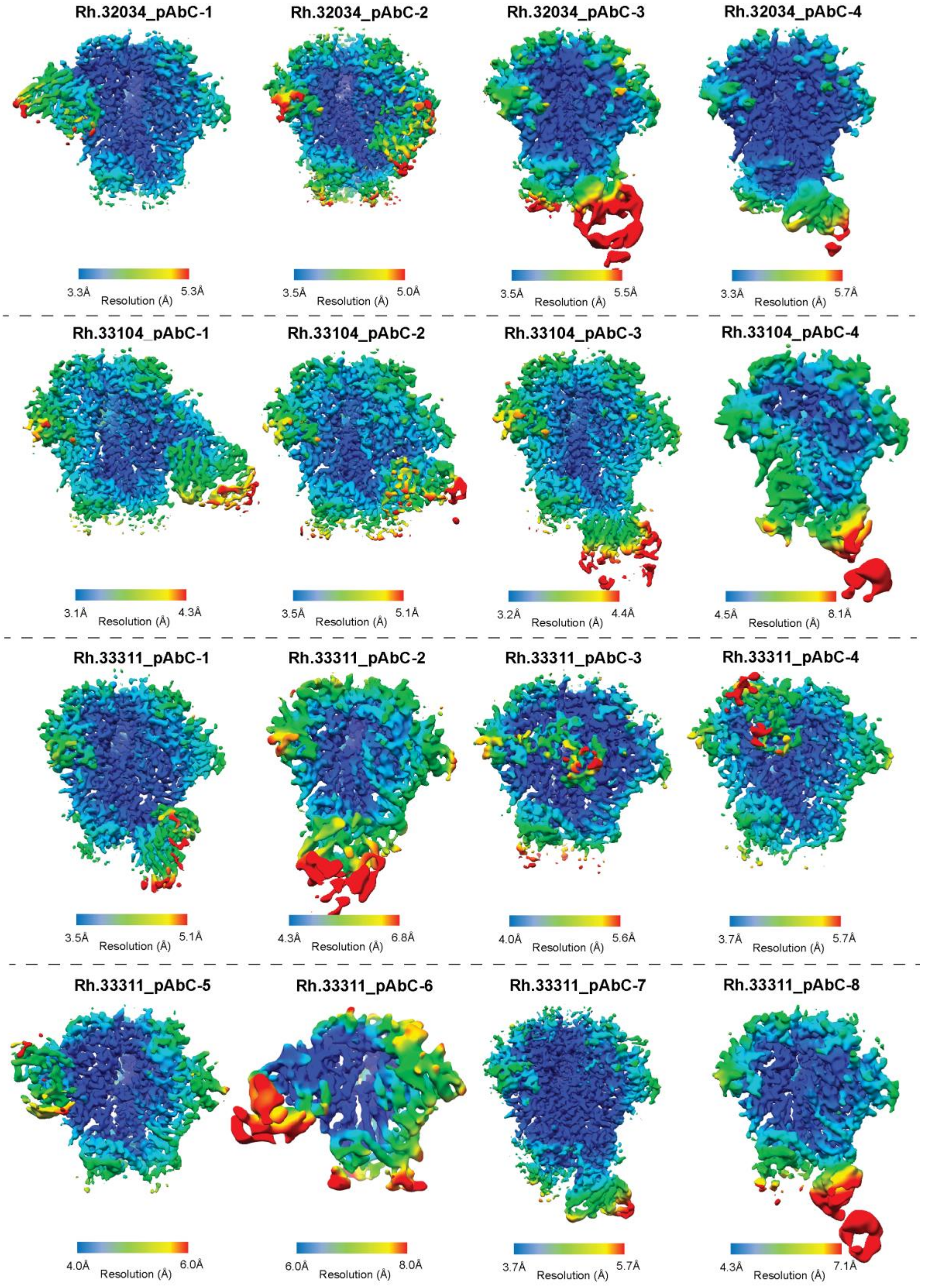


**Figure S4.** Local resolution plots for EM maps reconstructed by cryoEMPEM analysis of polyclonal Fab samples isolated from animals Rh.32034, Rh.33104 and Rh.33311. The plots were generated in Relion/3.0.

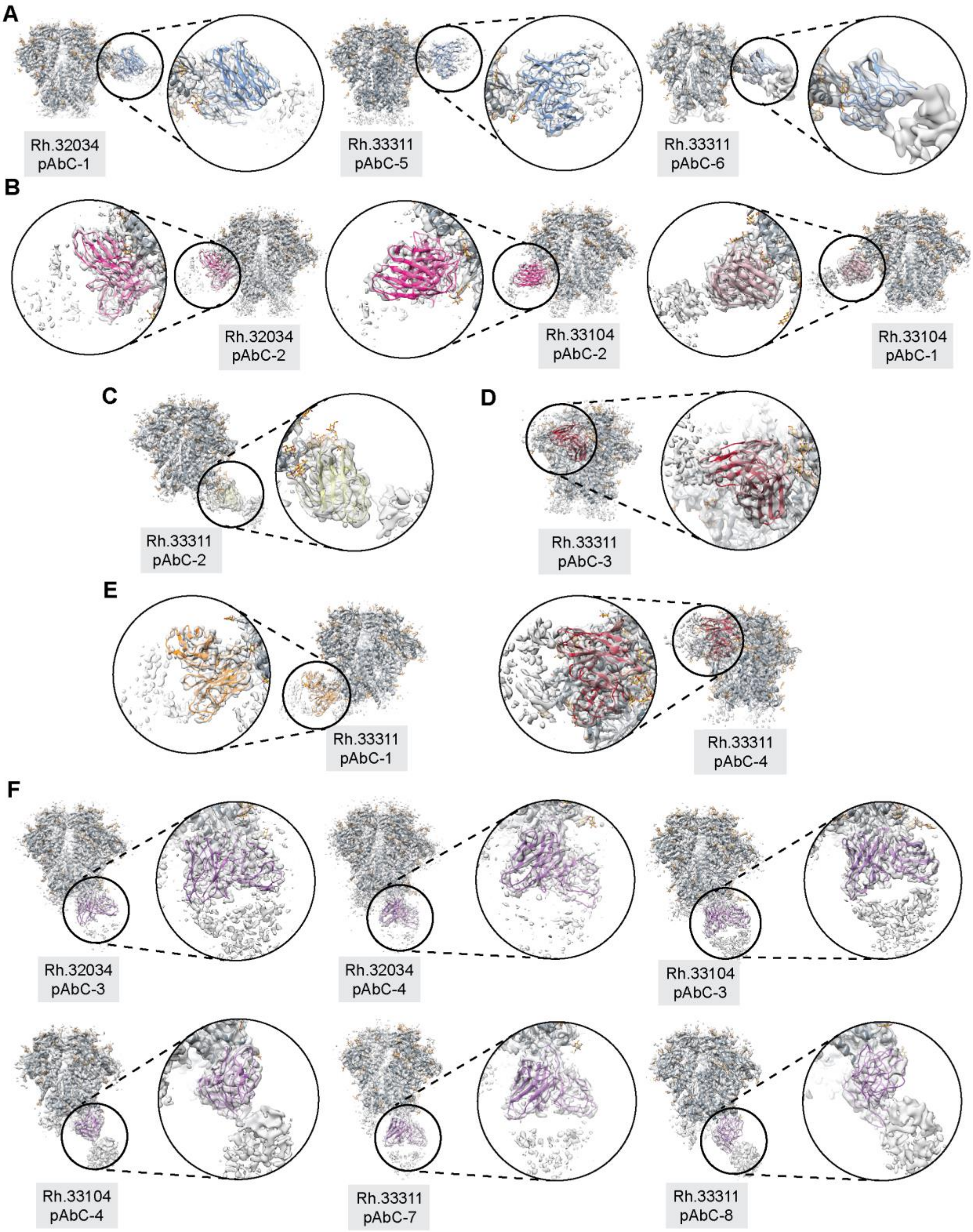


**Figure S5.** Models-to-map fit of BG505 SOSIP antigens in complex with reconstructed antibodies targeting V5/C3 epitope **[A]**, N241/N289 glycan hole **[B]**, N611 glycan epitope **[C]** fusion peptide **[D]**, gp120-gp120 interface **[E]** and base of the trimer **[F]**. A close-up view of the antibody backbone model (poly-Alanine) in the corresponding density is shown.

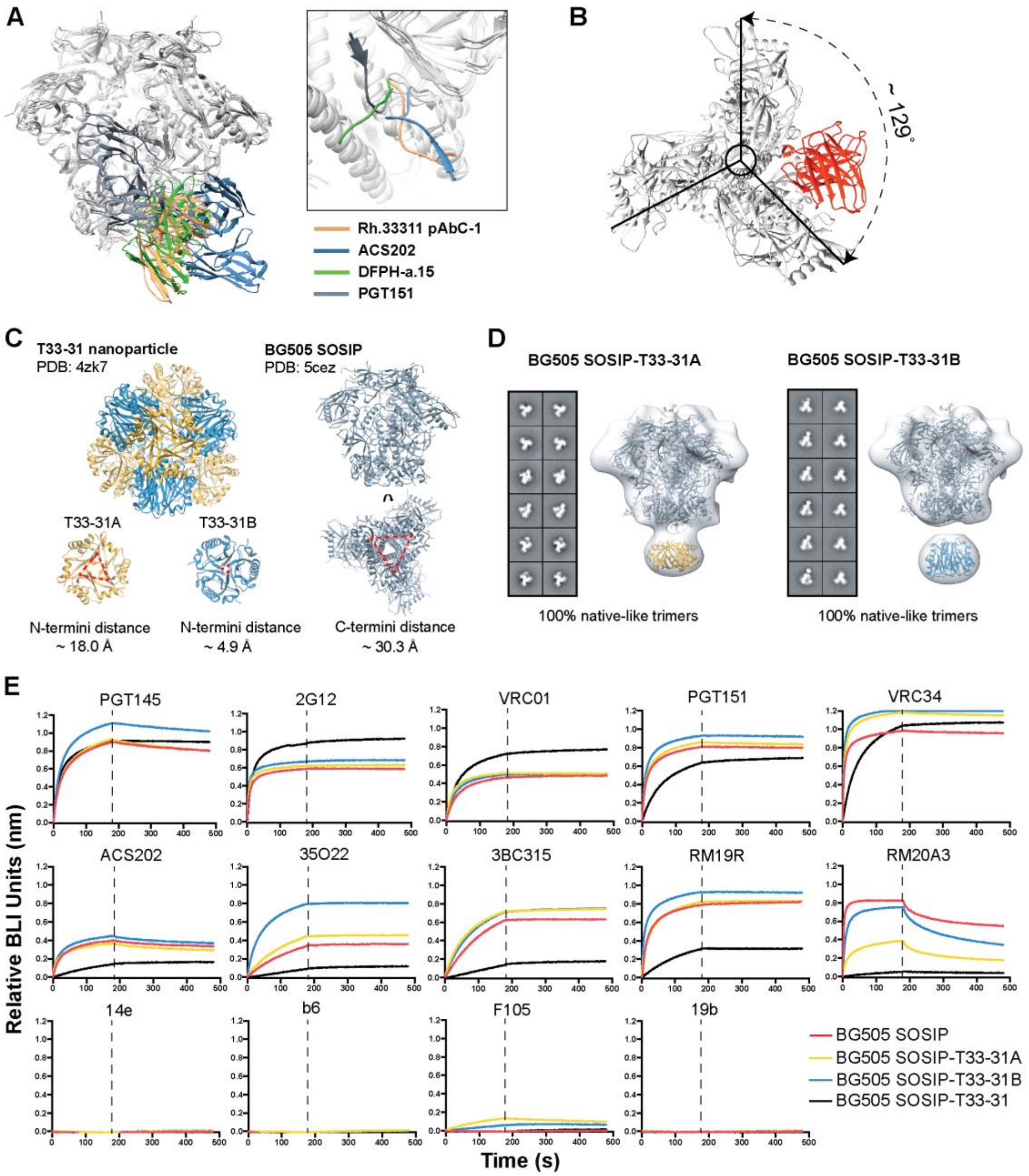


**Figure S6. [A]** The alignment of structures of FP-targeting bNAbs and Rh.33311 pAbC-1. The figure on the left shows full Fab complex structures. FP conformation in the antibody-bound state is shown in the panel on the right. **[B]** A view down the 3-fold symmetry axis of the BG505 SOSIP trimer in the Rh.33311 pAbC-4 complex structure showing the splaying of the trimer upon antibody binding. **[C]** The structure of the T33-31 two-component nanoparticle is shown in the top left corner and the BG505 SOSIP structure is shown on the right (ribbon representation is used; component A is colored yellow; component B is colored blue; BG505 SOSIP is colored gray). The distances between corresponding fusion residues in each trimeric building block (N-termini for T33-31A and T33-31B and C-termini for BG505 SOSIP) are shown in the bottom panels and these residues are connected by red dashed lines. **[D]** nsEM characterization of the two nanoparticle components fused to C-termini of BG505 SOSIP trimers. 2D-class averages are shown on the left in each panel; reconstructed EM maps are displayed on the right as transparent white mesh with BG505 SOSIP and nanoparticle components docked into the corresponding parts of the map. **[E]** BLI-based antigenicity analysis of BG505-SOSIP.v5.2(7S) N241/N289 antigens as free trimers, fused to T33-31A and T33-31B nanoparticle building blocks and assembled into a T33-31 nanoparticle. Antibody name is indicated above the corresponding panel.

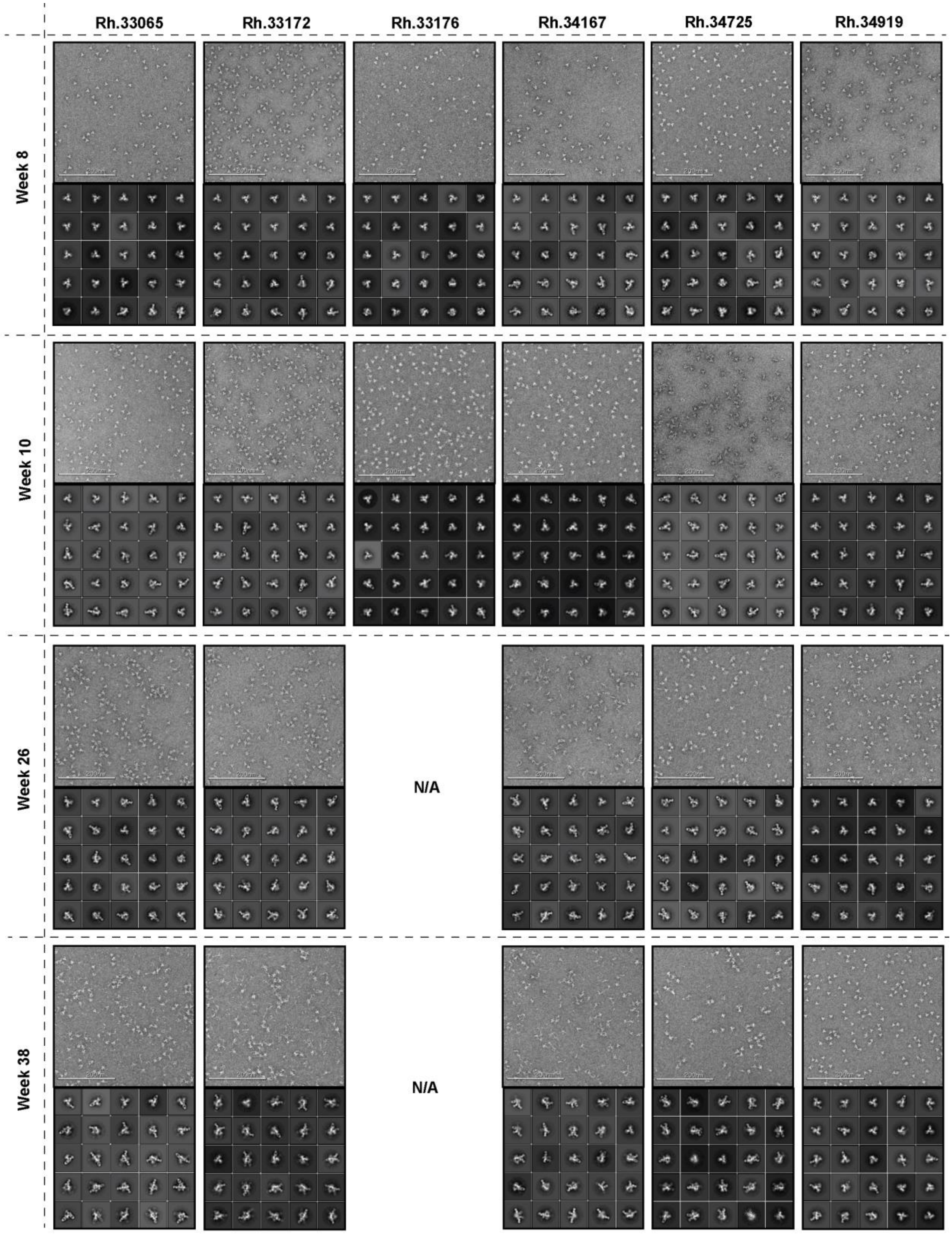


**Figure S7. Extended nsEMPEM data.** Representative raw EM micrographs (top) and 2D class averages (bottom) from the nsEMPEM datasets used for the generation of Grp 3 composite figures presented in Figure 6d. Animal ID is shown on top and the corresponding time points at which plasma samples were extracted are shown on the left.

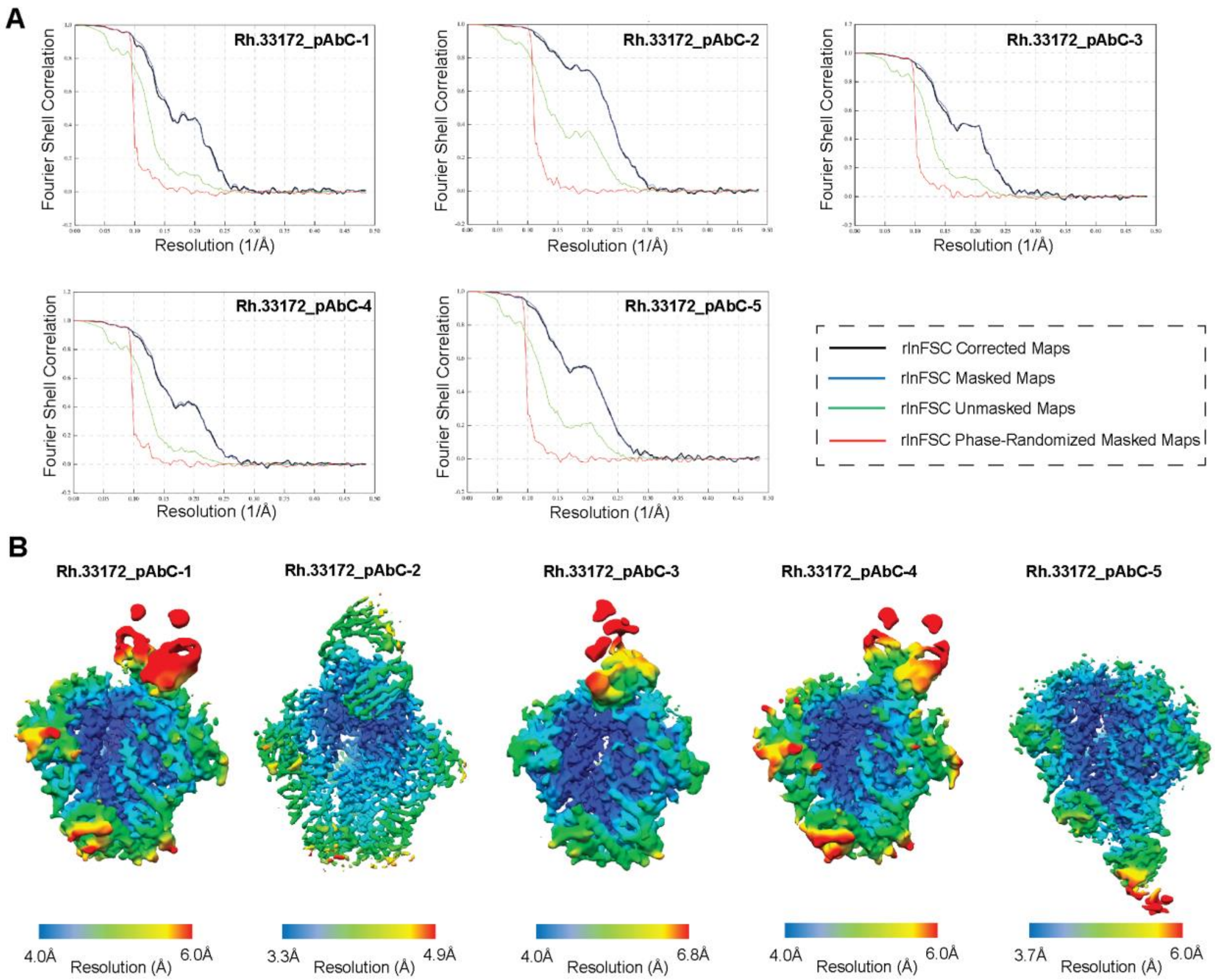


**Figure S8.** FSC resolution plots **[A]** and local resolution plots **[B]** for EM maps reconstructed by cryoEMPEM analysis of polyclonal Fab sample isolated from animal Rh.33172. The plots were generated in Relion/3.0.
